# High resolution analysis of proteolytic substrate processing

**DOI:** 10.1101/2022.03.24.485615

**Authors:** Jasmin Schillinger, Michelle Koci, Kenny Bravo-Rodriguez, Geronimo Heilmann, Farnusch Kaschani, Markus Kaiser, Christine Beuck, Hartmut Luecke, Robert Huber, Doris Hellerschmied, Steven G. Burston, Michael Ehrmann

## Abstract

Proteolysis is a key catalytic event in protein and thus cellular homeostasis. Despite the importance and wide implications of proteolytic processing and degradation, methods describing the degradation of folded proteins at high temporal and spatial resolution are not well established. However, this information is required to obtain a deep mechanistic understanding of proteolytic events and their consequences. Here, we describe an integrated method comprising time-resolved mass spectrometry, circular dichroism spectroscopy and bioinformatics to reveal the sequential degradation and unfolding of the model substrate annexin A1 by the human serine protease HTRA1. This workflow represents a general strategy for obtaining precise molecular insights into protease-substrate interactions that can be conveniently adapted to studying other posttranslational modifications such as phosphorylation in dynamic protein complexes.

---

Regulated proteolysis represents one type of protein-protein interaction that is essential to the fine-tuned regulation of cellular processes. To date, various methods characterize the proteolysis of substrate proteins including zymography, N-terminal sequencing, NMR spectroscopy or mass spectrometry (MS)^1–4^. However, current approaches provide only a limited understanding of the successive proteolytic events at high spatial and temporal resolution.

To more comprehensively understand substrate degradation, we employed the human endoprotease HTRA1. Members of the highly conserved high temperature requirement A (HtrA) family of serine proteases are implicated in protein quality control and cellular stress signaling^5–7^. Therefore, deregulation of human HTRA1 is associated with severe pathologies such as age-related macular degeneration, Alzheimer’s disease, cancer, arthritis, and familial ischemic cerebral small-vessel disease^5–7^. HTRA1 is a homotrimer in which each protomer is composed of an N-terminus corresponding to a partial insulin-like growth factor binding protein-7 domain of unknown function, a S1 serine protease domain and a C-terminal PDZ domain. High resolution structures of each of these domains were solved independently^8–10^ and a low resolution model of the complete protein has been proposed^11^. The substrate binding site of HTRA1 is located in a groove^9^. This architecture accommodates individual peptide chains at the active site resulting in limited sequence but rather conformational selectivity (Fig. 1A).

**Fig. 1.**
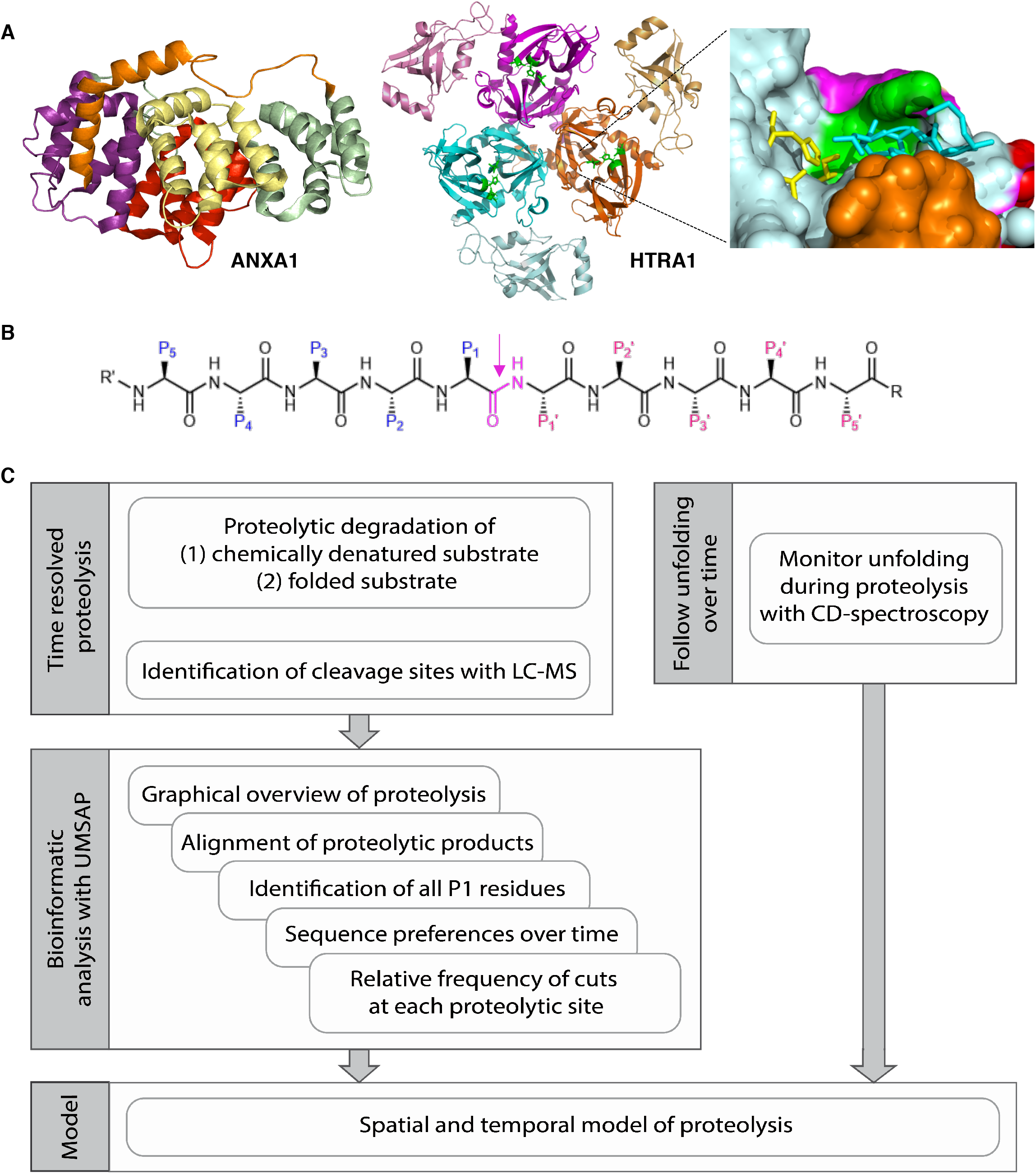
Proteolytic degradation of ANXA1 by HTRA1. A. Structures of ANXA1 (1HM6) and HTRA1 (3NZI). ANXA1 consists of an N-terminal domain of 41 residues (orange) and a core domain consisting of 4 helical repeats of about 75 amino acid residues each (purple, yellow, red and green). Model of the HTRA1 trimer. The protease domain was taken from 3NZI, the PDZ domains were modelled. The catalytic triad is shown in green. Right, close up of one active site with bound DPMFKLV-boro (stick representation, cyan), catalytic triad (H220, D250, S328; stick representation, yellow) and the activation domain i.e. loops L1 (green), L2 (red), L3 (orange) and LD (magenta). B. Model peptide and standard nomenclature. The residue of the scissile bond (magenta) is termed P1, which is often the major determinant of substrate specificity. Residues upstream of P1 are termed P2, P3 etc. Accordingly, residues located downstream to P1 are termed P1’, P2’, P3’ etc^28^. The proteolytic cleavage site is marked by an arrow. C. Workflow, see text for details.

A recent proteomics study identified annexin A1 (ANXA1) as a novel substrate of HTRA1^12^. ANXA1 is a 39 kDa protein composed of 21 helices and connecting loops (Fig. 1A). The crystal structures of ANXA1 revealed that its N-terminal domain is composed of 41 residues forming a helix and a loop, with the former inserting into the core domain. This core domain consists of 4 helical repeats of about 75 amino acid residues each^13,14^. In general, the annexin family of proteins is implicated in membrane biology including the repair of damaged cytoplasmic membranes^15^. While several proteases perform N-terminal processing of ANXA1^16^, our data suggest that it can also be completely degraded by HTRA1^12^. ANXA1 and HTRA1 were therefore chosen as a model to establish a general workflow to study the degradation of a folded substrate at high temporal and spatial resolution (Fig. 1C). The key elements of this workflow are the MS-based identification of proteolytic products and subsequent bioinformatic characterization of cleavage sites over a wide range of timepoints. Data analysis and representation is supported by UMSAP^17^ software for the analysis of MS data. In the context of targeted proteolysis, UMSAP uses a homogeneity of regression slopes test to identify relevant proteolytic products, the sequences of which are aligned along the primary amino acid sequence of the substrate, calculates relative frequency of cuts at each proteolytic site, as well as the amino acid sequence preferences of the protease^17^. In combination with circular dichroism spectroscopy (CD) spectroscopy, our method leads to a detailed understanding into the sequential events of initial substrate processing, concomitant unfolding and ultimate degradation into short peptides.

## RESULTS

### Proteolysis of chemically denatured substrate

The generation of a temporally and spatially resolved model of proteolysis of folded ANXA1 requires a reference for the identification of all potential cleavage sites and to distinguish those that are cleaved efficiently, poorly or not at all, irrespective of their accessibility in the folded substrate. Therefore, ANXA1 was chemically denatured in 8 M urea. Subsequently, substrate was diluted 13 fold into buffer and incubated with stoichiometric amounts of HTRA1 at 37°C. Samples were taken at eight timepoints ranging from 0 to up to 600 sec. As controls, denatured substrate and protease were directly added to acetone. Four biological replicates of each sample were subjected to LC-MS analyses to identify proteolytic products and the cleavage sites. The resulting data sets were analyzed using UMSAP software^17^. For an initial graphic overview, UMSAP groups proteolytic products into fragments and provides the total number of cleavage sites at each timepoint (Fig. 2). These data revealed 31 cleavage sites after 15 sec and 167 cleavage sites after 600 sec of incubation. The identified peptides span almost the entire sequence after 30 sec with the exception of short regions comprising residues 65-81, 146-160, 255-263 and 319-324. These gaps were not observed from the 90 sec timepoint onwards, indicating that the degradation of substrate was nearing completion (Fig. 2, Supplementary data 1).

**Fig. 2.**
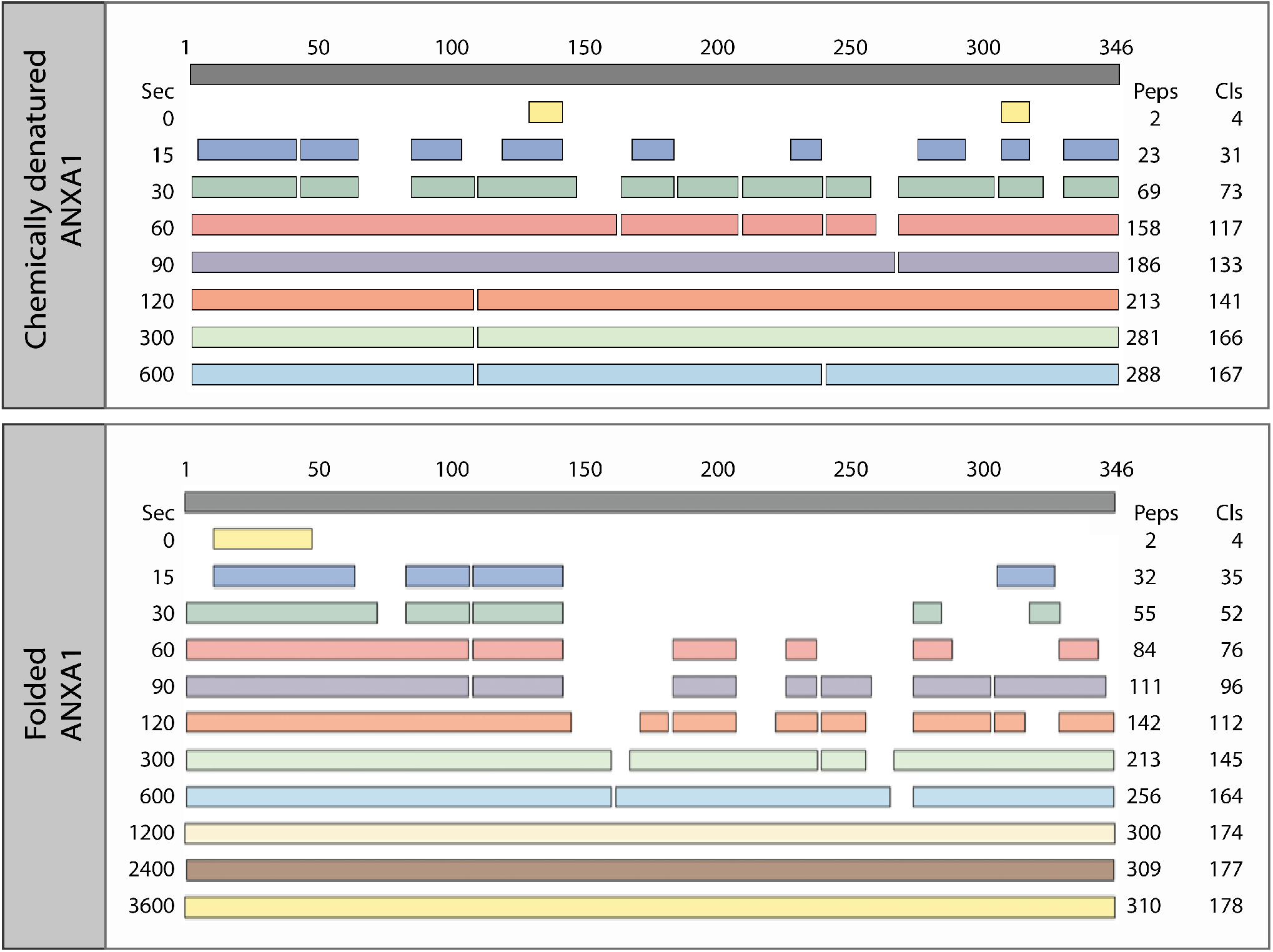
Proteolysis of chemically denatured and folded ANXA1. 20 μM chemically denatured and folded ANXA1 were incubated with 20 μM HTRA1 at 37°C. Samples were taken at the time points indicated and proteolytic products of ANXA1 were identified by LC-MS. Upper panesl, Top bar: linear representation of the entire substrate protein. For each of the 0 - 600 sec for denatured and 3600 sec for folded ANXA1 time points (Sec), respectively, peptide sequences that align without gaps are grouped into so-called fragments shown as bars. The number of peptides identified (Peps) and the total number of cleavage sites (Cls) are given at the right. The identified peptides aligned to the primary amino acid sequence of ANXA1 are provided in Supplementary data 1 (chemically denatured ANXA1) and 3 (Folded ANXA1).

Each proteolytic product is the result of two cuts, one at the N-terminus, where the N-terminal residue represents the P1’ residue, and another at the C-terminus, representing the P1 residue (Fig. 1B). UMSAP also calculates the relative frequency of cuts at each P1 residue for each timepoint (see Methods for details). These data provide quantitative information on how well HTRA1 cleaves the substrate at each position within its primary amino acid sequence as well as how proteolysis progresses over time (Fig. 3). In addition, based on histograms (SI Fig. 1), individual P1 residues can be grouped into several classes. The first class comprised 17 P1 residues where the relative frequency of cuts reached ≥20 (Table 1). Of these, L181, M276, L282, M300, Q316, C324 and A326 are buried in the folded protein. Their efficient processing indicates that chemical denaturation did occur. The second class comprised 25 P1 residues where the relative frequency of cuts lies between 11 and 19 (Fig. 3, SI Table 1). The third class comprised residues where the relative frequency of cuts was between 1-10. These were considered poor sites at all timepoints, probably resulting from a low affinity to the active site. The fourth class comprised residues for which no peptidic products could be detected. This lack of detection might be explained by various models: i) these sites did not bind to the active site, ii) the resulting products were not detectable by MS or iii) upon dilution of urea, this part of the substrate refolds into a stable conformation where the cleavage site remains inaccessible to the active site of HTRA1.

**Fig. 3.**
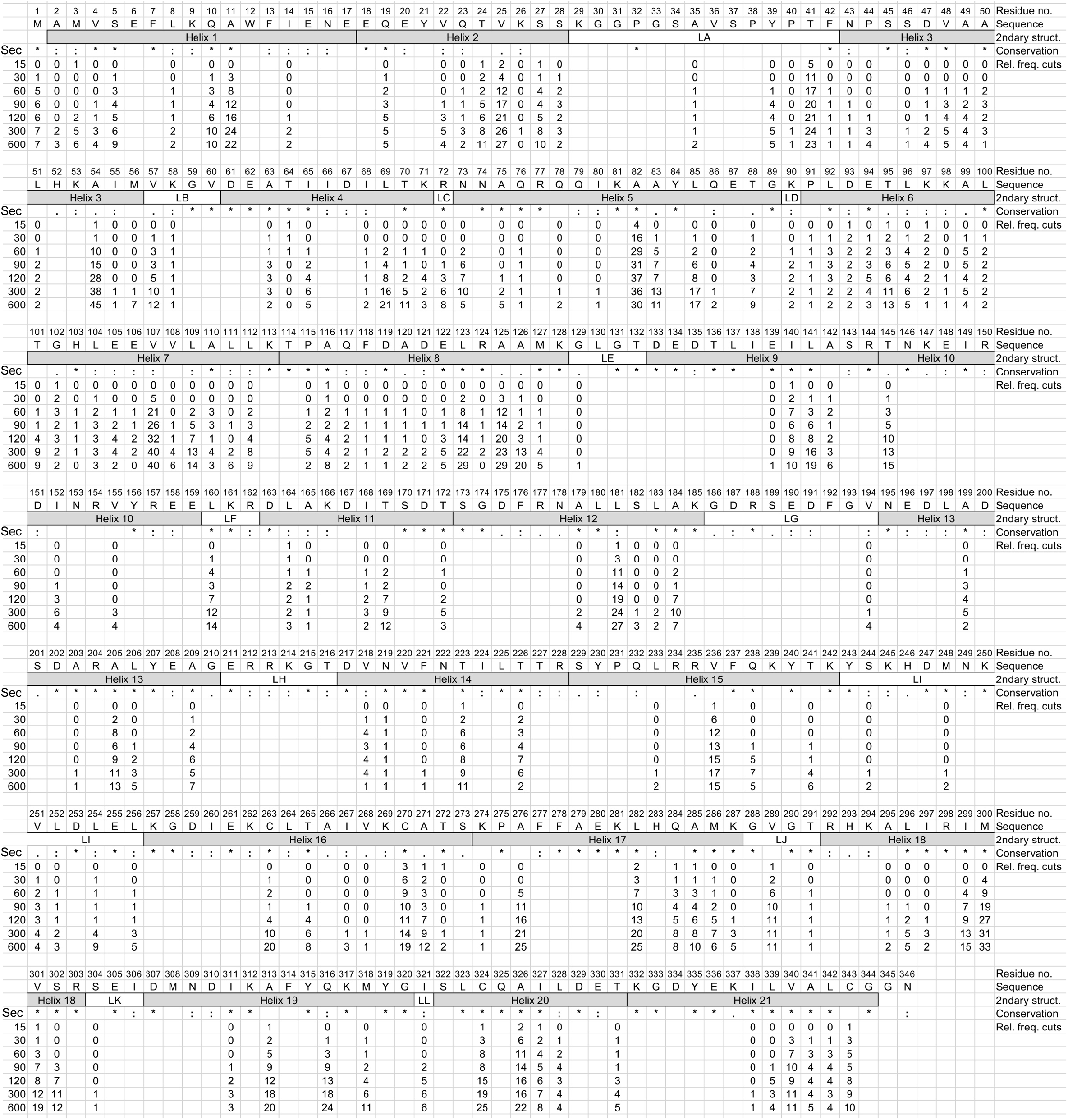
Proteolysis of chemically denatured ANXA1. Chemically denatured ANXA1 was incubated with HTRA1 at 37°C. Samples were taken at the time points indicated (sec) and proteolytic products of ANXA1 were identified by LC-MS. Structural elements i.e. helices and loops (2ndary struct.) as detected in the crystal structure of folded ANXA1 (pdb:1HM6) are indicated. Amino acid sequence conservation (Conservation) is derived from a multiple sequence alignment (Supplementary data 2). * = identical,: = conserved residues. All P1 residues identified in n=4 independent experiments that were significantly enriched compared to controls are represented by numbers. Numbers below individual P1 positions indicate the relative frequency of cuts (Rel. freq. cuts) at each time-point. No number indicates that no cleavage was detected at any of the time points investigated.

**Table 1.**
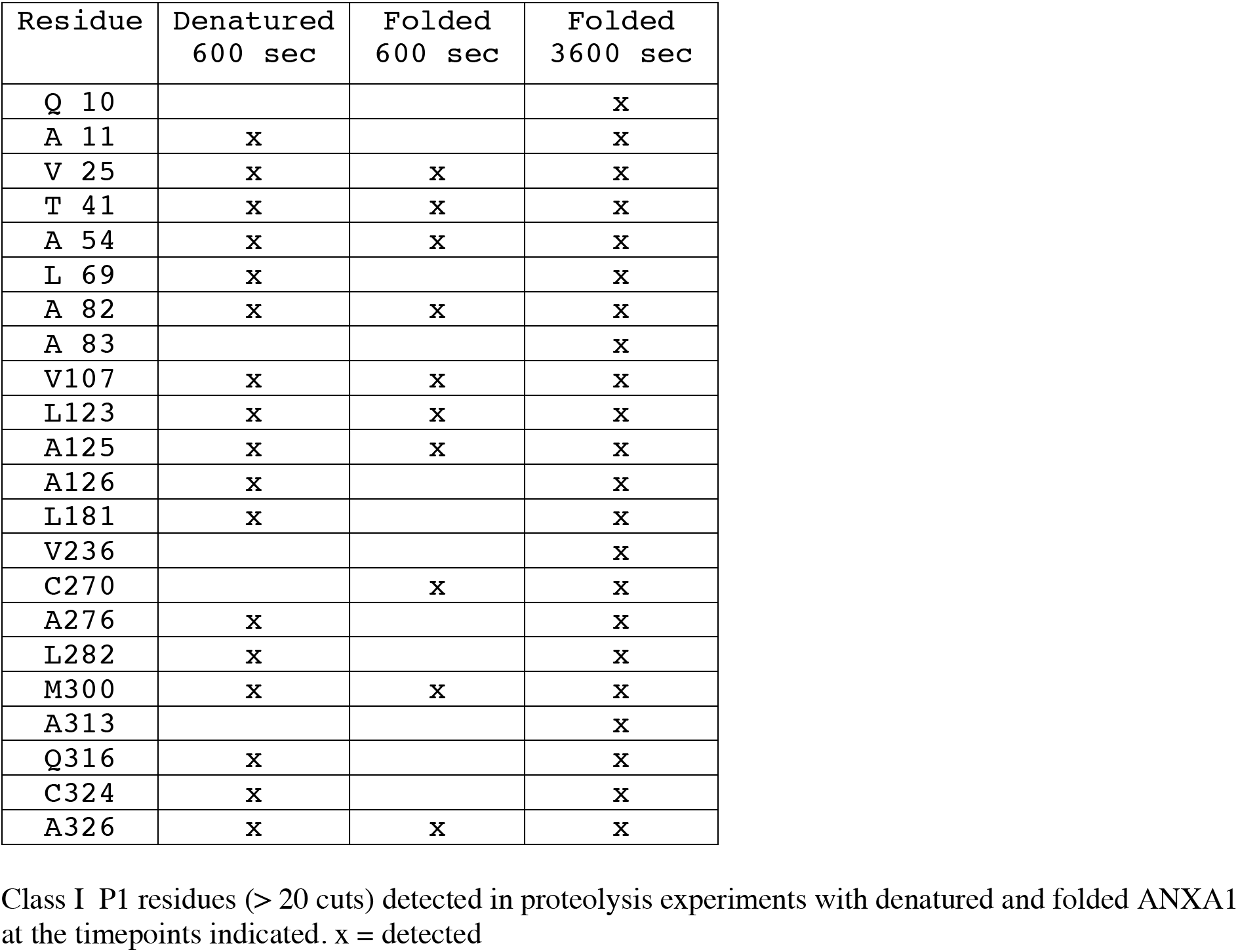
List of P1 residues with the highest relative numbers of cuts.

### Proteolysis of folded substrate

The analysis of degradation of folded substrate provides information on surface accessibility of substrate binding sites and the degrees of sequential structural relaxation and unfolding that occur during progressive degradation. Therefore, ANXA1 was incubated with HTRA1 at 37°C and samples were taken at eleven timepoints ranging from 0 sec to up to 3600 sec. The time of incubation was extended because proteolysis of folded substrate was slower compared to chemically denatured substrate. These data revealed 35 cleavage sites after 15 sec, 164 cleavage sites after 600 sec and 178 cleavage sites after 3600 sec of incubation. In contrast to the degradation of denatured ANXA1, the identified peptides did not span the entire sequence up to the 600 sec timepoints. Complete coverage was only observed from the 1200 sec timepoint onwards (Fig. 2, Supplementary data 3). When grouping P1 sites into classes according to the maximal relative number of cuts performed at each site, class 1 (≥20 cuts) comprised 20 residues (Table 1) and class 2 (11-19 cuts) comprised 31 residues (Fig. 4, SI Table 1).

**Fig. 4.**
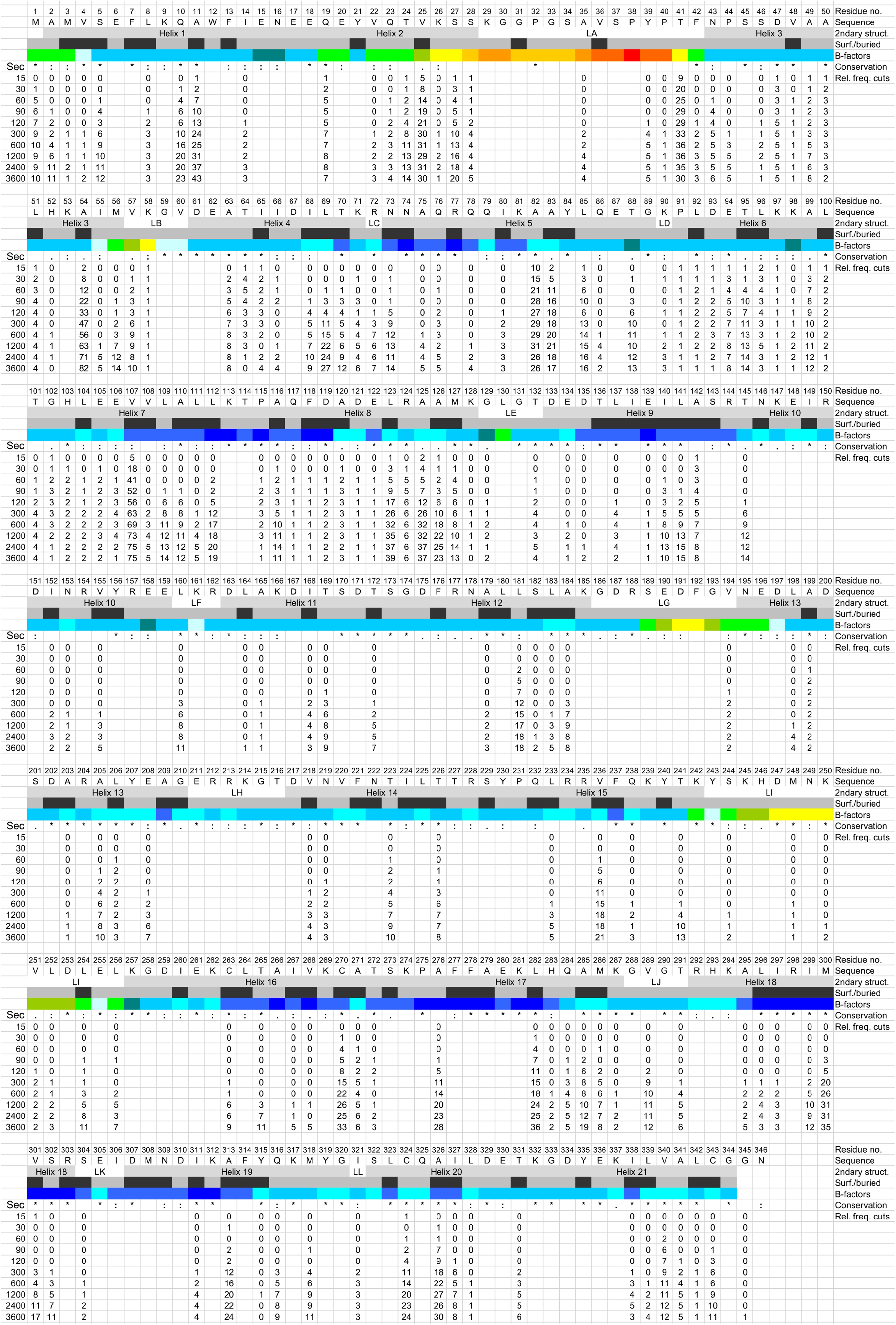
Proteolysis of folded ANXA1. Proteolysis of folded ANXA1 was done as described in Fig. 3. P1 residues identified in n=4 experiments as significantly enriched compared to controls are represented by numbers. Structural elements i.e. helices and loops (2ndary struct.), B-factors and surface accessibility (surf. (grey)/buried (black) as detected in the crystal structure of folded ANXA1 (pdb:1HM6) are indicated. The colour code for B-factors indicates gradually structural rigidity (blue) to flexibility (red). Amino acid sequence conservation (Conservation) is derived from a multiple sequence alignment (Supplementary data 2). * = identical,: = conserved residues. Numbers below individual P1 positions indicate the relative frequency of cuts (Rel. freq. cuts) at each time point. No number indicates that no cleavage was detected at any of the time points investigated.

In contrast to chemically denatured substrate, the initial 35 cleavage sites at the 15 sec timepoint were, with the exceptions of S302 and C324, exclusively located in the N-terminal region of ANXA1 up to residue A142, comprising helices 1-9 (Fig. 4). At this timepoint, the P1 sites exhibiting the highest relative frequencies of cuts are A82 in helix 5, T41 in loop LA, V107 in helix 7 and V25 in helix 2 (Fig. 4). The 178 cleavage sites identified across the entire ANXA1 sequence at 3600 sec timepoint correlates with the 167 cuts at the 600 sec timepoint of denatured substrate indicating that the extend of proteolysis of denatured and folded substrates was similar at these timepoints. Here, the P1 sites exhibiting the highest relative frequencies of cuts are A54 at the C-terminus of helix 3, V107 in helix 7, A11 in helix 1, L123 and A125 in helix 8 as well as L282 in helix 17.

These data suggest that early proteolytic events may destabilize the substrate’s folded structure causing accessibility of sites that were not surface exposed. Interestingly, the tendency of more potent processing of the 9 N-terminal helices was also observed with chemically denatured protein (Fig. 2), suggesting that the C-terminal part of ANXA1 might have evolved to be more protease resistant at least towards HTRA1. The latter notion is supported by the larger gaps within the C-terminal half of ANXA1 (Fig. 2 - 4) as well as by the increased number of P1 sites that are cleaved more efficiently in denatured versus folded ANXA1 (SI Table 2).

In addition, we observed three features for folded and denatured substrate. The appearance of many products sharing only one identical cleavage site suggests that these products are the result of one high affinity and various low affinity binding sites (Supplementary data 1, 3, 4). This model is supported by the observed tendency of longer proteolytic fragments, again sharing one identical cleavage site, arising with higher frequency at later timepoints suggesting that these additional cleavage sites are of even lower affinity. Moreover, sequences corresponding to central parts of loops LA, LG, LH and LI appear to be less well cleaved compared to helices. This might be caused by their amino acid composition resulting in conformations that are inaccessible to the active site.

### Comparison of the amino acid sequences surrounding the cleavage sites

When comparing the amino acid sequences surrounding the cleavage sites (SI Fig. 2)^17^, the patterns of preferred residues at late timepoints (600 sec for denatured and 3600 sec for folded ANXA1) are very similar for the denatured and folded substrates e.g. at P1 sites, Ala (26.7 and 26.6%, respectively), Leu (16.8 and 16.4%), Val (14.0 and 13.6%) and Thr (11.9 and 12.2%) dominate, while Ala (13.5 and 13.5%) and Lys (12.0 and 11.5%) are enriched at P1’ sites. This feature is in agreement with published data^9^. This similarity is also observed for the residues P2-P5, P2’and P5’ (SI Fig. 2). In contrast, the preferred residues differ at early timepoints and converge over time. These observations were also made when examining consensus sequences (SI Fig. 3). Here, the various consensus sequences at each position were reached two to three timepoints earlier for denatured compared to folded ANXA1. These observations suggest that for the folded protein, the limited availability of the preferred small hydrophobic residues reflects conformational exclusion of these sites at early timepoints. This phenomenon is lost at later timepoints, probably because proteolytic digests cause progressive substrate unfolding.

### Initial events in degradation of folded ANXA1

Given the importance of initial cuts at early timepoints, we sought to increase the resolution of these events by lowering the number of initial cuts. We therefore reduced the concentration of protease 5, 10 and 20-fold, took samples at 0, 15 and 30 sec and identified the proteolytic products by LC-MS. At 1 μM HTRA1, a single peptidic product was identified after 15 sec of incubation, while 6 peptides were identified after 30 sec, resulting from 8 cuts within the N-terminal 125 residues (SI Fig. 4, Supplementary data 4). At 2 μM HTRA1, again a single cut was identified after 15 sec of incubation, while 13 peptides were identified after 30 sec, resulting from 17 cuts within the N-terminal 126 residues (SI Fig. 4, Supplementary data 4). At 4 μM HTRA1, 8 peptides were identified resulting from 12 cuts after 15 sec, while 26 peptides were identified resulting from 29 cuts after 30 sec of incubation. One additional product was identified located at the C-terminus, resulting from cuts at P1 residues A326 and V340. As this peptide was only detected at the 15 sec but not at the 30 sec timepoint and only from the 60 sec timepoint onwards at 20 *μ*M HTRA1 concentration (Supplementary data 3), this peptide was omitted in the model of sequential substrate unfolding described below.

The analysis of initial events must consider the P1 residues per se but also the peptidic products resulting from additional cuts occurring either up-or downstream of the P1 residue. For example, cuts at A82 result in 6 products at 2 *μ*M HTRA1 at 30 sec that are generated by additional cuts at E94, L96, K97, A99, G102, and V107. At 4 *μ*M HTRA1 and 30 sec cuts at A82 result in 9 products resulting from 3 additional cuts at D93, T95 and L100. Another example is residue T41. Here, additional cuts upstream, i.e. at V25, K26 and K29 as well as downstream at A54, K58 and T64 are producing 6 peptides. These events lead to the efficient degradation of the region comprising V25 to T64 with the understanding that the cut at T41 is the main event (Supplementary data 4).

### Model of sequential unfolding by progressive proteolysis

A model of sequential unfolding by proteolytic fragmentation can be postulated by considering the occurrence of cuts at low HTRA1 concentrations at early timepoints in combination with high relative frequency of cuts at individual P1 residues at higher HTRA1 concentrations. Taken together, our data suggest that the degradation of ANXA1 starts at its N-terminus causing fragmentation up to helix 8, followed by the degradation of the remaining C-terminal parts (Fig. 5A).

**Fig. 5.**
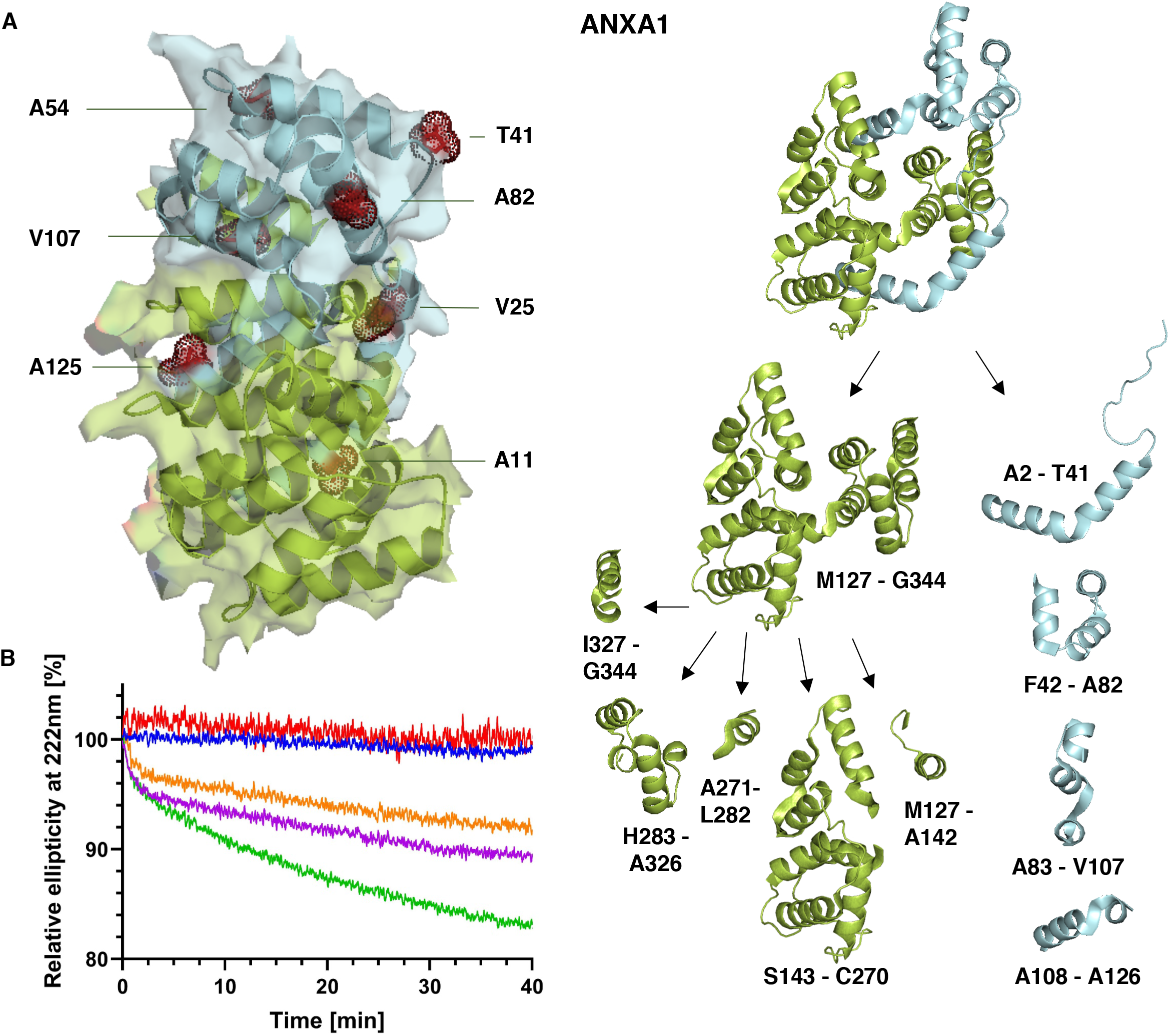
Early cleavage events in folded ANXA1 and unfolding by fragmentation. A. Left, surface representation of ANXA1 (side view). Key P1 residues are shown as sticks and dot representation. Surface exposed residues are shown in red, buried residues in orange. Right, model of how folded ANXA1 (top view) is first converted into 4 N-terminal fragments comprising residues A2-A126 (lime) generating a C-terminal fragment comprising M127-G344 (limon). Subsequently, the C-terminal fragment is further processed into 5 fragments. B. Circular dichroism spectroscopy. HTRA1 and ANXA1 were mixed in a 1:1 (*green*), 1:10 (*magenta*) and 1:20 (*orange*) ratio and the ellipticity at 222 nm monitored over time. To facilitate comparison between datasets, in each case the initial signal was set at 100% and the loss of the negative ellipticity at 222 nm is shown as a percentage of the overall signal. 5 *μ*M HTRA1 alone (*red*) or 5 *μ*M ANXA1 alone (*blue*) control experiments are also shown.

Residues T41, A82 and V107 are prime candidates for being cleaved early and at high frequency. Among these, the surface exposed and non-conserved T41 located at the fusion joint of the N-terminal and the core domain of ANXA1, as well as at the C-terminal end of the large loop LA is likely the first P1 residue to be cleaved (Fig. 4). Subsequently, the site at V25, located at the C-terminus of helix 2 immediately preceding loop LA, could be cleaved next. This cut is likely followed by cleavage at A11, an event that would require helix 1 (residues 2-17) to move away from the folded structure, which is likely to occur after the position of helix 2 (residues 8-28) had been destabilized by the cut at V25. In parallel, cuts after A82, located in the middle of helix 5, will cleave this helix in half. This event is likely to destabilize the positions of helices 3 and 4, allowing a cut at A54 near the end of helix 3. Concerning V107, this cut in the middle of helix7 should allow subsequent cuts at A125/A126 in helix 8 that are completing the removal of the N-terminal 8 helices from the folded structure (Fig. 5A).

The subsequent processing of the remaining C-terminal part of ANXA1 involves early cuts at A142, C270, L282 and A326 producing 5 fragments, i.e. M127-A142, S143-C270, A271-L282, H283-A326, I327-G344. Subsequent proteolytic events involve I140, L181, T223, V236, A313, C324 and V340. Ultimately, all fragments are further processed by multiple cuts. The regions in which no cuts were observed at the latest timepoints examined comprise no more than 9 residues i.e. K185-G193 and A209-D217, suggesting that unfolding and degradation of substrate is complete.

### CD spectroscopy during proteolysis

To independently follow substrate unfolding, CD spectroscopy was employed (Fig. 5B, SI Fig. 5). The CD spectra of ANXA1 and HTRA1 alone exhibited the classic double minimum (208 nm/222 nm) allowing us to monitor the *α*-helical content at 222 nm. Since ANXA1 is composed only of *α*-helices and HTRA1 predominantly of *β*-sheets, the signal of HTRA1 at 222 nm was significantly lower compared to that of ANXA1. When ANXA1 was mixed with HTRA1 and subjected to CD spectroscopy after 5 h digestion, a 75% reduction of the ANXA1 signal at 222 nm, but only a 50% signal reduction is observed at 208 nm, with the spectral minimum being shifted to the lower wavelength (SI Fig. 5). This indicates that proteolytic fragmentation is indeed accompanied by a loss of ANXA1 structure as indicated by the overall loss of signal intensity while the shift towards lower wavelengths indicates increased random coil content. The decrease of a-helical content of the folded structure in the digest reaction was followed continuously by measuring the CD signal at 222 nm over time (Fig. 5B). Notably, an initial exponential burst phase during the first 60 sec was observed, which correlates with the fast initial cuts caused mainly by the cuts at P1 residues T41, A82 and V107. This relatively rapid loss of signal indicates that the peptidic products resulting from cuts at these sites is accompanied by their rapid dissociation and unfolding, suggesting that both the tertiary contacts and the contiguity of the polypeptide chain is necessary to stabilize the secondary structure of the cleaved peptides within the intact protein. This fast initial phase is followed by a more slowly proceeding phase, indicating a likely scenario where the remainder of the undigested ANXA1 has sufficiently destabilized tertiary structure and core hydration that unfolding events take place. This results in a competition between refolding and proteolysis, which eventually leads to a progressive digestion of the whole ANXA1 at available sites along its linear sequence, although it is possible that some residual structure may remain and occlude sites leading to slower digestion. This is supported by the fact that as HTRA1 moves from stoichiometric to sub-stoichiometric concentrations, the rate of the gradual loss of the CD signal following the initial burst phase decreases (Fig. 5B).

## DISCUSSION

Our method combines time-resolved MS identification of peptidic products, the corresponding protease cleavage sites and the relative frequency of cuts at each specific site. Moreover, correlating the temporal occurrence and the location of the observed cuts in the folded substrate reveals the sequential degradation of the model substrate ANXA1 by the serine protease HTRA1 at unprecedented temporal and spatial resolution. These data provide detailed information about substrate degradation that does not require prior modification and ATP-driven unfolding and proteolysis as exemplified by the ubiquitin, p97, proteasome system^18^.

The advantages of the method include insights into how a folded substrate is degraded in sequential steps i.e. from initial processing followed by local structural relaxation to stepwise unfolding of the substrate and subsequent complete degradation of the initially produced large fragments into short peptidic products. We expect the method to be applicable to heterooligomeric complexes and other structures e.g. disease-related amyloid fibrils, the dissociation and degradation of which is not yet understood at high resolution^19–21^. The breath of data generated also provides the basis for further improvements of downstream computational processing e.g. the development of predictive algorithms concerning structural changes caused by proteolytic processing as well as for conformational and amino acid sequence selectivity of the proteases under investigation. In addition, our method provides structural insights i.e. information on surface accessibility of individual residues in particular, which would be useful for proteins for which high resolution structures are unavailable. Moreover, the generated data will allow to postulate experimentally addressable hypotheses concerning the structural and functional consequences of missense mutations identified in pathologic events such as cancer and other genetic diseases.

The main limitation of the method is due to MS constraints i.e. the inability to detect specific peptidic fragments in an unforeseeable manner. However, this issue is likely to soften given the continued development of new instruments with improved resolution and method development towards optimized sample preparations.

Our data lead to additional insights. Interestingly, B-factors and evolutionary conservation of amino acid sequences, both typically used to distinguish between regions of high and low structural rigidity and relevance, respectively, can be directly correlated with the efficiency of proteolytic processing. That is, regions of high structural flexibility and low sequence conservation as observed for loops LA, LB, LG and LI would suggest efficient proteolysis, while structurally rigid and conserved parts such as helices 5, 8 and 9 less efficient proteolysis. Surprisingly, our data did not yield the expected correlation as these loops were processed more poorly compared to these helices (Fig. 4). This curiosity is best explained by an evolutionarily adaptation towards sequences of poor affinity to the active site of the protease or conformational rigidity, thereby allowing the protein to maintain regions of surface exposed structural flexibility. The opposite of this feature is observed in e.g. natural serine protease inhibitors (serpins) where a conserved surface exposed loop has evolved to be a substrate of proteases that are inhibited by efficient serpin processing because one proteolytic product remains covalently bound at the catalytic Ser residue^22^.

Another rather unexpected particularity was the generation of several proteolytic products that resulted from one identical site that was cleaved with high efficiency and various additional secondary cleavage sites that were processed at lower efficiency. A related phenomenon was that larger peptidic products of this class were produced at later timepoints in contrast to the expectation that larger products would be produced early to be further degraded over time. Again, these larger products are likely the result of a high affinity cleavage site at one end and a secondary site of lower and rate limiting affinity.

The wider implications of our method are that in addition to studying protein degradation, it could be adapted to study other posttranslational modifications such as phosphorylation in dynamic protein complexes. In this case, MS would identify phosphorylated residues, instead of proteolytic products. In analogy to the current study, the incubation of the chemically denatured proteins with a protein kinase would identify all potential phosphorylation sites. The subsequent mapping of these sites over time on the intact protein complex present in various functional states would provide valuable information about the conformational differences and the resulting changes of phosphorylation patterns, the functional implications of which can be addressed experimentally.

## ONLINE METHODS

### Purification of human HTRA1 and annexin A1 (ANXA1)

As the activity of HTRA1 does not depend on its N-terminal domain^8^, we used a derivative composed of the protease and PDZ domain in this study. Human HTRA1 comprised residues 158-480 and was purified as published previously^20^ with minor alterations: HTRA1 carried an N-terminal StrepII-tag. Therefore, affinity-chromatography was performed with a strep-tactin resin (IBA Lifesciences). Human ANXA1 was expressed in *E. coli* (BL21 Rosetta 2) grown in LB medium. Protein expression was induced with 300 *μ*M IPTG for 3 h at 37°C. ANXA1 was affinity purified via a Ni-NTA superflow column (Qiagen), eluted stepwise with increasing concentrations of imidazole (12.5, 25, 50, 100 and 250 mM) and further purified by size exclusion chromatography using a Superdex 200 preparation grade column (GE Healthcare) in 20 mM HEPES, 50 mM NaCl, pH 7.5. Protein concentrations were determined via Bradford assays and SDS-PAGE.

### Time-resolved proteolysis of ANXA1

Proteolysis of folded recombinant ANXA1 was done by mixing 20 *μ*M ANXA1 with 20 *μ*M HTRA1 in 150 mM NaH_2_PO_4_, 380 mM NaCl, pH 8. Samples were taken at the various timepoints indicated. Alternatively, for the proteolysis of chemically denatured substrate, ANXA1 was denatured in 8 M urea. Denatured ANXA1 was diluted into 150 mM NaH_2_PO_4_, 380 mM NaCl, pH 8 containing 0.6 M urea before adding HTRA1. As controls, substrate and protease were mixed under denaturing conditions to keep the protease inactive. For MS-analysis 5 *μ*l of each sample were added to 30 *μ*l ice-cold acetone for precipitation at −80°C overnight. Precipitated proteins were sedimented (20,000 g, 4°C, 1h) and the supernatant was lyophilized in a SpeedVac centrifuge (Eppendorf) at 30°C for 90 min. For SDS-PAGE the samples were mixed with SDS-loading dye and 100 mM DTT, heated for 2 min at 95°C, and flash frozen in liquid nitrogen.

### LC/MS/MS

Experiments were performed on an Orbitrap Elite mass spectrometer (Thermo Fischer Scientific, Waltham, Massachusetts, USA) that was coupled to an Evosep One liquid chromatography (LC) system (Evosep Biosystems, Odense, Denmark). Analysis on the Evosep One was peformed on a commercially available EV-1064 Analytical Column – 60 & 100 samples/day (Length (LC) = 8 cm; ID = 100 *μ*m; OD = 360 mm; emitter EV-1086 Stainless steel emitter).

The LC system was equipped with two mobile phases: solvent A (0.1% formic acid, FA, in water) and solvent B (0.1% FA in acetonitrile, ACN). All solvents were of UHPLC (ultra-high-performance liquid chromatography) grade (Honeywell, Seelze, Germany). For analysis with the Evosep One, samples were first loaded onto Evotips by following the manufacturers guidelines. For peptide separation, we used the 60 samples per day gradient which has an effective gradient of 21 min.

The mass spectrometer was operated using Xcalibur software (Elite: v2.2 SP1.48). The mass spectrometer was set in the positive ion mode. Precursor ion scanning (MS1) was performed in the Orbitrap analyzer (FTMS; Fourier Transform Mass Spectrometry with the internal lock mass option turned on (lock mass was 445.120025 m/z, polysiloxane)^23^. MS2 Product ion spectra were recorded only from ions with a charge > +1 and in a data dependent fashion in the ITMS. All relevant MS settings (Resolution, scan range, AGC, ion acquisition time, charge states isolation window, fragmentation type and details, cycle time, number of scans performed, and various other settings) for the individual experiments can be found in Supplementary data 5.

### Peptide and Protein identification using MaxQuant

RAW spectra were submitted to an Andromeda^24^ search in MaxQuant (version 1.5.3.30) using the default settings e.g. search for peptides between 8 and 25 residues^25^. Label-free quantification (LFQ)^26^ and match between runs was activated. Normalization in MaxQaunt was switched off. MS/MS spectra data were searched against the ACE_0653_UP000000625_83333.fasta (4450 entries) custom database. The search database contains the Uniprot reference database for *E. coli* supplemented by the sequences of HtrA1 (Q92743) and ANXA1 (P04083, plus additional N-term His-tag). All searches included a contaminants database (as implemented in MaxQuant, 246 sequences). The contaminants database contains known MS contaminants and was included to estimate the level of contamination. Andromeda searches allowed oxidation of methionine residues (16 Da) and acetylation of the protein N-terminus (42 Da). No static modifications were set. Enzyme specificity was set to “unspecific”. The instrument type in Andromeda searches was set to Orbitrap and the precursor mass tolerance was set to ±20 ppm (first search) and ±4.5 ppm (main search). The MS/MS match tolerance was set to ±20 ppm. The peptide spectrum match FDR and the protein FDR were set to 0.01 (based on target-decoy approach). Minimum peptide length was 7 amino acids. For protein quantification, unique and razor peptides were allowed. In addition to unmodified peptides, modified peptides with dynamic modifications were allowed for quantification. The minimum score for modified peptides was set to 40.

### UMSAP

MS-data were evaluated with the Targeted Proteolysis module of UMSAP 2.1.0^17^. Mass spec data of the ANXA1 alone samples served as reference for all UMSAP calculations. The significance level was set to 0.05 and the minimum score value to 50. A log2 transformation was applied to the data previous to the analysis. The amino acid distribution around the cleavage sites included 5 residues in each direction. Chain A of the PDB file 1HM6 was used for mapping of the cleavage sites to the ANXA1 structure. The native and recombinant sequences used for this calculation can be found in Supplementary data 1. Individual samples that were below the cutoff of 0.4 in Pearson correlation analyses were excluded^17^.

### Calculation of the relative cleavage frequency by UMSAP

The calculation of the cleavage frequency is a new feature of UMSAP that will be included in version 2.2.0. Therefore, we provide here a brief outlook about how the calculation is done. The calculation is performed in 2 steps. First, UMSAP groups all MS-detected peptides that share the same P1-P1’ bond. Subsequently, the relative cleavage frequency is calculated as follows. For each peptide and experiment the average intensities are calculated. Subsequently, for each peptide the average intensity ratios are calculated taking as reference the first average intensity greater than zero along the timepoints for each peptide. Finally, the relative cleavage frequency for a P1 site at a timepoint is calculated as the sum of the average intensity ratios of all peptides that share the same P1 site. If the peptide was not detected in a timepoint or the intensity values are not significantly different to the control experiments the average intensity is set to zero for this timepoint. A numeric example is provided in Supplementary data 6.

### CD spectroscopy

CD spectra were recorded on a Jasco J-170 CD spectrometer in a 1 mm quartz cuvette at 37 °C from 190-260nm, averaging 5 scans. Samples consisted of 200 *μ*L with 5 *μ*M ANXA1, 5 *μ*M HTRA1 or 5 *μ*M ANXA1 and 5 *μ*M HTRA1 together in 150 mM NaH_2_PO_4_ pH 8, 380 mM NaCl. The secondary structure composition was analyzed using DichroWeb (http://dichroweb.cryst.bbk.ac.uk) employing the CDSSTR algorithm^27,28^. Kinetic measurements were performed with 5 *μ*M ANXA1 and varying ratios of HTRA1 (HTRA1:ANXA1 = 1:1 to 1:10) or each separate protein (5 *μ*M) as control. The CD signal was monitored at 222 nm, which is characteristic for α-helices. Measurements over the timeframe of 5 h were collected with a data pitch of 5 s and a response time of 1 s, while measurements over the course of 60 min were collected with a data pitch of 0.5 s and response time of 0.5 s. The data were normalized to represent the relative signal decrease at 222 nm and smoothed using GraphPad Prism software version 8.4.1.

## ACKNOWLEDGEMENTS

We thank Helmut Tourné and Svenja Heimann for technical assistance. This work was supported by Deutsche Forschungsgemeinschaft: EH100/18-1 to ME and CRC1093 to MK.

## AUTHOR CONTRIBUTIONS

J.S. designed and carried out experiments and analyzed data; M.C. and C.B. carried out experiments and analyzed data; G.H., F.K. and M.K performed mass spectrometry; K.B.-R. developed UMSAP and analyzed data; S.B. analyzed data; J.S., H.L., R.H., D.H., S.B. and M.E. contributed to the conception of the project, the interpretation of data and wrote the manuscript.

## COMPETING INTERESTS

The authors declare no competing interests.

## ADDITIONAL INFORMATION

### Data availability

The mass spectrometry proteomics data for the digestions have been deposited to the ProteomeXchange Consortium via the PRIDE partner repository (https://www.ebi.ac.uk/pride/archive/) with the dataset identifier PXD031534. During the review process the data can be accessed via a reviewer account (Username; Password: 0R6FNeYX).

## Notes

### Competing Interest Statement

The authors have declared no competing interest.

