## Supplementary data for "High resolution analysis of proteolytic substrate processing"

**Supplementary data 1**
Sequences of native and recombinant ANXA1
Alignment of proteolytic products of denatured substrate to the primary amino acid sequence of ANXA1

**Supplementary data 2**
Multiple sequence alignment of ANXA1 proteins

**Supplementary data 3**
Alignment of proteolytic products of folded substrate to the primary amino acid sequence of ANXA1

**Supplementary data 4**
Alignment of proteolytic products of folded substrate obtained with 1, 2 and 4 $\mu$M HTRA1 to the primary amino acid sequence of ANXA1

**Supplementary data 5**
Overview of MS samples

**Supplementary data 6**
Numeric example for how UMSAP calculates the relative frequency of cleavages

**Supplementary Figures 1 - 5**
**SI Fig. 1.** Histogram of relative cleavage efficiencies.
**SI Fig. 2.** Relative amino acid distribution at sites P1-P5 and P1'-P5' after various times of incubation.
**SI Fig. 3.** Consensus sequences derived from relative amino acid distribution at sites P1-P5 and P1'-P5' after various times of incubation.
**SI Fig. 4.** Proteolysis of folded ANXA1 at low HTRA1 concentrations
**SI Fig. 5.** Circular dichroism spectroscopy

**Supplementary Table 1**
List of P1 residues with second highest rates of cuts

**Supplementary Table 2**
Comparison of the relative numbers of cuts at selected P1 residues in denatured and folded substrate

### Supplementary data 1

**Native ANXA1 sequence**

MAMVSEFLKQAWFIENEEQEYVQTVKSSKGGPGSAVSPYPTFNPSSDVAALHKAIMVKGVDEATIIDILTKRNNA
QRQQIKAAYLQETGKPLDETLKKALTGHLEEVVLALLKTPAQFDADELRAAMKGLGTDEDTLIEILASRTNKEIR
DINRVYREELKRDLAKDITSDTSGDFRNALLSLAKGDRSEDFGVNEDLADSDARALYEAGERRKGTDVNVFNTIL
TTRSYPQLRRVFQKYTKYSKHDMNKVLDLELKGDIEKCLTAIVKCATSKPAFFAEKLHQAMKGVGTRHKALIRIM
VSRSEIDMNDIKAFYQKMYGISLCQAILDETKGDYEKILVALCGGN

**Recombinant ANXA1 sequence**

MGHHHHHHHHHHSSGHIDDDDKHMAMVSEFLKQAWFIENEEQEYVQTVKSSKGGPGSAVSPYPTFNPSSDVAALH
KAIMVKGVDEATIIDILTKRNNAQRQQIKAAYLQETGKPLDETLKKALTGHLEEVVLALLKTPAQFDADELRAAM
KGLGTDEDTLIEILASRTNKEIRDINRVYREELKRDLAKDITSDTSGDFRNALLSLAKGDRSEDFGVNEDLADSD
ARALYEAGERRKGTDVNVFNTILTTRSYPQLRRVFQKYTKYSKHDMNKVLDLELKGDIEKCLTAIVKCATSKPAF
FAEKLHQAMKGVGTRHKALIRIMVSRSEIDMNDIKAFYQKMYGISLCQAILDETKGDYEKILVALCGGN

**Denatured ANXA1 0 sec**

```
1        10        20        30        40        50
MAMVSEFLKQAWFIENEEQEYVQTVKSSKGGPGSAVSPYPTFNPSSDVAA

51       60        70        80        90        100
LHKAIMVKGVDEATIIDILTKRNNAQRQQIKAAYLQETGKPLDETLKKAL

101      110       120       130       140       150
TGHLEEVVLALLKTPAQFDADELRAAMKGLGTDEDTLIEILASRTNKEIR
                          MKGLGTDEDTLIEI

151      160       170       180       190       200
DINRVYREELKRDLAKDITSDTSGDFRNALLSLAKGDRSEDFGVNEDLAD

201      210       220       230       240       250
SDARALYEAGERRKGTDVNVFNTILTTRSYPQLRRVFQKYTKYSKHDMNK

251      260       270       280       290       300
VLDLELKGDIEKCLTAIVKCATSKPAFFAEKLHQAMKGVGTRHKALIRIM

301      310       320       330       340   346
VSRSEIDMNDIKAFYQKMYGISLCQAILDETKGDYEKILVALCGGN
 SRSEIDMNDIKA
```

**Denatured ANXA1 15 sec**

```
1        10        20        30        40        50
MAMVSEFLKQAWFIENEEQEYVQTVKSSKGGPGSAVSPYPTFNPSSDVAA
   VSEFLKQAWFIENEEQEYVQTV
                        VKSSKGGPGSAVSPYPT
                         KSSKGGPGSAVSPYPT
                           SKGGPGSAVSPYPT
                                         FNPSSDVAA
                                         FNPSSDVAA

51       60        70        80        90        100
LHKAIMVKGVDEATIIDILTKRNNAQRQQIKAAYLQETGKPLDETLKKAL
LHKA
LHKAIMVKGVDEAT
                                AYLQETGKPLD
                                AYLQETGKPLDET
                                AYLQETGKPLDETLK
                                AYLQETGKPLDETLKKAL

101      110       120       130       140       150
TGHLEEVVLALLKTPAQFDADELRAAMKGLGTDEDTLIEILASRTNKEIR
TG
                QFDADELRAAMKGLGTDEDTLIEI

151      160       170       180       190       200
DINRVYREELKRDLAKDITSDTSGDFRNALLSLAKGDRSEDFGVNEDLAD
              AKDITSDTSGDFRNALL

201      210       220       230       240       250
SDARALYEAGERRKGTDVNVFNTILTTRSYPQLRRVFQKYTKYSKHDMNK
                       ILTTRSYPQLRRV

251      260       270       280       290       300
VLDLELKGDIEKCLTAIVKCATSKPAFFAEKLHQAMKGVGTRHKALIRIM
                    ATSKPAFFAEKLHQ
                    ATSKPAFFAEKLHQA
                    ATSKPAFFAEKLHQAMKGV
                     TSKPAFFAEKL
                      SKPAFFAEKL

301      310       320       330       340   346
VSRSEIDMNDIKAFYQKMYGISLCQAILDETKGDYEKILVALCGGN
 SRSEIDMNDIKA
                        QAILDETKGDYEKILVALCGGN
                          ILDETKGDYEKILVALC
                          ILDETKGDYEKILVALCGGN
                           LDETKGDYEKILVALCGGN
```

**Denatured ANXA1 30 sec**

```
1        10        20        30        40        50
MAMVSEFLKQAWFIENEEQEYVQTVKSSKGGPGSAVSPYPTFNPSSDVAA
 AMVSEFLKQA
     EFLKQAWFIENEEQEYVQTV
          AWFIENEEQEYVQTV
           WFIENEEQ
           WFIENEEQEYVQTV
                        VKSSKGGPGSAVSPYPT
                         KSSKGGPGSAVSPYPT
                           SKGGPGSAVSPYPT
                            KGGPGSAVSPYPT
                                         FNPSSDVAA
                                         FNPSSDVAA
                                         FNPSSDVAA
                                         FNPSSDVAA
                                         FNPSSDVAA

51       60        70        80        90        100
LHKAIMVKGVDEATIIDILTKRNNAQRQQIKAAYLQETGKPLDETLKKAL
LHKA
LHKAIMV
LHKAIMVK
LHKAIMVKGVDEA
LHKAIMVKGVDEAT
                                AYLQETGKP
                                AYLQETGKPL
                                AYLQETGKPLD
                                AYLQETGKPLDE
                                AYLQETGKPLDET
                                AYLQETGKPLDETL
                                AYLQETGKPLDETLK
                                AYLQETGKPLDETLKKAL
                                AYLQETGKPLDETLKKAL
                                AYLQETGKPLDETLKKAL
                                AYLQETGKPLDETLKKAL
                                 YLQETGKPLDETLKKA
                                   QETGKPLDETLKKAL
                                      GKPLDETLKKAL

101      110       120       130       140       150
TGHLEEVVLALLKTPAQFDADELRAAMKGLGTDEDTLIEILASRTNKEIR
TG
TGHL
TGHLEEV
TGHLEEV
TGHLEEV
       VLALLKTPA
       VLALLKTPAQFDADELRAA
                       RAAMKGLGTDEDTLIEI
                       RAAMKGLGTDEDTLIEIL
                         AMKGLGTDEDTLIEI
                         AMKGLGTDEDTLIEILA
                         AMKGLGTDEDTLIEILASRT

151      160       170       180       190       200
DINRVYREELKRDLAKDITSDTSGDFRNALLSLAKGDRSEDFGVNEDLAD
          KRDLAKDITSDTSGDFRNALL
              AKDITSDTSGDFRNALL
                               SLAKGDRSEDFGVNEDLAD
```

```
201      210       220       230       240       250
SDARALYEAGERRKGTDVNVFNTILTTRSYPQLRRVFQKYTKYSKHDMNK
SDARA
     LYEAGERRKGTDVNVFNT
         GERRKGTDVNVFNTILT
                  NVFNTILTTRSYPQLRRV
                   VFNTILTTRSYPQLRRV
                       ILTTRSYPQLRRV
                          TRSYPQLRRV
                                    FQKYTKYSKHDMNK
                                    FQKYTKYSKHDMNK

251      260       270       280       290       300
VLDLELKGDIEKCLTAIVKCATSKPAFFAEKLHQAMKGVGTRHKALIRIM
V
VLDL
             LTAIVKCA
                    ATSKPAFFAEKL
                    ATSKPAFFAEKLHQ
                    ATSKPAFFAEKLHQA
                    ATSKPAFFAEKLHQAM
                    ATSKPAFFAEKLHQAMKGV
                     TSKPAFFAEKL
                                HQAMKGVGTRHKALIRIM

301      310       320       330       340   346
VSRSEIDMNDIKAFYQKMYGISLCQAILDETKGDYEKILVALCGGN
VSRSEIDMNDIKA
VSRSEIDMNDIKAFYQ
VSRSEIDMNDIKAFYQKM
 SRSEIDMNDIKA
                        QAILDETKGDYEKILVALC
                        QAILDETKGDYEKILVALCGGN
                          ILDETKGDYEKILV
                          ILDETKGDYEKILVA
                          ILDETKGDYEKILVAL
                          ILDETKGDYEKILVALC
                          ILDETKGDYEKILVALCGGN
                           LDETKGDYEKILV
                           LDETKGDYEKILVALCGGN
                            DETKGDYEKILV
                               KGDYEKILVALCGGN
```

**Denatured ANXA1 60 sec**

```
1        10        20        30        40        50
MAMVSEFLKQAWFIENEEQEYVQTVKSSKGGPGSAVSPYPTFNPSSDVAA
 AMVSEFLKQ
 AMVSEFLKQA
     EFLKQAWFIENEEQEYVQTV
        KQAWFIENEEQEYVQTV
          AWFIENEEQEYVQTV
           WFIENEEQ
           WFIENEEQEYVQTV
           WFIENEEQEYVQTVKS
           WFIENEEQEYVQTVKSS
           WFIENEEQEYVQTVKSSKGGPGSA
                   EYVQTVKSSKGGPGSAVSPYPT
                       TVKSSKGGPGSAVSPYPT
                        VKSSKGGPGSAVSPYPT
                        VKSSKGGPGSAVSPYPTFNPSSDVA
                         KSSKGGPGSAVSPYPT
                         KSSKGGPGSAVSPYPTFNPSSD
                         KSSKGGPGSAVSPYPTFNPSSDV
                         KSSKGGPGSAVSPYPTFNPSSDVA
                         KSSKGGPGSAVSPYPTFNPSSDVAA
                           SKGGPGSAVSPYPT
                            KGGPGSAVSPYPT
                                       PTFNPSSDVAA
                                         FNPSSDVAA
                                         FNPSSDVAA
                                         FNPSSDVAA
                                         FNPSSDVAA
                                         FNPSSDVAA
                                         FNPSSDVAA
                                         FNPSSDVAA
                                          NPSSDVAA
51       60        70        80        90        100
LHKAIMVKGVDEATIIDILTKRNNAQRQQIKAAYLQETGKPLDETLKKAL
LHKA
L
LHKA
LHKAIMV
LHKAIMVK
LHKAIMVKGVDEA
LHKAIMVKGVDEAT
LHKA
    IMVKGVDEATI
    IMVKGVDEATIIDIL
    IMVKGVDEATIIDILT
    IMVKGVDEATIIDILTK
    IMVKGVDEATIIDILTKRN
    IMVKGVDEATIIDILTKRNNAQ
       KGVDEATIIDILTKRNNAQRQQIKA
                   TKRNNAQRQQIKAAYLQETGKPLDE
                       NAQRQQIKAAYL
                                AYLQETGKP
                                AYLQETGKPL
                                AYLQETGKPLD
                                AYLQETGKPLDE
                                AYLQETGKPLDET
                                AYLQETGKPLDETL
                                AYLQETGKPLDETLK
                                AYLQETGKPLDETLKKA
                                AYLQETGKPLDETLKKAL
```

```
                                AYLQETGKPLDETLKKAL
                                AYLQETGKPLDETLKKAL
                                AYLQETGKPLDETLKKAL
                                AYLQETGKPLDETLKKAL
                                AYLQETGKPLDETLKKAL
                                AYLQETGKPLDETLKKAL
                                AYLQETGKPLDETLKKAL
                                 YLQETGKPLDETL
                                 YLQETGKPLDETLKKA
                                 YLQETGKPLDETLKKAL
                                 YLQETGKPLDETLKKAL
                                   QETGKPLDETLKKAL
                                      GKPLDETLKKAL
                                        PLDETLKKAL
                                             LKKAL
                                                 L
                                                 L

101      110       120       130       140       150
TGHLEEVVLALLKTPAQFDADELRAAMKGLGTDEDTLIEILASRTNKEIR
T
TG
TGH
TGHL
TGHLE
TGHLEE
TGHLEEV
TG
TGHLEEV
TGHLEEV
TGHLEEV
TGHLEEV
TGHLEEV
TGHLEEV
TGHLEEV
TGHLEEVVLA
       VLALLKTP
       VLALLKTPA
       VLALLKTPAQ
       VLALLKTPAQF
       VLALLKTPAQFD
       VLALLKTPAQFDA
       VLALLKTPAQFDAD
       VLALLKTPAQFDADE
       VLALLKTPAQFDADEL
       VLALLKTPAQFDADELR
       VLALLKTPAQFDADELRA
       VLALLKTPAQFDADELRAA
       VLALLKTPAQFDADELRAAM
         ALLKTPAQFDADEL
         ALLKTPAQFDADELRA
          LLKTPAQFDADEL
          LLKTPAQFDADELRA
            KTPAQFDADEL
            KTPAQFDADELRA
                QFDADELRAAMKGLGTDEDTLIEI
                       RAAMKGLGTDEDTLIEI
                       RAAMKGLGTDEDTLIEIL
                       RAAMKGLGTDEDTLIEILASRT
                         AMKGLGTDEDTLIEI
                         AMKGLGTDEDTLIEIL
                         AMKGLGTDEDTLIEILA
                         AMKGLGTDEDTLIEILASRT
```

```
                                        LASRTNKEIR

151      160       170       180       190       200
DINRVYREELKRDLAKDITSDTSGDFRNALLSLAKGDRSEDFGVNEDLAD
DINRVYREEL
          KRDLAKDITSDTSGDFRNALL
              AKDITSDTSGDFRNALL
               KDITSDTSGDFRNALL
                  TSDTSGDFRNALL
                   SDTSGDFRNALL
                   SDTSGDFRNALLSLA
                      SGDFRNALLSLA
                               SLAKGDRSEDFGVNEDLA
                               SLAKGDRSEDFGVNEDLAD

201      210       220       230       240       250
SDARALYEAGERRKGTDVNVFNTILTTRSYPQLRRVFQKYTKYSKHDMNK
SDARA
     LYEAGERRKGTDVNVFNT
     LYEAGERRKGTDVNVFNTILT
         GERRKGTDVNVFNT
         GERRKGTDVNVFNTILT
                  NVFNTILTTRSYPQLRRV
                   VFNTILTTRSYPQLRRV
                       ILTTRSYPQLRRV
                          TRSYPQLRRV
                                    FQKYTKYSKHDMNK
                                    FQKYTKYSKHDMNK
                                    FQKYTKYSKHDMNK
                                    FQKYTKYSKHDMNK

251      260       270       280       290       300
VLDLELKGDIEKCLTAIVKCATSKPAFFAEKLHQAMKGVGTRHKALIRIM
V
VL
VLDL
VLDLEL
             LTAIVKCA
             LTAIVKCATSKPAFFAEKL
                    ATSKPAFFAEKL
                    ATSKPAFFAEKLHQ
                    ATSKPAFFAEKLHQA
                    ATSKPAFFAEKLHQAM
                    ATSKPAFFAEKLHQAMKGV
                    ATSKPAFFAEKLHQAMKGVGT
                     TSKPAFFAEKL
                     TSKPAFFAEKLHQAMKGV
                          FFAEKLHQ
                          FFAEKLHQA
                          FFAEKLHQAMKGV
                          FFAEKLHQAMKGVGTRHKALIRI
                          FFAEKLHQAMKGVGTRHKALIRIM
                                HQAMKGVGTRHKALIRI
                                HQAMKGVGTRHKALIRIM
                                HQAMKGVGTRHKALIRIM
                                  AMKGVGTRHKALIRIM
                                   MKGVGTRHKALIRIM
                                       GTRHKALIRIM
                                                 M
                                                 M

301      310       320       330       340   346
```

```
VSRSEIDMNDIKAFYQKMYGISLCQAILDETKGDYEKILVALCGGN
V
VSRSEIDMNDIKA
VSRSEIDMNDIKAFYQ
VSRSEIDMNDIKA
VSRSEIDMNDIKAFYQ
VSRSEIDMNDIKAFYQKM
VSRSEIDMNDIKAFYQKMYGISLC
 SRSEIDMNDIKA
 SRSEIDMNDIKAFYQ
             FYQKMYGISLC
             FYQKMYGISLCQA
                        QAILDETKGDYEKILV
                        QAILDETKGDYEKILVA
                        QAILDETKGDYEKILVALC
                        QAILDETKGDYEKILVALCGGN
                          ILDETKGDYEKILV
                          ILDETKGDYEKILVA
                          ILDETKGDYEKILVAL
                          ILDETKGDYEKILVALC
                          ILDETKGDYEKILVALCGGN
                           LDETKGDYEKILV
                           LDETKGDYEKILVALCGGN
                            DETKGDYEKILV
                               KGDYEKILVALC
                               KGDYEKILVALCGGN
```

**Denatured ANXA1 90 sec (5)**

```
1        10        20        30        40        50
MAMVSEFLKQAWFIENEEQEYVQTVKSSKGGPGSAVSPYPTFNPSSDVAA
 AMVSEFLKQ
 AMVSEFLKQA
    SEFLKQAWFIENEEQEYVQTV
     EFLKQAWFIENEEQEYVQTV
        KQAWFIENEEQEYVQTV
          AWFIENEEQ
          AWFIENEEQEYVQTV
           WFIENEEQ
           WFIENEEQEYVQT
           WFIENEEQEYVQTV
           WFIENEEQEYVQTVKS
           WFIENEEQEYVQTVKSS
           WFIENEEQEYVQTVKSSKGGPGSA
                   EYVQTVKSSKGGPGSAVSPYPT
                       TVKSSKGGPGSAVSPYPT
                        VKSSKGGPGSAVSPYPT
                        VKSSKGGPGSAVSPYPTFNPSSDVA
                         KSSKGGPGSAVSPYPT
                         KSSKGGPGSAVSPYPTFNPSSD
                         KSSKGGPGSAVSPYPTFNPSSDV
                         KSSKGGPGSAVSPYPTFNPSSDVA
                         KSSKGGPGSAVSPYPTFNPSSDVAA
                           SKGGPGSAVSPYPT
                            KGGPGSAVSPYPT
                                     PTFNPSSDVAA
                                         FNPSSDVAA
                                         FNPSSDVAA
                                         FNPSSDVAA
                                         FNPSSDVAA
                                         FNPSSDVAA
                                         FNPSSDVAA
                                          NPSSDVAA
                                           PSSDVAA
51       60        70        80        90        100
LHKAIMVKGVDEATIIDILTKRNNAQRQQIKAAYLQETGKPLDETLKKAL
LHKA
L
LHKA
LHKAIMV
LHKAIMVK
LHKAIMVKGVDEA
LHKA
LHKA
    IMVKGVDEA
    IMVKGVDEATI
    IMVKGVDEATIIDI
    IMVKGVDEATIIDIL
    IMVKGVDEATIIDILT
    IMVKGVDEATIIDILTKR
    IMVKGVDEATIIDILTKRN
    IMVKGVDEATIIDILTKRNNAQ
       KGVDEATIIDILTKRNNAQRQQIKA
               IDILTKRNNAQRQQIKA
                   TKRNNAQRQQIKAAYLQET
                   TKRNNAQRQQIKAAYLQETGKPLDE
                       NAQRQQIKAAYL
                       NAQRQQIKAAYLQET
                                AYLQETGKP
```

```
                                AYLQETGKPL
                                AYLQETGKPLD
                                AYLQETGKPLDE
                                AYLQETGKPLDET
                                AYLQETGKPLDETL
                                AYLQETGKPLDETLK
                                AYLQETGKPLDETLKKA
                                AYLQETGKPLDETLKKAL
                                AYLQETGKPLDETLKKAL
                                AYLQETGKPLDETLKKAL
                                AYLQETGKPLDETLKKAL
                                AYLQETGKPLDETLKKAL
                                AYLQETGKPLDETLKKAL
                                AYLQETGKPLDETLKKAL
                                AYLQETGKPLDETLKKAL
                                 YLQETGKPLDET
                                 YLQETGKPLDETL
                                 YLQETGKPLDETLKKA
                                 YLQETGKPLDETLKKAL
                                   QETGKPLDETLKKAL
                                      GKPLDETLKKAL
                                        PLDETLKKAL
                                             LKKAL
                                                 L
                                                 L

101      110       120       130       140       150
TGHLEEVVLALLKTPAQFDADELRAAMKGLGTDEDTLIEILASRTNKEIR
T
TG
TGH
TGHL
TGHLE
TGHLEE
TGHLEEV
TGHLEEV
TG
TGHLEEV
TGHLEEV
TGHLEEV
TGHLEEV
TGHLEEVVLA
       VLALLKTP
       VLALLKTPA
       VLALLKTPAQ
       VLALLKTPAQF
       VLALLKTPAQFD
       VLALLKTPAQFDA
       VLALLKTPAQFDAD
       VLALLKTPAQFDADE
       VLALLKTPAQFDADEL
       VLALLKTPAQFDADELR
       VLALLKTPAQFDADELRA
       VLALLKTPAQFDADELRAA
       VLALLKTPAQFDADELRAAM
        LALLKTPAQFDADELRA
         ALLKTPAQFDADEL
         ALLKTPAQFDADELRA
          LLKTPAQFDADEL
          LLKTPAQFDADELRA
           LKTPAQFDADELRAA
            KTPAQFDADEL
            KTPAQFDADELRA
```

```
                QFDADELRA
                       RAAMKGLGTDEDTLIEI
                       RAAMKGLGTDEDTLIEIL
                       RAAMKGLGTDEDTLIEILASRT
                         AMKGLGTDEDTLIEI
                         AMKGLGTDEDTLIEIL
                         AMKGLGTDEDTLIEILA
                         AMKGLGTDEDTLIEILASRT
                                         ASRTNKEIR
                                             NKEIR
                                             NKEIR

151      160       170       180       190       200
DINRVYREELKRDLAKDITSDTSGDFRNALLSLAKGDRSEDFGVNEDLAD
DI
DINRVYREEL
DINRVYREELKRDLAKDIT
          KRDLAKDITSDTSGDFRNALL
              AKDITSDTSGDFRNALL
               KDITSDTSGDFRNALL
                  TSDTSGDFRNALL
                   SDTSGDFRNALL
                   SDTSGDFRNALLSLA
                               SLAKGDRSEDFGVNEDLA
                               SLAKGDRSEDFGVNEDLAD
                               SLAKGDRSEDFGVNEDLAD

201      210       220       230       240       250
SDARALYEAGERRKGTDVNVFNTILTTRSYPQLRRVFQKYTKYSKHDMNK
SDARA
SDARAL
     LYEAGERRKGTDVNVFNT
     LYEAGERRKGTDVNVFNTILT
         GERRKGTDVNVFNT
         GERRKGTDVNVFNTILT
                  NVFNTILTTRSYPQLRRV
                   VFNTILTTRSYPQLRRV
                       ILTTRSYPQLRRV
                          TRSYPQLRRV
                          TRSYPQLRRVFQ
                                    FQKYTKYSKHDMNK
                                    FQKYTKYSKHDMNK
                                    FQKYTKYSKHDMNK
                                    FQKYTKYSKHDMNK
                                        KYSKHDMNK

251      260       270       280       290       300
VLDLELKGDIEKCLTAIVKCATSKPAFFAEKLHQAMKGVGTRHKALIRIM
V
VL
VLDL
VLDLEL
VLDLELKGDIEKC
             LTAIVKCATSKPAFFAEKL
               AIVKCATSKPAFFAEKL
                    ATSKPAFFAEKL
                    ATSKPAFFAEKLHQ
                    ATSKPAFFAEKLHQA
                    ATSKPAFFAEKLHQAM
                    ATSKPAFFAEKLHQAMKGV
                    ATSKPAFFAEKLHQAMKGVGT
                    ATSKPAFFAEKLHQAMKGVGTRHKA
                     TSKPAFFAEKL
```

```
                     TSKPAFFAEKLHQAMKGV
                        PAFFAEKLHQA
                          FFAEKLHQ
                          FFAEKLHQA
                          FFAEKLHQAM
                          FFAEKLHQAMKGV
                          FFAEKLHQAMKGVGTRHKALIRI
                          FFAEKLHQAMKGVGTRHKALIRIM
                                HQAMKGVGTRHKALIRI
                                HQAMKGVGTRHKALIRIM
                                HQAMKGVGTRHKALIRIM
                                HQAMKGVGTRHKALIRIM
                                  AMKGVGTRHKALIRIM
                                   MKGVGTRHKALIRIM
                                       GTRHKALIRIM
                                              IRIM
                                                 M
                                                 M

301      310       320       330       340   346
VSRSEIDMNDIKAFYQKMYGISLCQAILDETKGDYEKILVALCGGN
V
VS
VSRSEIDMNDIKA
VSRSEIDMNDIKA
VSRSEIDMNDIKAFYQ
VSRSEIDMNDI
VSRSEIDMNDIKA
VSRSEIDMNDIKAFYQ
VSRSEIDMNDIKAFYQKM
VSRSEIDMNDIKAFYQKMYGI
VSRSEIDMNDIKAFYQKMYGISLC
 SRSEIDMNDIKA
 SRSEIDMNDIKAFYQ
  RSEIDMNDIKA
  RSEIDMNDIKAFYQ
             FYQKMYGI
             FYQKMYGISLC
             FYQKMYGISLCQA
                KMYGISLC
                        QAILDETKGDYEKIL
                        QAILDETKGDYEKILV
                        QAILDETKGDYEKILVA
                        QAILDETKGDYEKILVALC
                        QAILDETKGDYEKILVALCGGN
                          ILDETKGDYEKILV
                          ILDETKGDYEKILVA
                          ILDETKGDYEKILVAL
                          ILDETKGDYEKILVALC
                          ILDETKGDYEKILVALCGGN
                           LDETKGDYEKILV
                           LDETKGDYEKILVALCGGN
                            DETKGDYEKILV
                               KGDYEKILVALC
                               KGDYEKILVALCGGN
```

**Denatured ANXA1 120 sec (6)**

```
1        10        20        30        40        50
MAMVSEFLKQAWFIENEEQEYVQTVKSSKGGPGSAVSPYPTFNPSSDVAA
 AMVSEFLKQ
 AMVSEFLKQA
   VSEFLKQA
   VSEFLKQAWFIENEEQEYVQTV
    SEFLKQAWFIENEEQEYVQTV
     EFLKQAWFIENEEQEYVQTV
     EFLKQAWFIENEEQEYVQTVKS
        KQAWFIENEEQEYVQTV
          AWFIENEEQ
          AWFIENEEQEYVQTV
           WFIENEEQ
           WFIENEEQEYV
           WFIENEEQEYVQT
           WFIENEEQEYVQTV
           WFIENEEQEYVQTVKS
           WFIENEEQEYVQTVKSS
           WFIENEEQEYVQTVKSSKGGPGSA
              ENEEQEYVQTV
                   EYVQTVKSSKGGPGSAVSPYPT
                       TVKSSKGGPGSAVSPYPT
                        VKSSKGGPGSAVSPYPT
                        VKSSKGGPGSAVSPYPTFNPSSDVA
                         KSSKGGPGSAVSPYPT
                         KSSKGGPGSAVSPYPTFNPSSD
                         KSSKGGPGSAVSPYPTFNPSSDV
                         KSSKGGPGSAVSPYPTFNPSSDVA
                         KSSKGGPGSAVSPYPTFNPSSDVAA
                           SKGGPGSAVSPYPT
                            KGGPGSAVSPYPT
                                       PTFNPSSDVAA
                                         FNPSSDVAA
                                         FNPSSDVAA
                                         FNPSSDVAA
                                         FNPSSDVAA
                                         FNPSSDVAA
                                         FNPSSDVAA
                                          NPSSDVAA
                                           PSSDVAA
                                            SSDVAA
                                                 A

51       60        70        80        90        100
LHKAIMVKGVDEATIIDILTKRNNAQRQQIKAAYLQETGKPLDETLKKAL
LHKA
L
LHKA
LHKAIMV
LHKAIMVK
LHKAIMVKGVDEA
LHKA
LHKA
LHKA
LHKAIMVKGVDEATIIDIL
    IMVKGVDEA
    IMVKGVDEATI
    IMVKGVDEATIIDI
    IMVKGVDEATIIDIL
```

```
    IMVKGVDEATIIDILT
    IMVKGVDEATIIDILTK
    IMVKGVDEATIIDILTKR
    IMVKGVDEATIIDILTKRN
    IMVKGVDEATIIDILTKRNNA
    IMVKGVDEATIIDILTKRNNAQ
       KGVDEATIIDIL
       KGVDEATIIDILTKRNNAQRQQIKA
               IDILTKRNNAQRQQIKA
                   TKRNNAQRQQIKAAYL
                   TKRNNAQRQQIKAAYLQETGKPLDE
                       NAQRQQIKAAYL
                       NAQRQQIKAAYLQET
                                AYLQETGKP
                                AYLQETGKPL
                                AYLQETGKPLD
                                AYLQETGKPLDE
                                AYLQETGKPLDET
                                AYLQETGKPLDETL
                                AYLQETGKPLDETLK
                                AYLQETGKPLDETLKK
                                AYLQETGKPLDETLKKA
                                AYLQETGKPLDETLKKAL
                                AYLQETGKPLDETLKKAL
                                AYLQETGKPLDETLKKAL
                                AYLQETGKPLDETLKKAL
                                AYLQETGKPLDETLKKAL
                                AYLQETGKPLDETLKKAL
                                AYLQETGKPLDETLKKAL
                                AYLQETGKPLDETLKKAL
                                 YLQETGKPLDET
                                 YLQETGKPLDETL
                                 YLQETGKPLDETLKKA
                                 YLQETGKPLDETLKKAL
                                 YLQETGKPLDETLKKAL
                                 YLQETGKPLDETLKKAL
                                 YLQETGKPLDETLKKAL
                                   QETGKPLDETLKKAL
                                   QETGKPLDETLKKAL
                                      GKPLDETLKKAL
                                        PLDETLKKAL
                                             LKKAL
                                                AL
                                                 L
101      110       120       130       140      150
TGHLEEVVLALLKTPAQFDADELRAAMKGLGTDEDTLIEILASRTNKEIR
T
TG
TGH
TGHL
TGHLE
TGHLEE
TGHLEEV
T
TG
TGHLEEV
T
TGHLEEV
TGHLEEV
TGHLEEV
TGHLEEV
TGHLEEV
TGHLEEV
```

```
       VLALLKTP
       VLALLKTPA
       VLALLKTPAQ
       VLALLKTPAQF
       VLALLKTPAQFD
       VLALLKTPAQFDA
       VLALLKTPAQFDAD
       VLALLKTPAQFDADE
       VLALLKTPAQFDADEL
       VLALLKTPAQFDADELR
       VLALLKTPAQFDADELRA
       VLALLKTPAQFDADELRAA
       VLALLKTPAQFDADELRAAM
        LALLKTPAQFDADELRA
         ALLKTPAQFDADEL
         ALLKTPAQFDADELRA
            KTPAQFDADEL
            KTPAQFDADELRA
            KTPAQFDADELRAA
                       RAAMKGLGTDEDTLIEI
                       RAAMKGLGTDEDTLIEIL
                       RAAMKGLGTDEDTLIEILASRT
                         AMKGLGTDEDTLIEI
                         AMKGLGTDEDTLIEIL
                         AMKGLGTDEDTLIEILA
                         AMKGLGTDEDTLIEILASRT
                                        LASRTNKEIR
                                        LASRTNKEIR
                                         ASRTNKEIR
                                         ASRTNKEIR
                                             NKEIR
                                             NKEIR

151      160       170       180       190       200
DINRVYREELKRDLAKDITSDTSGDFRNALLSLAKGDRSEDFGVNEDLAD
DI
DINRVYREEL
DI
DINRVYREEL
DINRVYREEL
DINRVYREELKRDLAKDIT
          KRDLAKDITSDTSGDFRNALL
              AKDITSDTSGDFRNALL
               KDITSDTSGDFRNALL
                  TSDTSGDFRNALL
                  TSDTSGDFRNALLSLA
                   SDTSGDFRNALL
                   SDTSGDFRNALLSLA
                      SGDFRNALLSLA
                               SLAKGDRSEDFGVNEDLA
                               SLAKGDRSEDFGVNEDLAD
                               SLAKGDRSEDFGVNEDLAD
                                  KGDRSEDFGVNEDLAD
                                  KGDRSEDFGVNEDLAD

201      210       220       230       240       250
SDARALYEAGERRKGTDVNVFNTILTTRSYPQLRRVFQKYTKYSKHDMNK
SDARA
SDARAL
SDARA
SDARALYEA
     LYEAGERRKGTDVNVFNT
```

```
     LYEAGERRKGTDVNVFNTILT
         GERRKGTDVNVFNT
         GERRKGTDVNVFNTILT
                  NVFNTILTTRSYPQLRRV
                   VFNTILTTRSYPQLRRV
                       ILTTRSYPQLRRV
                          TRSYPQLRRV
                          TRSYPQLRRVFQ
                                    FQKYTKYSKHDMNK
                                    FQKYTKYSKHDMNK
                                    FQKYTKYSKHDMNK
                                    FQKYTKYSKHDMNK
                                      KYTKYSKHDMNK
                                         KYSKHDMNK

251      260       270       280       290       300
VLDLELKGDIEKCLTAIVKCATSKPAFFAEKLHQAMKGVGTRHKALIRIM
V
VL
VLDL
VLDLEL
VLDLELKGDIEKC
VLDLELKGDIEKCLT
             LTAIVKCA
             LTAIVKCATSKPA
               AIVKCATSKPAFFAEKL
                    ATSKPAFFAEKL
                    ATSKPAFFAEKLHQ
                    ATSKPAFFAEKLHQA
                    ATSKPAFFAEKLHQAM
                    ATSKPAFFAEKLHQAMKGV
                    ATSKPAFFAEKLHQAMKGVGT
                    ATSKPAFFAEKLHQAMKGVGTRHKA
                     TSKPAFFAEKL
                     TSKPAFFAEKLHQ
                     TSKPAFFAEKLHQAMKGV
                        PAFFAEKLHQA
                          FFAEKLHQ
                          FFAEKLHQA
                          FFAEKLHQAM
                          FFAEKLHQAMKGV
                          FFAEKLHQAMKGVGTRHKALIRI
                          FFAEKLHQAMKGVGTRHKALIRIM
                                HQAMKGVGTRHKALI
                                HQAMKGVGTRHKALIRI
                                HQAMKGVGTRHKALIRIM
                                HQAMKGVGTRHKALIRIM
                                HQAMKGVGTRHKALIRIM
                                  AMKGVGTRHKALIRIM
                                   MKGVGTRHKALIRIM
                                   MKGVGTRHKALIRIM
                                     GVGTRHKALIRIM
                                       GTRHKALIRIM
                                              IRIM
                                                 M
                                                 M

301      310       320       330       340   346
VSRSEIDMNDIKAFYQKMYGISLCQAILDETKGDYEKILVALCGGN
V
VS
VS
VSRSEIDMNDIKA
```

```
VSRSEIDMNDIKA
VSRSEIDMNDIKAFYQ
VSRSEIDMNDI
VSRSEIDMNDIKA
VSRSEIDMNDIKAFYQ
VSRSEIDMNDIKAFYQKM
VSRSEIDMNDIKAFYQKMYGI
VSRSEIDMNDIKAFYQKMYGISLC
 SRSEIDMNDIKA
 SRSEIDMNDIKAFYQ
  RSEIDMNDIKA
  RSEIDMNDIKAFYQ
             FYQKMYGI
             FYQKMYGISLC
             FYQKMYGISLCQA
                KMYGISLC
                        QAILDETKGDYEKIL
                        QAILDETKGDYEKILV
                        QAILDETKGDYEKILVA
                        QAILDETKGDYEKILVALC
                        QAILDETKGDYEKILVALCGGN
                          ILDETKGDYEKIL
                          ILDETKGDYEKILV
                          ILDETKGDYEKILVA
                          ILDETKGDYEKILVAL
                          ILDETKGDYEKILVALC
                          ILDETKGDYEKILVALCGGN
                           LDETKGDYEKILV
                           LDETKGDYEKILVALC
                           LDETKGDYEKILVALCGGN
                            DETKGDYEKILV
                               KGDYEKILVALC
                               KGDYEKILVALCGGN
```

**Denatured ANXA1 300 sec**

```
1        10        20        30        40        50
MAMVSEFLKQAWFIENEEQEYVQTVKSSKGGPGSAVSPYPTFNPSSDVAA
 AMVSEFLKQ
 AMVSEFLKQA
  MVSEFLKQ
  MVSEFLKQA
   VSEFLKQA
   VSEFLKQAWFIENEEQEYVQTV
    SEFLKQAWFIENEEQEYVQTV
     EFLKQAWFIENEEQEYVQTV
     EFLKQAWFIENEEQEYVQTVKS
        KQAWFIENEEQEYVQTV
          AWFIENEEQ
          AWFIENEEQEYVQT
          AWFIENEEQEYVQTV
          AWFIENEEQEYVQTVKS
           WFIENEEQ
           WFIENEEQEYV
           WFIENEEQEYVQT
           WFIENEEQEYVQTV
           WFIENEEQEYVQTVKS
           WFIENEEQEYVQTVKSS
           WFIENEEQEYVQTVKSSKGGPGSA
              ENEEQEYVQTV
                   EYVQTVKSSKGGPGSAVSPYPT
                      QTVKSSKGGPGSAVSPYPT
                       TVKSSKGGPGSAVSPYPT
                        VKSSKGGPGSAVSPYPT
                        VKSSKGGPGSAVSPYPTFNPSSD
                        VKSSKGGPGSAVSPYPTFNPSSDVA
                         KSSKGGPGSAVSPYPT
                         KSSKGGPGSAVSPYPTFNPSS
                         KSSKGGPGSAVSPYPTFNPSSD
                         KSSKGGPGSAVSPYPTFNPSSDV
                         KSSKGGPGSAVSPYPTFNPSSDVA
                         KSSKGGPGSAVSPYPTFNPSSDVAA
                          SSKGGPGSAVSPYPT
                           SKGGPGSAVSPYPT
                            KGGPGSAVSPYPT
                                       PTFNPSSDVAA
                                        TFNPSSDVAA
                                         FNPSSDVAA
                                         FNPSSDVAA
                                         FNPSSDVAA
                                         FNPSSDVAA
                                         FNPSSDVAA
                                         FNPSSDVAA
                                          NPSSDVAA
                                           PSSDVAA
                                            SSDVAA
                                                 A

51       60        70        80        90        100
LHKAIMVKGVDEATIIDILTKRNNAQRQQIKAAYLQETGKPLDETLKKAL
LHKA
LHKA
L
LHKA
LHKAIMV
```

```
LHKAIMVK
LHKAIMVKGVDEA
LHKA
LHKA
LHKA
LHKAIMVKGVDEATIIDIL
    IMVKGVDEA
    IMVKGVDEATI
    IMVKGVDEATIIDI
    IMVKGVDEATIIDIL
    IMVKGVDEATIIDILT
    IMVKGVDEATIIDILTK
    IMVKGVDEATIIDILTKR
    IMVKGVDEATIIDILTKRN
    IMVKGVDEATIIDILTKRNNA
    IMVKGVDEATIIDILTKRNNAQ
     MVKGVDEATIIDIL
      VKGVDEATIIDIL
       KGVDEATIIDIL
       KGVDEATIIDILT
       KGVDEATIIDILTKRN
       KGVDEATIIDILTKRNNAQRQQIKA
               IDILTKRNNAQRQQIKA
                   TKRNNAQRQQIKAAYL
                   TKRNNAQRQQIKAAYLQET
                   TKRNNAQRQQIKAAYLQETGKPLDE
                      NNAQRQQIKAAYL
                      NNAQRQQIKAAYLQET
                       NAQRQQIKAAYL
                       NAQRQQIKAAYLQET
                              KAAYLQET
                                AYLQETGKP
                                AYLQETGKPL
                                AYLQETGKPLD
                                AYLQETGKPLDE
                                AYLQETGKPLDET
                                AYLQETGKPLDETL
                                AYLQETGKPLDETLK
                                AYLQETGKPLDETLKKA
                                AYLQETGKPLDETLKKAL
                                AYLQETGKPLDETLKKAL
                                AYLQETGKPLDETLKKAL
                                AYLQETGKPLDETLKKAL
                                AYLQETGKPLDETLKKAL
                                AYLQETGKPLDETLKKAL
                                AYLQETGKPLDETLKKAL
                                AYLQETGKPLDETLKKAL
                                 YLQETGKPLDE
                                 YLQETGKPLDET
                                 YLQETGKPLDETL
                                 YLQETGKPLDETLKKA
                                 YLQETGKPLDETLKKAL
                                 YLQETGKPLDETLKKAL
                                 YLQETGKPLDETLKKAL
                                   QETGKPLDET
                                   QETGKPLDETLKKAL
                                   QETGKPLDETLKKAL
                                    ETGKPLDETLKKAL
                                      GKPLDETLKKAL
                                        PLDETLKKAL
                                             LKKAL
                                              KKAL
                                                AL
```

```
                                                 L

101      110       120       130       140       150
TGHLEEVVLALLKTPAQFDADELRAAMKGLGTDEDTLIEILASRTNKEIR
T
TG
TGH
TGHL
TGHLE
TGHLEE
TGHLEEV
T
TG
TGHLEEV
T
TGHLEEV
TGHLEEV
TGHLEEV
TGHLEEV
TGHLEEV
TGHLEEV
TGHLEEV
TGHLEEV
       VLALLKTP
       VLALLKTPA
       VLALLKTPAQ
       VLALLKTPAQF
       VLALLKTPAQFD
       VLALLKTPAQFDA
       VLALLKTPAQFDAD
       VLALLKTPAQFDADE
       VLALLKTPAQFDADEL
       VLALLKTPAQFDADELR
       VLALLKTPAQFDADELRA
       VLALLKTPAQFDADELRAA
       VLALLKTPAQFDADELRAAM
        LALLKTPAQFDADEL
        LALLKTPAQFDADELRA
         ALLKTPAQFDADEL
         ALLKTPAQFDADELRA
         ALLKTPAQFDADELRAA
         ALLKTPAQFDADELRAAM
          LLKTPAQFDADEL
          LLKTPAQFDADELRA
          LLKTPAQFDADELRAAM
           LKTPAQFDADEL
           LKTPAQFDADELRA
            KTPAQFDADEL
            KTPAQFDADELRA
            KTPAQFDADELRAA
                QFDADELRA
                       RAAMKGLGTDEDTLIEI
                       RAAMKGLGTDEDTLIEIL
                       RAAMKGLGTDEDTLIEILA
                       RAAMKGLGTDEDTLIEILASRT
                         AMKGLGTDEDTLIEI
                         AMKGLGTDEDTLIEIL
                         AMKGLGTDEDTLIEILA
                         AMKGLGTDEDTLIEILASRT
                          MKGLGTDEDTLIEI
                          MKGLGTDEDTLIEIL
                          MKGLGTDEDTLIEILASRT
```

```
                           KGLGTDEDTLIEIL
                                         LASRTNKEIR
                                         LASRTNKEIR
                                          ASRTNKEIR
                                          ASRTNKEIR
                                              NKEIR
                                              NKEIR
                                              NKEIR
```

```
151      160       170       180       190       200
DINRVYREELKRDLAKDITSDTSGDFRNALLSLAKGDRSEDFGVNEDLAD
DI
DINRVYREEL
DI
DINRVYREEL
DINRV
DINRVYREEL
DINRVYREELKRDLAKDIT
     YREELKRDLAKDITSDT
     YREELKRDLAKDITSDTSGDFRNA
          KRDLAKDITSDT
          KRDLAKDITSDTSGDFRNALL
          KRDLAKDITSDTSGDFRNALLSLA
              AKDITSDTSGDFRNALL
               KDITSDTSGDFRNALL
                  TSDTSGDFRNALL
                  TSDTSGDFRNALLSLA
                   SDTSGDFRNA
                   SDTSGDFRNALL
                   SDTSGDFRNALLSL
                   SDTSGDFRNALLSLA
                      SGDFRNALLSL
                      SGDFRNALLSLA
                               SLAKGDRSEDFGV
                               SLAKGDRSEDFGVNEDLA
                               SLAKGDRSEDFGVNEDLAD
                               SLAKGDRSEDFGVNEDLAD
                               SLAKGDRSEDFGVNEDLAD
                                LAKGDRSEDFGVNEDLAD
                                  KGDRSEDFGVNEDLAD
                                  KGDRSEDFGVNEDLAD
```

```
201      210       220       230       240       250
SDARALYEAGERRKGTDVNVFNTILTTRSYPQLRRVFQKYTKYSKHDMNK
SDA
SDARA
SDARAL
SDARA
SDARA
SDARALYEA
     LYEAGERRKGTDVNVFNT
     LYEAGERRKGTDVNVFNTILT
         GERRKGTDVNVFNT
         GERRKGTDVNVFNTILT
                  NVFNTILTTRSYPQLRRV
                   VFNTILTTRSYPQLRRV
                     NTILTTRSYPQLRRV
                       ILTTRSYPQL
                       ILTTRSYPQLRRV
                          TRSYPQLRRV
                          TRSYPQLRRVFQ
                                    FQKYTKYSKHDM
                                    FQKYTKYSKHDMNK
```

```
                                    FQKYTKYSKHDMNK
                                    FQKYTKYSKHDMNK
                                    FQKYTKYSKHDMNK
                                      KYTKYSKHDMNK
                                         KYSKHDMNK
                                         KYSKHDMNK
                                         KYSKHDMNK
                                            KHDMNK

251      260       270       280       290       300
VLDLELKGDIEKCLTAIVKCATSKPAFFAEKLHQAMKGVGTRHKALIRIM
V
VL
VLDL
VLDLEL
VLDLELKGDIEKC
VLDL
VLDLELKGDIEKC
VLDLELKGDIEKCLT
VLDLELKGDIEKC
    ELKGDIEKC
    ELKGDIEKCLTAIVKC
      KGDIEKCLTAIVKC
             LTAIVKCA
             LTAIVKCATSKPA
             LTAIVKCATSKPAFFAEKL
               AIVKCATSKPAFFAEKL
                 VKCATSKPAFFAEKL
                  KCATSKPAFFAEKL
                    ATSKPAFFAEKL
                    ATSKPAFFAEKLHQ
                    ATSKPAFFAEKLHQA
                    ATSKPAFFAEKLHQAM
                    ATSKPAFFAEKLHQAMKGV
                    ATSKPAFFAEKLHQAMKGVGT
                    ATSKPAFFAEKLHQAMKGVGTRHKA
                     TSKPAFFAEKL
                     TSKPAFFAEKLHQ
                     TSKPAFFAEKLHQAMKGV
                      SKPAFFAEKL
                        PAFFAEKLHQA
                          FFAEKLHQ
                          FFAEKLHQA
                          FFAEKLHQAM
                          FFAEKLHQAMKGV
                          FFAEKLHQAMKGVGTRHKALIRI
                          FFAEKLHQAMKGVGTRHKALIRIM
                          FFAEKLHQAMKGVGTRHKALIRIM
                                HQAMKGVGTRHKALI
                                HQAMKGVGTRHKALIRI
                                HQAMKGVGTRHKALIRIM
                                HQAMKGVGTRHKALIRIM
                                HQAMKGVGTRHKALIRIM
                                  AMKGVGTRHKALIRIM
                                   MKGVGTRHKALIRIM
                                   MKGVGTRHKALIRIM
                                     GVGTRHKALIRIM
                                       GTRHKALIRIM
                                              IRIM
                                               RIM
                                                 M
                                                 M
```

```
                                                 M
                                                 M
                                                 M

301      310       320       330       340   346
VSRSEIDMNDIKAFYQKMYGISLCQAILDETKGDYEKILVALCGGN
V
V
VS
VS
VSRSEIDMNDIKA
VSRSEIDMNDIKA
VSRSEIDMNDI
VSRSEIDMNDIKA
VSRSEIDMNDIKAFYQ
VSRSEIDMNDIKAFYQKM
VSRSEIDMNDIKAFYQKMYGISLC
VSRSEIDMNDI
VSRSEIDMNDIKA
VSRSEIDMNDIKAFYQ
VSRSEIDMNDIKAFYQKM
VSRSEIDMNDIKAFYQKMYGI
VSRSEIDMNDIKAFYQKMYGISLC
 SRSEIDMNDIKA
 SRSEIDMNDIKAFYQ
 SRSEIDMNDIKAFYQKM
 SRSEIDMNDIKAFYQKMYGISLC
  RSEIDMNDIKA
  RSEIDMNDIKAFYQ
    EIDMNDIKA
             FYQKMYGI
             FYQKMYGISLC
             FYQKMYGISLCQA
                KMYGISLC
                KMYGISLCQA
                        QAILDETKGDYEKI
                        QAILDETKGDYEKIL
                        QAILDETKGDYEKILV
                        QAILDETKGDYEKILVA
                        QAILDETKGDYEKILVALC
                        QAILDETKGDYEKILVALCGGN
                          ILDETKGDYEKIL
                          ILDETKGDYEKILV
                          ILDETKGDYEKILVA
                          ILDETKGDYEKILVAL
                          ILDETKGDYEKILVALC
                          ILDETKGDYEKILVALCGGN
                           LDETKGDYEKILV
                           LDETKGDYEKILVALC
                           LDETKGDYEKILVALCGGN
                            DETKGDYEKILV
                               KGDYEKILVALC
                               KGDYEKILVALCGGN
```

**Denatured ANXA1 600 sec**

```
1        10        20        30        40        50
MAMVSEFLKQAWFIENEEQEYVQTVKSSKGGPGSAVSPYPTFNPSSDVAA
 AMVSEFLKQ
 AMVSEFLKQA
  MVSEFLKQ
  MVSEFLKQA
   VSEFLKQA
VSEFLKQAWFIENEEQEYVQTV
 SEFLKQAWFIENEEQEYVQTV
  EFLKQAWFIENEEQEYVQT
  EFLKQAWFIENEEQEYVQTV
  EFLKQAWFIENEEQEYVQTVKS
     KQAWFIENEEQEYVQTV
       AWFIENEEQ
       AWFIENEEQEYVQT
       AWFIENEEQEYVQTV
       AWFIENEEQEYVQTVKS
        WFIENEEQ
        WFIENEEQEYV
        WFIENEEQEYVQ
        WFIENEEQEYVQT
        WFIENEEQEYVQTV
        WFIENEEQEYVQTVKS
        WFIENEEQEYVQTVKSS
        WFIENEEQEYVQTVKSSKGGPGSA
           ENEEQEYVQTV
                EYVQTVKSSKGGPGSAVSPYPT
                   QTVKSSKGGPGSAVSPYPT
                    TVKSSKGGPGSAVSPYPT
                     VKSSKGGPGSAVSPYPT
                     VKSSKGGPGSAVSPYPTFNPSSD
                     VKSSKGGPGSAVSPYPTFNPSSDVA
                      KSSKGGPGSAVSPYPT
                      KSSKGGPGSAVSPYPTFNPSS
                      KSSKGGPGSAVSPYPTFNPSSD
                      KSSKGGPGSAVSPYPTFNPSSDV
                      KSSKGGPGSAVSPYPTFNPSSDVA
                       SSKGGPGSAVSPYPT
                        SKGGPGSAVSPYPT
                         KGGPGSAVSPYPT
                                    PTFNPSSDVAA
                                     TFNPSSDVAA
                                      FNPSSDVAA
                                      FNPSSDVAA
                                      FNPSSDVAA
                                      FNPSSDVAA
                                      FNPSSDVAA
                                      FNPSSDVAA
                                      FNPSSDVAA
                                       NPSSDVAA
                                        PSSDVAA
                                         SSDVAA
                                              A

51       60        70        80        90        100
LHKAIMVKGVDEATIIDILTKRNNAQRQQIKAAYLQETGKPLDETLKKAL
LHKA
LHKA
L
LHKA
LHKAIM
```

```
LHKAIMV
LHKAIMVK
LHKAIMVKGVDEA
LHKA
LHKA
LHKA
LHKAIMVKGVDEATIIDIL
    IMVKGVDEA
    IMVKGVDEATI
    IMVKGVDEATIIDI
    IMVKGVDEATIIDIL
    IMVKGVDEATIIDILT
    IMVKGVDEATIIDILTK
    IMVKGVDEATIIDILTKR
    IMVKGVDEATIIDILTKRN
    IMVKGVDEATIIDILTKRNNA
    IMVKGVDEATIIDILTKRNNAQ
    IMVKGVDEATIIDILTKRNNAQRQ
     MVKGVDEATIIDIL
      VKGVDEATIIDIL
      VKGVDEATIIDILT
      VKGVDEATIIDILTKRN
       KGVDEATIIDIL
       KGVDEATIIDILT
       KGVDEATIIDILTKRNNAQRQQIKA
               IDILTKRNNAQRQQIKA
                   TKRNNAQRQQIKAAYL
                   TKRNNAQRQQIKAAYLQET
                   TKRNNAQRQQIKAAYLQETGKPLDE
                    KRNNAQRQQIKAAYL
                      NNAQRQQIKAAYL
                      NNAQRQQIKAAYLQET
                       NAQRQQIKAAYL
                       NAQRQQIKAAYLQET
                              KAAYLQET
                                AYLQETGKP
                                AYLQETGKPL
                                AYLQETGKPLD
                                AYLQETGKPLDE
                                AYLQETGKPLDET
                                AYLQETGKPLDETL
                                AYLQETGKPLDETLK
                                AYLQETGKPLDETLKKA
                                AYLQETGKPLDETLKKAL
                                AYLQETGKPLDETLKKAL
                                AYLQETGKPLDETLKKAL
                                AYLQETGKPLDETLKKAL
                                AYLQETGKPLDETLKKAL
                                AYLQETGKPLDETLKKAL
                                AYLQETGKPLDETLKKAL
                                 YLQETGKPLDE
                                 YLQETGKPLDET
                                 YLQETGKPLDETLKKA
                                 YLQETGKPLDETLKKAL
                                 YLQETGKPLDETLKKAL
                                 YLQETGKPLDETLKKAL
                                   QETGKPLDET
                                   QETGKPLDETLKKAL
                                   QETGKPLDETLKKAL
                                    ETGKPLDETLKKAL
                                      GKPLDETLKKAL
                                        PLDETLKKAL
                                             LKKAL
```

```
                                              KKAL
                                                AL
                                                 L

101      110       120       130       140       150
TGHLEEVVLALLKTPAQFDADELRAAMKGLGTDEDTLIEILASRTNKEIR
T
TG
TGH
TGHL
TGHLE
TGHLEEV
T
TG
TGHLEEV
T
TGHLEEV
TGHLEEV
TGHLEEV
TGHLEEV
TGHLEEV
TGHLEEV
TGHLEEV
TGHLEEV
       VLALLKTP
       VLALLKTPA
       VLALLKTPAQ
       VLALLKTPAQF
       VLALLKTPAQFD
       VLALLKTPAQFDA
       VLALLKTPAQFDAD
       VLALLKTPAQFDADE
       VLALLKTPAQFDADEL
       VLALLKTPAQFDADELR
       VLALLKTPAQFDADELRA
       VLALLKTPAQFDADELRAA
       VLALLKTPAQFDADELRAAM
       VLALLKTPAQFDADELRAAMK
       VLALLKTPAQFDADELRAAMKG
        LALLKTPAQFDADEL
        LALLKTPAQFDADELRA
        LALLKTPAQFDADELRAA
         ALLKTPAQFDADEL
         ALLKTPAQFDADELRA
         ALLKTPAQFDADELRAA
         ALLKTPAQFDADELRAAM
          LLKTPAQFDADEL
          LLKTPAQFDADELRA
          LLKTPAQFDADELRAA
           LKTPAQFDADELRA
            KTPAQFDADEL
            KTPAQFDADELRA
            KTPAQFDADELRAA
            KTPAQFDADELRAAM
                QFDADELRA
                QFDADELRAA
                       RAAMKGLGTDEDTLIEI
                       RAAMKGLGTDEDTLIEIL
                       RAAMKGLGTDEDTLIEILA
                       RAAMKGLGTDEDTLIEILASRT
                         AMKGLGTDEDTLIE
                         AMKGLGTDEDTLIEI
```

```
                         AMKGLGTDEDTLIEIL
                         AMKGLGTDEDTLIEILA
                         AMKGLGTDEDTLIEILASRT
                          MKGLGTDEDTLIEI
                          MKGLGTDEDTLIEIL
                          MKGLGTDEDTLIEILASRT
                           KGLGTDEDTLIEIL
                           KGLGTDEDTLIEILA
                                        LASRTNKEIR
                                         ASRTNKEIR
                                         ASRTNKEIR
                                         ASRTNKEIR
                                             NKEIR
                                             NKEIR
                                             NKEIR

151      160       170       180       190       200
DINRVYREELKRDLAKDITSDTSGDFRNALLSLAKGDRSEDFGVNEDLAD
DINRVYREEL
DI
DINRV
DINRVYREEL
DINRV
DINRVYREEL
DINRVYREELKRDLAKDIT
     YREELKRDLAKDITSDT
     YREELKRDLAKDITSDTSGDFRNA
          KRDLAKDITSDT
          KRDLAKDITSDTSGDFRNALL
          KRDLAKDITSDTSGDFRNALLSLA
              AKDITSDTSGDFRNALL
               KDITSDTSGDFRNALL
                  TSDTSGDFRNA
                  TSDTSGDFRNALL
                   SDTSGDFRNA
                   SDTSGDFRNALL
                   SDTSGDFRNALLSL
                   SDTSGDFRNALLSLA
                      SGDFRNALLSLA
                               SLAKGDRSEDFGV
                               SLAKGDRSEDFGVNEDLA
                               SLAKGDRSEDFGVNEDLAD
                               SLAKGDRSEDFGVNEDLAD
                               SLAKGDRSEDFGVNEDLAD
                                LAKGDRSEDFGVNEDLAD
                                  KGDRSEDFGVNEDLAD
                                  KGDRSEDFGVNEDLAD

201      210       220       230       240       250
SDARALYEAGERRKGTDVNVFNTILTTRSYPQLRRVFQKYTKYSKHDMNK
SDA
SDARA
SDARAL
SDARA
SDARA
SDARALYEA
     LYEAGERRKGTDVNVFNT
         GERRKGTDVNVFNT
         GERRKGTDVNVFNTILT
                  NVFNTILTTRSYPQLRRV
                   VFNTILTTRSYPQLRRV
                     NTILTTRSYPQLRRV
                       ILTTRSYPQL
```

```
                       ILTTRSYPQLRRV
                          TRSYPQLRRV
                          TRSYPQLRRVFQ
                                    FQKYTKYSKHDM
                                    FQKYTKYSKHDMNK
                                    FQKYTKYSKHDMNK
                                    FQKYTKYSKHDMNK
                                    FQKYTKYSKHDMNK
                                      KYTKYSKHDMNK
                                         KYSKHDMNK
                                         KYSKHDMNK
                                         KYSKHDMNK
                                            KHDMNK

251      260       270       280       290      300
VLDLELKGDIEKCLTAIVKCATSKPAFFAEKLHQAMKGVGTRHKALIRIM
V
VL
VLDL
VLDLEL
VLDLELKGDIEKC
VL
VLDL
VLDLELKGDIEKCLT
VLDLELKGDIEKC
    ELKGDIEKC
    ELKGDIEKCLTAIVKC
      KGDIEKCLTAIVKC
             LTAIVKCA
             LTAIVKCATSKPA
             LTAIVKCATSKPAFFAEKL
               AIVKCATSKPA
               AIVKCATSKPAFFAEKL
                 VKCATSKPAFFAEKL
                  KCATSKPAFFAEKL
                    ATSKPAFFAEKL
                    ATSKPAFFAEKLH
                    ATSKPAFFAEKLHQ
                    ATSKPAFFAEKLHQA
                    ATSKPAFFAEKLHQAM
                    ATSKPAFFAEKLHQAMKGV
                    ATSKPAFFAEKLHQAMKGVGT
                    ATSKPAFFAEKLHQAMKGVGTRHKA
                     TSKPAFFAEKL
                     TSKPAFFAEKLHQ
                     TSKPAFFAEKLHQAMKGV
                      SKPAFFAEKL
                        PAFFAEKLHQA
                          FFAEKLHQ
                          FFAEKLHQA
                          FFAEKLHQAM
                          FFAEKLHQAMKGV
                          FFAEKLHQAMKGVGTRHKALIRI
                          FFAEKLHQAMKGVGTRHKALIRIM
                          FFAEKLHQAMKGVGTRHKALIRIM
                                HQAMKGVGTRHKAL
                                HQAMKGVGTRHKALI
                                HQAMKGVGTRHKALIRI
                                HQAMKGVGTRHKALIRIM
                                HQAMKGVGTRHKALIRIM
                                HQAMKGVGTRHKALIRIM
                                  AMKGVGTRHKALIRIM
                                   MKGVGTRHKALIRI
```

```
                                   MKGVGTRHKALIRIM
                                   MKGVGTRHKALIRIM
                                   MKGVGTRHKALIRIM
                                     GVGTRHKALIRIM
                                       GTRHKALIRIM
                                              IRIM
                                               RIM
                                                 M
                                                 M
                                                 M
                                                 M
                                                 M

301      310       320       330       340   346
VSRSEIDMNDIKAFYQKMYGISLCQAILDETKGDYEKILVALCGGN
V
V
VS
V
VS
VSRSEIDMNDIKA
VSRSEIDMNDIKA
VSRSEIDMNDI
VSRSEIDMNDIKA
VSRSEIDMNDIKAFYQ
VSRSEIDMNDIKAFYQKM
VSRSEIDMNDIKAFYQKMYGISLC
VSRSEIDMNDI
VSRSEIDMNDIKA
VSRSEIDMNDIKAFYQ
VSRSEIDMNDIKAFYQKM
VSRSEIDMNDIKAFYQKMYGI
VSRSEIDMNDIKAFYQKMYGISLC
 SRSEIDMNDIKA
 SRSEIDMNDIKAFYQ
 SRSEIDMNDIKAFYQKM
 SRSEIDMNDIKAFYQKMYGISLC
  RSEIDMNDIKA
  RSEIDMNDIKAFYQ
    EIDMNDIKA
             FYQKMYGI
             FYQKMYGISLC
             FYQKMYGISLCQA
                KMYGISLC
                KMYGISLCQA
                        QAILDETKGDYEKI
                        QAILDETKGDYEKIL
                        QAILDETKGDYEKILV
                        QAILDETKGDYEKILVA
                        QAILDETKGDYEKILVALC
                        QAILDETKGDYEKILVALCGGN
                          ILDETKGDYEKIL
                          ILDETKGDYEKILV
                          ILDETKGDYEKILVA
                          ILDETKGDYEKILVAL
                          ILDETKGDYEKILVALC
                          ILDETKGDYEKILVALCGGN
                           LDETKGDYEKILV
                           LDETKGDYEKILVALCGGN
                            DETKGDYEKILV
                               KGDYEKILVALC
                               KGDYEKILVALCGGN
```

### **Supplementary data 2**

CLUSTAL O(1.2.4) multiple sequence alignment

```
                                1         10        20        30        40        50        60
P14950|Columba livia p35        MAVVSEFLKQAWFMENLEQECIKCTQCVHGVP-----QQTNFDPSADVVALEKAMTAKGV        55
Q92040|Columba livia p37        MAMVSEFLKQAWFMEHQEQEYIKSVKGGPVVPQ----QQPNFDPSADVVALEKAMTAKGV        56
P24551|Rodentia sp.             MAMVSEFINQACYLEKQEQEYIEIVKSYKGGPAHAVSPYPSFDPSSDVAALHKGIMVNGV        60
P07150|Rattus norvegicus        MAMVSEFLKQACYIEKQEQEYVQAVKSYKGGPGSAVSPYPSFNPSSDVAALHKAIMVKGV        60
P10107|Mus musculus             MAMVSEFLKQARFLENQEQEYVQAVKSYKGGPGSAVSPYPSFNVSSDVAALHKAIMVKGV        60
Q8HZM6|Equus caballus           MSMVSAFLKQAWFIENEEQEYIKAVKGSKGGPGSAVSPYPSFNPSSDVDALHKAITVKGV        60
P19619|Sus scrofa               MAMVSEFLKQAWFIDNEEQEYIKTVKGSKGGPGSAVSPYPTFNPSSDVEASHKAITVKGV        60
P46193|Bos taurus               MAMVSEFLKQAWFIENEEQEYIKTVKGSKGGPGSAVSPYPTFNPSSDVEALHKAITVKGV        60
P14087|Cavia cutleri            MSMVSEFLKQAYFIDNQEQDYVKTVKSSKGGPGSAVSPYPSFDASSDVAALHKAITVKGV        60
P51662|Oryctolagus cuniculus    MAMVSEFLKQAWFIDNEEQDYINTVKTYKGGPGSAVSPYPAFNPSSDVAALHQAIMVKGV        60
Q5REL2|Pongo abelii             MAMVSEFLKQAWFIENEEQEYVQTVKSSKGGPGSAVSPYPTFNPSSDVAALHKAIMVKGV        60
A5A6M2|Pan troglodytes          MAMVSEFLKQAWFIENEEQEYVQTVKSSKGGPGSAVSPYPTFNPSSDVAALHKAIMVKGV        60
P04083|Homo sapiens             MAMVSEFLKQAWFIENEEQEYVQTVKSSKGGPGSAVSPYPTFNPSSDVAALHKAIMVKGV        60
                                *::** *::** :::: **: :: .:     *         *: *:** * .:.: .:**

                                61        70        80        90        100       110       120
P14950|Columba livia p35        DEATIIDIMTTRTNAQRPRIKAAYHKAKGKSLEEAMKRVLKSHLEDVVVALLKTPAQFDA        115
Q92040|Columba livia p37        DEATIIDIMTKRTNAQRHRIKAAYQKAKGKSLEEAMKRVLKSHLEDVVVALLKTPAQFDA        116
P24551|Rodentia sp.             DEATILDLLTKRYNAQRHHLKAVYIQETGEPLDETLKKALTGHIQELLLAMIKTPAQFDG        120
P07150|Rattus norvegicus        DEATIIDILTKRTNAQRQQIKAAYLQETGKPLDETLKKALTGHLEEVVLAMLKTPAQFDA        120
P10107|Mus musculus             DEATIIDILTKRTNAQRQQIKAAYLQENGKPLDEVLRKALTGHLEEVVLAMLKTPAQFDA        120
Q8HZM6|Equus caballus           DEATIIEILTKRNNAQRQQIKAAYLQEKGKPLDEALKKALTGHLEDVALALLKTPARFDA        120
P19619|Sus scrofa               DEATIIEIHTKRTNAQRQQIKAAYLQEKGKPLDEALKKALTGHLEEVALALLKTPAQFDA        120
P46193|Bos taurus               DEATIIEILTKRNNAQRQQIKAAYLQEKGKPLDEVLKKALLGHLEEVVLALLKTPAQFDA        120
P14087|Cavia cutleri            DEATIIDILTKRNNAQRQQIKAAYLQEKGKPLDEALKKALTGHLEEVVLALLKTPAQLDA        120
P51662|Oryctolagus cuniculus    DEATIIDILTKRNNAQRQQIKAAYLQEKGKPLDEVLKKALTGHLEEVVLALLKTPAQFDA        120
Q5REL2|Pongo abelii             DEATIIDVLTKRNNAQRQQIKAAYLQETGKPLDETLKKALTGHLEEVVLALLKTPAQFDA        120
A5A6M2|Pan troglodytes          DEATIIDILTRRNNAQRQQIKAAYLQETGKPLDETLKKALTGHLEEVVLALLKTPAQFDA        120
P04083|Homo sapiens             DEATIIDILTKRNNAQRQQIKAAYLQETGKPLDETLKKALTGHLEEVVLALLKTPAQFDA        120
                                *****::: * * **** ::**.* : .*: *:*.:::.* .*:::: :*::****::*.

                                121       130       140       150       160       170       180
P14950|Columba livia p35        EELRACMKGHGTDEDTLIEILASRNNKEIREACRYYKEVLKRDLTQDIISDTSGDFQKAL        175
Q92040|Columba livia p37        EELRACMKGLGTDEDTLIEILASRSNKEIREASRYYKEVLKRDLTQDIISDTSGHFQKAL        176
P24551|Rodentia sp.             NELRAAMKAVGTDEETLIEILWTRSNQQIREITSVYREELKKDIAKYQTSDTSGEFRDAL        180
P07150|Rattus norvegicus        DELRAAMKGLGTDEDTLIEILTTRSNQQIREITRVYREELKRDLAKDITSDTSGDFRNAL        180
P10107|Mus musculus             DELRGAMKGLGTDEDTLIEILTTRSNEQIREINRVYREELKRDLAKDITSDTSGDFRKAL        180
Q8HZM6|Equus caballus           DELRAAMKGLGTDEDTLIEILTSRTNKEIREINRVYREELKRDLAKDITSDTSGDFQKAL        180
P19619|Sus scrofa               DELRAAMKGLGTDEDTLNEILASRTNREIREINRVYKEELKRDLAKDITSDTSGDYQKAL        180
P46193|Bos taurus               EELRAAMKGLGTDEDTLNEILASRTNREIREINRVYREELKRDLAKDIASDTSGDYEKAL        180
P14087|Cavia cutleri            DELRAAMKGLGTDEDTLIEILVSRKNREIKEINRVYRDELKRDLAKDITSDTSGDFQKAL        180
P51662|Oryctolagus cuniculus    DELRAAMKGLGTDEDTLIEILASRNNKEIREINRVYREELKRDLAKDIASDTSGDFQKAL        180
Q5REL2|Pongo abelii             DELRAAMKGLGTDEDTLIEILASRTNKEIRDINRVYREELKRDLAKDITSDTSGDFRNAL        180
A5A6M2|Pan troglodytes          DELRAAMKGLGTDEDTLIEILASRTNKEIRDINRVYREELKRDLAKDITSDTSGDFRNAL        180
P04083|Homo sapiens             DELRAAMKGLGTDEDTLIEILASRTNKEIRDINRVYREELKRDLAKDITSDTSGDFRNAL        180
                                :***..**. ****:** *** :*.*.:*::    *:: **:*:::   *****.:..**

                                181       190       200       210       220       230       240
P14950|Columba livia p35        VSLAKADRCENPHVNDELAEKDARALYEAGEQKKGTDINVFVTVLTARSYPHS-EVFQKY        234
Q92040|Columba livia p37        VVLAKGDRCEDPHVNDDLADNDARALYEAGEQKKGTDVNVFVTVLTARSYPHLRRVFQKY        236
P24551|Rodentia sp.             LALAKGNRCEDMSVNQDIADTDARALYQAAERRNGTDVNVFNTILTTKKYPHLRNKFQNY        240
P07150|Rattus norvegicus        LALAKGDRCEDMSVNQDLADTDARALYEAGERRKGTDVNVFNTILTTRSYPHLRKVFQNY        240
P10107|Mus musculus             LALAKGDRCQDLSVNQDLADTDARALYEAGERRKGTDVNVFTTILTSRSFPHLRRVFQNY        240
Q8HZM6|Equus caballus           LSLAKGDRSEDFGVNDDLADSDARALYEAGERRKGTDVNVFNTILTTRSYPHLRRVFQMY        240
P19619|Sus scrofa               LSLAKGDRSEDLAINDDLADTDARALYEAGERRKGTDLNVFITILTTRSYLHLRRVFQKY        240
P46193|Bos taurus               LSLAKGDRSEELAVNDDLADSDARALYEAGERRKGTDVNVFITILTTRSYPHLRRVFQKY        240
P14087|Cavia cutleri            LSLAKGDRCEDLSVNDDLADSDARALYEAGERRKGTDVNVFITILTTRSYSHLRRVFQKY        240
P51662|Oryctolagus cuniculus    LSLAKGDRSEDFGVNEDLADTDARALYEAGERRKGADVNVFTTILTTRSYLHLRRVFQKY        240
Q5REL2|Pongo abelii             LSLAKGDRSEDFGVNEDLADSDARALYEAGERRKGTDVNVFNTILTTRSYPQLRRVFQKY        240
A5A6M2|Pan troglodytes          LSLAKGDRSEDFGVNEDLADSDARALYEAGERRKGTDVNVFNTILTTRSYPQLRRVFQKY        240
P04083|Homo sapiens             LSLAKGDRSEDFGVNEDLADSDARALYEAGERRKGTDVNVFNTILTTRSYPQLRRVFQKY        240
                                : ***.:*.::  :*:::*:.******:*.*:::*:*:*** *:**::.: :  . ** *

                                241       250       260       270       280       290       300
P14950|Columba livia p35        TKYSKHDMNKAVDMEMKGDIEKCLTALVKCATSKPAFFAEKLHMAMKGFGTQHRDLIRIM        294
Q92040|Columba livia p37        TKYSKHDMNKVVDMELKGDIEKCLTALVKCATSKPAFFAEKLHLAMKGFGTRHKDLIRIM        296
P24551|Rodentia sp.             RKYTEEDMKKALDIELKGQIEKCLTTIAKCGTSTPAFFAEKLYEAMKGAGTRHKTLIRIM        300
P07150|Rattus norvegicus        RKYSQHDMNKALDLELKGDIEKCLTTIVKCATSTPAFFAEKLYEAMKGAGTRHKTLIRIM        300
P10107|Mus musculus             GKYSQHDMNKALDLELKGDIEKCLTTIVKCATSTPAFFAEKLYEAMKGAGTRHKALIRIM        300
Q8HZM6|Equus caballus           TKYSKHDMNKVLDLEMKGDVENCFTAIVKCATSKPMFFAEKLHNAMKGAGTRDKILIRIM        300
P19619|Sus scrofa               SKYSKHDMNKVLDLELKGDIENCLTVVVKCATSKPMFFAEKLHQAMKGNGTRHKTLIRIM        300
P46193|Bos taurus               SKYSKHDMNKVLDLELKGDIEKCLTVIVKCATSQPMFFAEKLHQAMKGIGTRHKTLIRIM        300
P14087|Cavia cutleri            TKYSQHDMNKALDLELKGDIENCLTAIVKCATSTPAFFAEKLHLAMKGAGTRHKALIRIM        300
P51662|Oryctolagus cuniculus    SKYSQHDMNKVLDLELKGDIEKCLTAIVQCATCKPAYFAEKLYQAMKGAGTRHKALIRIM        300
Q5REL2|Pongo abelii             TKYSKHDMNKVLDLELKGDIEKCLTAIVKCATSKPAFFAEKLHQAMKGVGTRHKALIRIM        300
A5A6M2|Pan troglodytes          TKYSKHDMNKVLDLELKGDIEKCLTAIVKCATSKPAFFAEKLHQAMKGVGTRHKALIRIM        300
P04083|Homo sapiens             TKYSKHDMNKVLDLELKGDIEKCLTAIVKCATSKPAFFAEKLHQAMKGVGTRHKALIRIM        300
                                 **::.**:*.:*:*:**::*:*:*.:.:*.*. * :*****: **** **:.: *****

                                301       310       320       330       340
P14950|Columba livia p35        VSRHEVDMNEIKGYYKKMYGISLCQAIMDELKGGYETILVALCGSDN                     341
Q92040|Columba livia p37        VSRHEVDMNEIKCYYKKMYGISLCQAIMDDLKGDYETILVALCGSDN                     343
P24551|Rodentia sp.             VSRSEIDSDQIKVFYQKKYGVPLCQAILDETKGAYEKILVALEGGN-                     346
P07150|Rattus norvegicus        VSRSEIDMNEIKVFYQKKYGIPLCQAILDETKGDYEKILVALCGGN-                     346
P10107|Mus musculus             VSRSEIDMNEIKVFYQKKYGISLCQAILDETKGDYEKILVALCGGN-                     346
Q8HZM6|Equus caballus           VSRSEVDMNDIKACYQKLYGISLCQAILDETKGDYEKILVALCGRD-                     346
P19619|Sus scrofa               VSRSEIDMNDIKACYQKLYGISLCQAILDETKGDYEKILVALCGGD-                     346
P46193|Bos taurus               VSRSEIDMNDIKACYQKLYGISLCQAILDETKGDYEKILVALCGRD-                     346
P14087|Cavia cutleri            VSRSEIDMNDIKVYYQKMYGISLCQAILDETKGDYEKILVALCGGQ-                     346
P51662|Oryctolagus cuniculus    VSRSEVDMNDIKAFYQKKYGVSLCQAILDETKGDYEKILVALCGGN-                     346
Q5REL2|Pongo abelii             VSRSEIDMNDIKAFYQKMYGISLCQAILDETKGDYEKILVALCGGN-                     346
A5A6M2|Pan troglodytes          VSRSEIDMNDIKAFYQKMYGISLCQAILDETKGDYEKILVALCGGN-                     346
P04083|Homo sapiens             VSRSEIDMNDIKAFYQKMYGISLCQAILDETKGDYEKILVALCGGN-                     346
                                *** *:* ::**  *:* **: *****:*: ** **.***** * :
```

### Supplementary data 3

**Folded ANXA1 0 sec**

```
1        10        20        30        40        50
MAMVSEFLKQAWFIENEEQEYVQTVKSSKGGPGSAVSPYPTFNPSSDVAA
           WFIENEEQEYVQTVKSSKGGPGSA
                           SKGGPGSAVSPYPTFNPSSDVA

51       60        70        80        90        100
LHKAIMVKGVDEATIIDILTKRNNAQRQQIKAAYLQETGKPLDETLKKAL

101      110       120       130       140       150
TGHLEEVVLALLKTPAQFDADELRAAMKGLGTDEDTLIEILASRTNKEIR

151      160       170       180       190       200
DINRVYREELKRDLAKDITSDTSGDFRNALLSLAKGDRSEDFGVNEDLAD

201      210       220       230       240       250
SDARALYEAGERRKGTDVNVFNTILTTRSYPQLRRVFQKYTKYSKHDMNK

251      260       270       280       290       300
VLDLELKGDIEKCLTAIVKCATSKPAFFAEKLHQAMKGVGTRHKALIRIM

301      310       320       330       340   346
VSRSEIDMNDIKAFYQKMYGISLCQAILDETKGDYEKILVALCGGN
```

**Folded ANXA1 15 sec**

```
1        10        20        30        40        50
MAMVSEFLKQAWFIENEEQEYVQTVKSSKGGPGSAVSPYPTFNPSSDVAA
           WFIENEEQEYVQTV
                   EYVQTVKSSKGGPGSAVSPYPT
                        VKSSKGGPGSAVSPYPT
                         KSSKGGPGSAVSPYPT
                         KSSKGGPGSAVSPYPTFNPSSD
                         KSSKGGPGSAVSPYPTFNPSSDVA
                         KSSKGGPGSAVSPYPTFNPSSDVAA
                           SKGGPGSAVSPYPT
                            KGGPGSAVSPYPT
                                         FNPSSDVAA
                                         FNPSSDVAA
                                         FNPSSDVAA
                                         FNPSSDVAA

51       60        70        80        90        100
LHKAIMVKGVDEATIIDILTKRNNAQRQQIKAAYLQETGKPLDETLKKAL
L
LHKA
LHKAIMVK
LHKAIMVKGVDEAT
    IMVKGVDEATIIDI
                                AYLQETGKP
                                AYLQETGKPL
                                AYLQETGKPLD
                                AYLQETGKPLDE
                                AYLQETGKPLDET
                                AYLQETGKPLDETL
                                AYLQETGKPLDETLK
                                AYLQETGKPLDETLKKAL
                                AYLQETGKPLDETLKKAL
                                AYLQETGKPLDETLKKAL
                                   QETGKPLDETLKKAL

101      110       120       130       140       150
TGHLEEVVLALLKTPAQFDADELRAAMKGLGTDEDTLIEILASRTNKEIR
TG
TGHLEEV
TGHLEEV
       VLALLKTPAQFDADEL
       VLALLKTPAQFDADELRA
       VLALLKTPAQFDADELRAA
                         AMKGLGTDEDTLIEILA

151      160       170       180       190       200
DINRVYREELKRDLAKDITSDTSGDFRNALLSLAKGDRSEDFGVNEDLAD

201      210       220       230       240       250
SDARALYEAGERRKGTDVNVFNTILTTRSYPQLRRVFQKYTKYSKHDMNK

251      260       270       280       290       300
VLDLELKGDIEKCLTAIVKCATSKPAFFAEKLHQAMKGVGTRHKALIRIM

301      310       320       330       340   346
VSRSEIDMNDIKAFYQKMYGISLCQAILDETKGDYEKILVALCGGN
 SRSEIDMNDIKAFYQKMYGISLC
```

**Folded ANXA1 30 sec**

```
1         10        20        30        40        50
MAMVSEFLKQAWFIENEEQEYVQTVKSSKGGPGSAVSPYPTFNPSSDVAA
 AMVSEFLKQA
          AWFIENEEQEYVQTV
           WFIENEEQEYVQTV
                   EYVQTVKSSKGGPGSAVSPYPT
                        VKSSKGGPGSAVSPYPT
                         KSSKGGPGSAVSPYPT
                         KSSKGGPGSAVSPYPTFNPSSD
                         KSSKGGPGSAVSPYPTFNPSSDVA
                         KSSKGGPGSAVSPYPTFNPSSDVAA
                           SKGGPGSAVSPYPT
                            KGGPGSAVSPYPT
                                         FNPSSDVAA
                                         FNPSSDVAA
                                         FNPSSDVAA
                                         FNPSSDVAA
                                         FNPSSDVAA
                                         FNPSSDVAA

51        60        70        80        90        100
LHKAIMVKGVDEATIIDILTKRNNAQRQQIKAAYLQETGKPLDETLKKAL
L
LHKA
LHKAIMV
LHKAIMVK
LHKAIMVKGVDEA
LHKAIMVKGVDEAT
    IMVKGVDEA
    IMVKGVDEAT
    IMVKGVDEATI
    IMVKGVDEATII
    IMVKGVDEATIIDILTK
    IMVKGVDEATIIDILTKRN
                                AYLQETGKP
                                AYLQETGKPL
                                AYLQETGKPLD
                                AYLQETGKPLDE
                                AYLQETGKPLDET
                                AYLQETGKPLDETL
                                AYLQETGKPLDETLK
                                AYLQETGKPLDETLKKA
                                AYLQETGKPLDETLKKAL
                                AYLQETGKPLDETLKKAL
                                AYLQETGKPLDETLKKAL
                                AYLQETGKPLDETLKKAL
                                AYLQETGKPLDETLKKAL
                                 YLQETGKPLDE
                                 YLQETGKPLDETL
                                 YLQETGKPLDETLKKA
                                 YLQETGKPLDETLKKAL
                                   QETGKPLDETLKKAL
                                      GKPLDETLKKAL
                                        PLDETLKKAL
                                                 L

101       110       120       130       140       150
TGHLEEVVLALLKTPAQFDADELRAAMKGLGTDEDTLIEILASRTNKEIR
TG
TGH
TGHLE
```

```
TGHLEEV
TGHLEEV
TGHLEEV
TGHLEEV
TGHLEEV
TGHLEEV
       VLALLKTPA
       VLALLKTPAQFDA
       VLALLKTPAQFDADEL
       VLALLKTPAQFDADELR
       VLALLKTPAQFDADELRA
       VLALLKTPAQFDADELRAA
       VLALLKTPAQFDADELRAAM
                         AMKGLGTDEDTLIEILA

151      160       170       180       190       200
DINRVYREELKRDLAKDITSDTSGDFRNALLSLAKGDRSEDFGVNEDLAD

201      210       220       230       240       250
SDARALYEAGERRKGTDVNVFNTILTTRSYPQLRRVFQKYTKYSKHDMNK

251      260       270       280       290       300
VLDLELKGDIEKCLTAIVKCATSKPAFFAEKLHQAMKGVGTRHKALIRIM
                    ATSKPAFFAEKL

301      310       320       330       340   346
VSRSEIDMNDIKAFYQKMYGISLCQAILDETKGDYEKILVALCGGN
             FYQKMYGISLCQA
```

**Folded ANXA1 60 sec**

```
1        10        20        30        40        50
MAMVSEFLKQAWFIENEEQEYVQTVKSSKGGPGSAVSPYPTFNPSSDVAA
 AMVSEFLKQA
     EFLKQAWFIENEEQEYVQTV
          AWFIENEEQEYVQTV
           WFIENEEQ
           WFIENEEQEYVQTV
                   EYVQTVKSSKGGPGSAVSPYPT
                       TVKSSKGGPGSAVSPYPT
                        VKSSKGGPGSAVSPYPT
                        VKSSKGGPGSAVSPYPTFNPSSDVA
                         KSSKGGPGSAVSPYPT
                         KSSKGGPGSAVSPYPTFNPSSD
                         KSSKGGPGSAVSPYPTFNPSSDV
                         KSSKGGPGSAVSPYPTFNPSSDVA
                         KSSKGGPGSAVSPYPTFNPSSDVAA
                           SKGGPGSAVSPYPT
                            KGGPGSAVSPYPT
                                         FNPSSDVAA
                                         FNPSSDVAA
                                         FNPSSDVAA
                                         FNPSSDVAA
                                         FNPSSDVAA
                                         FNPSSDVAA
                                           PSSDVAA
51       60        70        80        90        100
LHKAIMVKGVDEATIIDILTKRNNAQRQQIKAAYLQETGKPLDETLKKAL
L
LHKA
LHKAIMV
LHKAIMVK
LHKAIMVKGVDEA
LHKAIMVKGVDEAT
LHKA
    IMVKGVDEA
    IMVKGVDEAT
    IMVKGVDEATI
    IMVKGVDEATII
    IMVKGVDEATIIDIL
    IMVKGVDEATIIDILT
    IMVKGVDEATIIDILTKRNNAQ
                       NAQRQQIKAAYL
                                AYLQETGKP
                                AYLQETGKPL
                                AYLQETGKPLD
                                AYLQETGKPLDE
                                AYLQETGKPLDET
                                AYLQETGKPLDETL
                                AYLQETGKPLDETLK
                                AYLQETGKPLDETLKKA
                                AYLQETGKPLDETLKKAL
                                AYLQETGKPLDETLKKAL
                                AYLQETGKPLDETLKKAL
                                AYLQETGKPLDETLKKAL
                                AYLQETGKPLDETLKKAL
                                AYLQETGKPLDETLKKAL
                                AYLQETGKPLDETLKKAL
                                 YLQETGKPLDE
                                 YLQETGKPLDET
                                 YLQETGKPLDETL
```

```
                                 YLQETGKPLDETLKKA
                                 YLQETGKPLDETLKKAL
                                   QETGKPLDET
                                   QETGKPLDETLKKAL
                                             LKKAL
                                                 L

101      110       120       130       140       150
TGHLEEVVLALLKTPAQFDADELRAAMKGLGTDEDTLIEILASRTNKEIR
T
TG
TGH
TGHL
TGHLE
TGHLEE
TGHLEEV
TG
TGHLEEV
TGHLEEV
TGHLEEV
TGHLEEV
       VLALLKTP
       VLALLKTPA
       VLALLKTPAQ
       VLALLKTPAQF
       VLALLKTPAQFD
       VLALLKTPAQFDA
       VLALLKTPAQFDAD
       VLALLKTPAQFDADE
       VLALLKTPAQFDADEL
       VLALLKTPAQFDADELR
       VLALLKTPAQFDADELRA
       VLALLKTPAQFDADELRAA
       VLALLKTPAQFDADELRAAM
       VLALLKTPAQFDADELRAAMKGLGT
            KTPAQFDADELRA
            KTPAQFDADELRAA
                       RAAMKGLGTDEDTLIEI
                         AMKGLGTDEDTLIEILA

151      160       170       180       190       200
DINRVYREELKRDLAKDITSDTSGDFRNALLSLAKGDRSEDFGVNEDLAD
                               SLAKGDRSEDFGVNEDLA
```

```
                              SLAKGDRSEDFGVNEDLADSD

201      210       220       230       240       250
SDARALYEAGERRKGTDVNVFNTILTTRSYPQLRRVFQKYTKYSKHDMNK
SDARAL
                       ILTTRSYPQLRRV

251      260       270       280       290       300
VLDLELKGDIEKCLTAIVKCATSKPAFFAEKLHQAMKGVGTRHKALIRIM
                    ATSKPAFFAEKL
                    ATSKPAFFAEKLHQAM
                     TSKPAFFAEKL

301      310       320       330       340   346
VSRSEIDMNDIKAFYQKMYGISLCQAILDETKGDYEKILVALCGGN
                        QAILDETKGDYEKILV
                          ILDETKGDYEKILV
```

**Folded ANXA1 90 sec (5)**

```
1        10        20        30        40        50
MAMVSEFLKQAWFIENEEQEYVQTVKSSKGGPGSAVSPYPTFNPSSDVAA
 AMVSEFLKQ
 AMVSEFLKQA
  MVSEFLKQA
        KQAWFIENEEQEYVQTV
          AWFIENEEQEYVQTV
           WFIENEEQ
           WFIENEEQEYVQTV
           WFIENEEQEYVQTVKS
                   EYVQTVKSSKGGPGSAVSPYPT
                       TVKSSKGGPGSAVSPYPT
                        VKSSKGGPGSAVSPYPT
                        VKSSKGGPGSAVSPYPTFNPSSDVA
                         KSSKGGPGSAVSPYPT
                         KSSKGGPGSAVSPYPTFNPSSD
                         KSSKGGPGSAVSPYPTFNPSSDV
                         KSSKGGPGSAVSPYPTFNPSSDVA
                         KSSKGGPGSAVSPYPTFNPSSDVAA
                           SKGGPGSAVSPYPT
                            KGGPGSAVSPYPT
                                         FNPSSDVAA
                                         FNPSSDVAA
                                         FNPSSDVAA
                                         FNPSSDVAA
                                         FNPSSDVAA
                                         FNPSSDVAA
                                         FNPSSDVAA
                                           PSSDVAA
51       60        70        80        90        100
LHKAIMVKGVDEATIIDILTKRNNAQRQQIKAAYLQETGKPLDETLKKAL
L
LHKA
LHKAIM
LHKAIMV
LHKAIMVK
LHKAIMVKGVDEA
LHKAIMVKGVDEAT
LHKA
    IMVKGVDEA
    IMVKGVDEAT
    IMVKGVDEATI
    IMVKGVDEATII
    IMVKGVDEATIIDI
    IMVKGVDEATIIDIL
    IMVKGVDEATIIDILT
    IMVKGVDEATIIDILTK
       KGVDEATIIDILTKRNNAQRQQIKA
                   TKRNNAQRQQIKAAYLQETGKPLDE
                       NAQRQQIKAAYL
                                AYLQETGKP
                                AYLQETGKPL
                                AYLQETGKPLD
                                AYLQETGKPLDE
                                AYLQETGKPLDET
                                AYLQETGKPLDETL
                                AYLQETGKPLDETLK
                                AYLQETGKPLDETLKKA
                                AYLQETGKPLDETLKKAL
                                AYLQETGKPLDETLKKAL
```

```
                                AYLQETGKPLDETLKKAL
                                AYLQETGKPLDETLKKAL
                                AYLQETGKPLDETLKKAL
                                AYLQETGKPLDETLKKAL
                                AYLQETGKPLDETLKKAL
                                 YLQETGKPLD
                                 YLQETGKPLDE
                                 YLQETGKPLDET
                                 YLQETGKPLDETL
                                 YLQETGKPLDETLKKA
                                 YLQETGKPLDETLKKAL
                                   QETGKPLDET
                                   QETGKPLDETLKKAL
                                             LKKAL
                                                 L

101      110       120       130       140       150
TGHLEEVVLALLKTPAQFDADELRAAMKGLGTDEDTLIEIILASRTNKEIR
T
TG
TGH
TGHL
TGHLE
TGHLEE
TGHLEEV
TG
TGHLEEV
TGHLEEV
TGHLEEV
TGHLEEV
TGHLEEV
       VLALLKTP
       VLALLKTPA
       VLALLKTPAQ
       VLALLKTPAQF
       VLALLKTPAQFD
       VLALLKTPAQFDA
       VLALLKTPAQFDAD
       VLALLKTPAQFDADE
       VLALLKTPAQFDADEL
       VLALLKTPAQFDADELR
       VLALLKTPAQFDADELRA
       VLALLKTPAQFDADELRAA
       VLALLKTPAQFDADELRAAM
       VLALLKTPAQFDADELRAAMKGLGT
         ALLKTPAQFDADEL
          LLKTPAQFDADEL
            KTPAQFDADELRA
            KTPAQFDADELRAA
                       RAAMKGLGTDEDTLIEI
                       RAAMKGLGTDEDTLIEIL
                         AMKGLGTDEDTLIEI
                         AMKGLGTDEDTLIEILA

151      160       170       180       190       200
DINRVYREELKRDLAKDITSDTSGDFRNALLSLAKGDRSEDFGVNEDLAD
                               SLAKGDRSEDFGVNEDLA
```

```
                              SLAKGDRSEDFGVNEDLADSD
                              SLAKGDRSEDFGVNEDLADSD

201      210       220       230       240       250
SDARALYEAGERRKGTDVNVFNTILTTRSYPQLRRVFQKYTKYSKHDMNK
SDARA
SDARAL
                       ILTTRSYPQLRRV
                          TRSYPQLRRV
                                    FQKYTKYSKHDMNK
                                    FQKYTKYSKHDMNK

251      260       270       280       290       300
VLDLELKGDIEKCLTAIVKCATSKPAFFAEKLHQAMKGVGTRHKALIRIM
VLDL
VLDLEL
                    ATSKPAFFAEKL
                    ATSKPAFFAEKLHQ
                    ATSKPAFFAEKLHQA
                     TSKPAFFAEKL
                      SKPAFFAEKL
                           FAEKLHQA
                                HQAMKGVGTRHKALIRIM

301      310       320       330       340   346
VSRSEIDMNDIKAFYQKMYGISLCQAILDETKGDYEKILVALCGGN
VSRSEIDMNDIKA
VSRSEIDMNDIKAFYQKM
             FYQKMYGISLCQA
                        QAILDETKGDYEKILV
                          ILDETKGDYEKILV
                          ILDETKGDYEKILVALC
```

**Folded ANXA1 120 sec (6)**

```
1        10        20        30        40        50
MAMVSEFLKQAWFIENEEQEYVQTVKSSKGGPGSAVSPYPTFNPSSDVAA
 AMVSEFLKQ
 AMVSEFLKQA
  MVSEFLKQA
        KQAWFIENEEQEYVQTV
          AWFIENEEQEYVQTV
           WFIENEEQ
           WFIENEEQEYVQT
           WFIENEEQEYVQTV
           WFIENEEQEYVQTVKS
           WFIENEEQEYVQTVKSS
              ENEEQEYVQTV
                   EYVQTVKSSKGGPGSAVSPYPT
                       TVKSSKGGPGSAVSPYPT
                        VKSSKGGPGSAVSPYPT
                        VKSSKGGPGSAVSPYPTFNPSSD
                        VKSSKGGPGSAVSPYPTFNPSSDVA
                         KSSKGGPGSAVSPYPT
                         KSSKGGPGSAVSPYPTFNPSS
                         KSSKGGPGSAVSPYPTFNPSSD
                         KSSKGGPGSAVSPYPTFNPSSDV
                         KSSKGGPGSAVSPYPTFNPSSDVA
                         KSSKGGPGSAVSPYPTFNPSSDVAA
                           SKGGPGSAVSPYPT
                            KGGPGSAVSPYPT
                                       PTFNPSSDVAA
                                         FNPSSDVAA
                                         FNPSSDVAA
                                         FNPSSDVAA
                                         FNPSSDVAA
                                         FNPSSDVAA
                                         FNPSSDVAA
                                         FNPSSDVAA
                                         FNPSSDVAA
                                          NPSSDVAA
                                           PSSDVAA

51       60        70        80        90        100
LHKAIMVKGVDEATIIDILTKRNNAQRQQIKAAYLQETGKPLDETLKKAL
LHKA
L
LHKA
LHKAIM
LHKAIMV
LHKAIMVK
LHKAIMVKGVDEA
LHKAIMVKGVDEAT
LHKA
LHKA
    IMVKGVDEA
    IMVKGVDEATI
    IMVKGVDEATIIDI
    IMVKGVDEATIIDIL
    IMVKGVDEATIIDILT
    IMVKGVDEATIIDILTK
    IMVKGVDEATIIDILTKR
    IMVKGVDEATIIDILTKRN
    IMVKGVDEATIIDILTKRNNAQ
       KGVDEATIIDILTKRNNAQRQQIKA
```

```
                   TKRNNAQRQQIKAAYLQET
                   TKRNNAQRQQIKAAYLQETGKPLDE
                       NAQRQQIKAAYL
                       NAQRQQIKAAYLQET
                              KAAYLQET
                                AYLQETGKP
                                AYLQETGKPL
                                AYLQETGKPLD
                                AYLQETGKPLDE
                                AYLQETGKPLDET
                                AYLQETGKPLDETL
                                AYLQETGKPLDETLK
                                AYLQETGKPLDETLKK
                                AYLQETGKPLDETLKKA
                                AYLQETGKPLDETLKKAL
                                AYLQETGKPLDETLKKAL
                                AYLQETGKPLDETLKKAL
                                AYLQETGKPLDETLKKAL
                                AYLQETGKPLDETLKKAL
                                AYLQETGKPLDETLKKAL
                                AYLQETGKPLDETLKKAL
                                AYLQETGKPLDETLKKAL
                                 YLQETGKPLD
                                 YLQETGKPLDE
                                 YLQETGKPLDET
                                 YLQETGKPLDETL
                                 YLQETGKPLDETLKKA
                                 YLQETGKPLDETLKKAL
                                 YLQETGKPLDETLKKAL
                                   QETGKPLDET
                                   QETGKPLDETLKKAL
                                      GKPLDETLKKAL
                                        PLDETLKKAL
                                             LKKAL
                                                 L
                                                 L
101      110       120       130       140       150
TGHLEEVVLALLKTPAQFDADELRAAMKGLGTDEDTLIEILASRTNKEIR
T
TG
TGH
TGHL
TGHLE
TGHLEE
TGHLEEV
TG
TGHLEEV
TGHLEEV
TGHLEEV
TGHLEEV
TGHLEEV
TGHLEEV
TGHLEEVVLA
       VLALLKTP
       VLALLKTPA
       VLALLKTPAQ
       VLALLKTPAQF
       VLALLKTPAQFD
       VLALLKTPAQFDA
       VLALLKTPAQFDAD
       VLALLKTPAQFDADE
       VLALLKTPAQFDADEL
       VLALLKTPAQFDADELR
```

```
       VLALLKTPAQFDADELRA
       VLALLKTPAQFDADELRAA
       VLALLKTPAQFDADELRAAM
       VLALLKTPAQFDADELRAAMKG
       VLALLKTPAQFDADELRAAMKGLGT
         ALLKTPAQFDADEL
         ALLKTPAQFDADELRA
         ALLKTPAQFDADELRAA
          LLKTPAQFDADEL
          LLKTPAQFDADELRA
            KTPAQFDADEL
            KTPAQFDADELRA
            KTPAQFDADELRAA
                       RAAMKGLGTDEDTLIEI
                       RAAMKGLGTDEDTLIEIL
                         AMKGLGTDEDTL
                         AMKGLGTDEDTLIEI
                         AMKGLGTDEDTLIEIL
                         AMKGLGTDEDTLIEILA
                         AMKGLGTDEDTLIEILASRT
                           KGLGTDEDTLIEILA

151      160       170       180       190       200
DINRVYREELKRDLAKDITSDTSGDFRNALLSLAKGDRSEDFGVNEDLAD
                   SDTSGDFRNALL
                               SLAKGDRSEDFGV
                               SLAKGDRSEDFGVNEDLA
                               SLAKGDRSEDFGVNEDLAD
                               SLAKGDRSEDFGVNEDLAD

201      210       220       230       240       250
SDARALYEAGERRKGTDVNVFNTILTTRSYPQLRRVFQKYTKYSKHDMNK
SDARA
SDARAL
                   VFNTILTTRSYPQLRRV
                       ILTTRSYPQLRRV
                          TRSYPQLRRV
                                    FQKYTKYSKHDMNK
                                    FQKYTKYSKHDMNK

251      260       270       280       290       300
VLDLELKGDIEKCLTAIVKCATSKPAFFAEKLHQAMKGVGTRHKALIRIM
V
VLDL
                    ATSKPAFFAEKL
                    ATSKPAFFAEKLHQ
                    ATSKPAFFAEKLHQA
                    ATSKPAFFAEKLHQAM
                    ATSKPAFFAEKLHQAMKGV
                     TSKPAFFAEKL
                      SKPAFFAEKL
                          FFAEKLHQA
                          FFAEKLHQAMKGV
                                HQAMKGVGTRHKALIRIM

301      310       320       330       340   346
VSRSEIDMNDIKAFYQKMYGISLCQAILDETKGDYEKILVALCGGN
VSRSEIDMNDIKA
                        QAILDETKGDYEKILV
                        QAILDETKGDYEKILVALC
                        QAILDETKGDYEKILVALCGGN
                          ILDETKGDYEKILV
                          ILDETKGDYEKILVA
```

ILDETKGDYEKILVALC
ILDETKGDYEKILVALCGGN

**Folded ANXA1 300 sec**

```
1        10        20        30        40        50
MAMVSEFLKQAWFIENEEQEYVQTVKSSKGGPGSAVSPYPTFNPSSDVAA
 AMVSEFLKQ
 AMVSEFLKQA
 AMVSEFLKQAWFIENEEQEYVQTV
  MVSEFLKQ
  MVSEFLKQA
        KQAWFIENEEQEYVQTV
          AWFIENEEQ
          AWFIENEEQEYVQTV
           WFIENEEQ
           WFIENEEQEYV
           WFIENEEQEYVQT
           WFIENEEQEYVQTV
           WFIENEEQEYVQTVKS
           WFIENEEQEYVQTVKSS
           WFIENEEQEYVQTVKSSKGGPGSA
              ENEEQEYVQTV
                   EYVQTVKSSKGGPGSAVSPYPT
                       TVKSSKGGPGSAVSPYPT
                        VKSSKGGPGSAVSPYPT
                        VKSSKGGPGSAVSPYPTFNPSSD
                        VKSSKGGPGSAVSPYPTFNPSSDVA
                         KSSKGGPGSAVSPYPT
                         KSSKGGPGSAVSPYPTFNPSS
                         KSSKGGPGSAVSPYPTFNPSSD
                         KSSKGGPGSAVSPYPTFNPSSDV
                         KSSKGGPGSAVSPYPTFNPSSDVA
                         KSSKGGPGSAVSPYPTFNPSSDVAA
                          SSKGGPGSAVSPYPT
                           SKGGPGSAVSPYPT
                            KGGPGSAVSPYPT
                                       PTFNPSSDVAA
                                        TFNPSSDVAA
                                         FNPSSDVAA
                                         FNPSSDVAA
                                         FNPSSDVAA
                                         FNPSSDVAA
                                         FNPSSDVAA
                                         FNPSSDVAA
                                         FNPSSDVAA
                                         FNPSSDVAA
                                          NPSSDVAA
                                           PSSDVAA
                                            SSDVAA
                                                 A

51       60        70        80        90        100
LHKAIMVKGVDEATIIDILTKRNNAQRQQIKAAYLQETGKPLDETLKKAL
LHKA
LHKA
L
LHKA
LHKAIM
LHKAIMV
LHKAIMVK
LHKAIMVKGVDEA
LHKAIMVKGVDEAT
LHKA
```

```
LHKA
LHKA
LHKAIMVKGVDEATIIDIL
    IMVKGVDEA
    IMVKGVDEATI
    IMVKGVDEATIIDI
    IMVKGVDEATIIDIL
    IMVKGVDEATIIDILT
    IMVKGVDEATIIDILTK
    IMVKGVDEATIIDILTKR
    IMVKGVDEATIIDILTKRN
    IMVKGVDEATIIDILTKRNNAQ
      VKGVDEATIIDIL
       KGVDEATIIDIL
       KGVDEATIIDILTKRN
       KGVDEATIIDILTKRNNAQRQQIKA
                   TKRNNAQRQQIKAAYL
                   TKRNNAQRQQIKAAYLQET
                   TKRNNAQRQQIKAAYLQETGKPLDE
                      NNAQRQQIKAAYL
                      NNAQRQQIKAAYLQET
                       NAQRQQIKAAYL
                       NAQRQQIKAAYLQET
                              KAAYLQET
                                AYLQETGKP
                                AYLQETGKPL
                                AYLQETGKPLD
                                AYLQETGKPLDE
                                AYLQETGKPLDET
                                AYLQETGKPLDETL
                                AYLQETGKPLDETLK
                                AYLQETGKPLDETLKKA
                                AYLQETGKPLDETLKKAL
                                AYLQETGKPLDETLKKAL
                                AYLQETGKPLDETLKKAL
                                AYLQETGKPLDETLKKAL
                                AYLQETGKPLDETLKKAL
                                AYLQETGKPLDETLKKAL
                                AYLQETGKPLDETLKKAL
                                AYLQETGKPLDETLKKAL
                                 YLQETGKPLD
                                 YLQETGKPLDE
                                 YLQETGKPLDET
                                 YLQETGKPLDETL
                                 YLQETGKPLDETLKKA
                                 YLQETGKPLDETLKKAL
                                 YLQETGKPLDETLKKAL
                                 YLQETGKPLDETLKKAL
                                   QETGKPLDET
                                   QETGKPLDETLKKAL
                                   QETGKPLDETLKKAL
                                      GKPLDETLKKAL
                                             LKKAL
                                                AL
                                                 L
                                                 L

101      110       120       130       140       150
TGHLEEVVLALLKTPAQFDADELRAAMKGLGTDEDTLIEILASRTNKEIR
T
TG
TGH
TGHL
```

```
TGHLE
TGHLEE
TGHLEEV
T
TG
TGHLEEV
T
TGHLEEV
TGHLEEV
TGHLEEV
TGHLEEV
TGHLEEV
TGHLEEVVLA
       VLALLKTP
       VLALLKTPA
       VLALLKTPAQ
       VLALLKTPAQF
       VLALLKTPAQFD
       VLALLKTPAQFDA
       VLALLKTPAQFDAD
       VLALLKTPAQFDADE
       VLALLKTPAQFDADEL
       VLALLKTPAQFDADELR
       VLALLKTPAQFDADELRA
       VLALLKTPAQFDADELRAA
       VLALLKTPAQFDADELRAAM
       VLALLKTPAQFDADELRAAMK
       VLALLKTPAQFDADELRAAMKG
       VLALLKTPAQFDADELRAAMKGLGT
        LALLKTPAQFDADEL
        LALLKTPAQFDADELRA
         ALLKTPAQFDADEL
         ALLKTPAQFDADELRA
         ALLKTPAQFDADELRAA
          LLKTPAQFDADEL
          LLKTPAQFDADELRA
          LLKTPAQFDADELRAA
           LKTPAQFDADELRA
            KTPAQFDADEL
            KTPAQFDADELRA
            KTPAQFDADELRAA
                QFDADELRA
                QFDADELRAA
                       RAAMKGLGTDEDTLIEI
                       RAAMKGLGTDEDTLIEIL
                       RAAMKGLGTDEDTLIEILA
                       RAAMKGLGTDEDTLIEILASRT
                         AMKGLGTDEDTL
                         AMKGLGTDEDTLIE
                         AMKGLGTDEDTLIEI
                         AMKGLGTDEDTLIEIL
                         AMKGLGTDEDTLIEILA
                         AMKGLGTDEDTLIEILASRT
                                        LASRTNKEIR
                                         ASRTNKEIR
                                             NKEIR

151      160       170       180       190       200
DINRVYREELKRDLAKDITSDTSGDFRNALLSLAKGDRSEDFGVNEDLAD
DINRVYREEL
DINRVYREEL
DINRVYREEL
               KDITSDTSGDFRNALL
```

```
                  TSDTSGDFRNALL
                  TSDTSGDFRNALLSLA
                   SDTSGDFRNALL
                   SDTSGDFRNALLSLA
                      SGDFRNALLSLA
                               SLAKGDRSEDFGV
                               SLAKGDRSEDFGVNEDLA
                               SLAKGDRSEDFGVNEDLAD
                               SLAKGDRSEDFGVNEDLAD

201      210       220       230       240       250
SDARALYEAGERRKGTDVNVFNTILTTRSYPQLRRVFQKYTKYSKHDMNK
SDARA
SDARAL
     LYEAGERRKGTDVNVFNT
     LYEAGERRKGTDVNVFNTILT
         GERRKGTDVNVFNTILT
                  NVFNTILTTRSYPQLRRV
                   VFNTILTTRSYPQLRRV
                       ILTTRSYPQLRRV
                          TRSYPQLRRV
                                    FQKYTKYSKHDMNK
                                    FQKYTKYSKHDMNK
                                    FQKYTKYSKHDMNK

251      260       270       280       290       300
VLDLELKGDIEKCLTAIVKCATSKPAFFAEKLHQAMKGVGTRHKALIRIM
V
VL
VLDL
             LTAIVKCA
                    ATSKPAFFAEKL
                    ATSKPAFFAEKLHQ
                    ATSKPAFFAEKLHQA
                    ATSKPAFFAEKLHQAM
                    ATSKPAFFAEKLHQAMKGV
                    ATSKPAFFAEKLHQAMKGVGT
                    ATSKPAFFAEKLHQAMKGVGTRHKA
                     TSKPAFFAEKL
                     TSKPAFFAEKLHQAMKGV
                      SKPAFFAEKL
                          FFAEKLHQ
                          FFAEKLHQA
                          FFAEKLHQAMKGV
                                HQAMKGVGTRHKAL
                                HQAMKGVGTRHKALI
                                HQAMKGVGTRHKALIRIM
                                HQAMKGVGTRHKALIRIM
                                  AMKGVGTRHKALIRIM
                                   MKGVGTRHKALIRIM
                                       GTRHKALIRIM
                                                 M
                                                 M

301      310       320       330       340   346
VSRSEIDMNDIKAFYQKMYGISLCQAILDETKGDYEKILVALCGGN
V
VSRSEIDMNDIKA
VSRSEIDMNDIKAFYQ
VSRSEIDMNDI
VSRSEIDMNDIKA
VSRSEIDMNDIKAFYQ
VSRSEIDMNDIKAFYQKM
```

```
VSRSEIDMNDIKAFYQKMYGI
VSRSEIDMNDIKAFYQKMYGISLC
 SRSEIDMNDIKA
 SRSEIDMNDIKAFYQ
  RSEIDMNDIKA
             FYQKMYGI
             FYQKMYGISLC
             FYQKMYGISLCQA
                        QAILDETKGDYEKI
                        QAILDETKGDYEKILV
                        QAILDETKGDYEKILVA
                        QAILDETKGDYEKILVALC
                        QAILDETKGDYEKILVALCGGN
                          ILDETKGDYEKILV
                          ILDETKGDYEKILVA
                          ILDETKGDYEKILVAL
                          ILDETKGDYEKILVALC
                          ILDETKGDYEKILVALCGGN
                           LDETKGDYEKILV
                           LDETKGDYEKILVALCGGN
                               KGDYEKILVALC
                               KGDYEKILVALCGGN
```

**Folded ANXA1 600 sec**

```
1        10        20        30        40        50
MAMVSEFLKQAWFIENEEQEYVQTVKSSKGGPGSAVSPYPTFNPSSDVAA
 AMVSEFLKQ
 AMVSEFLKQA
 AMVSEFLKQAWFIENEEQEYVQTV
  MVSEFLKQ
VSEFLKQAWFIENEEQEYVQTV
 SEFLKQAWFIENEEQEYVQTV
  EFLKQAWFIENEEQEYVQT
  EFLKQAWFIENEEQEYVQTV
  EFLKQAWFIENEEQEYVQTVKS
     KQAWFIENEEQEYVQTV
       AWFIENEEQ
       AWFIENEEQEYVQT
       AWFIENEEQEYVQTV
       AWFIENEEQEYVQTVKS
        WFIENEEQ
        WFIENEEQEYV
        WFIENEEQEYVQ
        WFIENEEQEYVQT
        WFIENEEQEYVQTV
        WFIENEEQEYVQTVKS
        WFIENEEQEYVQTVKSS
        WFIENEEQEYVQTVKSSKGGPGSA
           ENEEQEYVQTV
                EYVQTVKSSKGGPGSAVSPYPT
                   QTVKSSKGGPGSAVSPYPT
                    TVKSSKGGPGSAVSPYPT
                     VKSSKGGPGSAVSPYPT
                     VKSSKGGPGSAVSPYPTFNPSSD
                     VKSSKGGPGSAVSPYPTFNPSSDVA
                      KSSKGGPGSAVSPYPT
                      KSSKGGPGSAVSPYPTFNPSS
                      KSSKGGPGSAVSPYPTFNPSSD
                      KSSKGGPGSAVSPYPTFNPSSDV
                      KSSKGGPGSAVSPYPTFNPSSDVA
                      KSSKGGPGSAVSPYPTFNPSSDVAA
                       SSKGGPGSAVSPYPT
                        SKGGPGSAVSPYPT
                         KGGPGSAVSPYPT
                                    PTFNPSSDVAA
                                     TFNPSSDVAA
                                      FNPSSDVAA
                                      FNPSSDVAA
                                      FNPSSDVAA
                                      FNPSSDVAA
                                      FNPSSDVAA
                                      FNPSSDVAA
                                      FNPSSDVAA
                                      FNPSSDVAA
                                      FNPSSDVAA
                                       NPSSDVAA
                                        PSSDVAA
                                         SSDVAA
                                              A

51       60        70        80        90        100
LHKAIMVKGVDEATIIDILTKRNNAQRQQIKAAYLQETGKPLDETLKKAL
LHKA
LHKA
L
```

```
LH
LHKA
LHKAIM
LHKAIMV
LHKAIMVK
LHKAIMVKGVDEA
LHKAIMVKGVDEAT
LHKA
LHKA
LHKA
LHKAIMVKGVDEATIIDIL
    IMVKGVDEA
    IMVKGVDEATI
    IMVKGVDEATIIDI
    IMVKGVDEATIIDIL
    IMVKGVDEATIIDILT
    IMVKGVDEATIIDILTK
    IMVKGVDEATIIDILTKR
    IMVKGVDEATIIDILTKRN
    IMVKGVDEATIIDILTKRNNA
    IMVKGVDEATIIDILTKRNNAQ
      VKGVDEATIIDIL
       KGVDEATIIDIL
       KGVDEATIIDILTKRN
       KGVDEATIIDILTKRNNAQRQQIKA
                   TKRNNAQRQQIKAAYL
                   TKRNNAQRQQIKAAYLQET
                   TKRNNAQRQQIKAAYLQETGKPLDE
                      NNAQRQQIKAAYL
                      NNAQRQQIKAAYLQET
                       NAQRQQIKAAYL
                       NAQRQQIKAAYLQET
                              KAAYLQET
                                AYLQETGKP
                                AYLQETGKPL
                                AYLQETGKPLD
                                AYLQETGKPLDE
                                AYLQETGKPLDET
                                AYLQETGKPLDETL
                                AYLQETGKPLDETLK
                                AYLQETGKPLDETLKKA
                                AYLQETGKPLDETLKKAL
                                AYLQETGKPLDETLKKAL
                                AYLQETGKPLDETLKKAL
                                AYLQETGKPLDETLKKAL
                                AYLQETGKPLDETLKKAL
                                AYLQETGKPLDETLKKAL
                                AYLQETGKPLDETLKKAL
                                AYLQETGKPLDETLKKAL
                                 YLQETGKPLD
                                 YLQETGKPLDE
                                 YLQETGKPLDET
                                 YLQETGKPLDETL
                                 YLQETGKPLDETLKKA
                                 YLQETGKPLDETLKKAL
                                 YLQETGKPLDETLKKAL
                                 YLQETGKPLDETLKKAL
                                   QETGKPLDET
                                   QETGKPLDETLKKAL
                                   QETGKPLDETLKKAL
                                    ETGKPLDETLKKAL
                                      GKPLDETLKKAL
                                        PLDETLKKAL
```

```
                                             LKKAL
                                              KKAL
                                                AL
                                                 L
                                                 L

101      110       120       130       140       150
TGHLEEVVLALLKTPAQFDADELRAAMKGLGTDEDTLIEILASRTNKEIR
T
TG
TGH
TGHL
TGHLE
TGHLEE
TGHLEEV
T
TG
TGHLEEV
T
TGHLEEV
TGHLEEV
TGHLEEV
TGHLEEV
TGHLEEV
TGHLEEV
TGHLEEV
TGHLEEV
TGHLEEVVLA
       VLALLKTP
       VLALLKTPA
       VLALLKTPAQ
       VLALLKTPAQF
       VLALLKTPAQFD
       VLALLKTPAQFDA
       VLALLKTPAQFDAD
       VLALLKTPAQFDADE
       VLALLKTPAQFDADEL
       VLALLKTPAQFDADELR
       VLALLKTPAQFDADELRA
       VLALLKTPAQFDADELRAA
       VLALLKTPAQFDADELRAAM
       VLALLKTPAQFDADELRAAMK
       VLALLKTPAQFDADELRAAMKG
       VLALLKTPAQFDADELRAAMKGLGT
        LALLKTPAQFDADEL
        LALLKTPAQFDADELRA
        LALLKTPAQFDADELRAA
         ALLKTPAQFDADEL
         ALLKTPAQFDADELRA
         ALLKTPAQFDADELRAA
         ALLKTPAQFDADELRAAM
          LLKTPAQFDADEL
          LLKTPAQFDADELRA
          LLKTPAQFDADELRAA
           LKTPAQFDADELRA
            KTPAQFDADEL
            KTPAQFDADELRA
            KTPAQFDADELRAA
            KTPAQFDADELRAAM
                QFDADELRA
                QFDADELRAA
                QFDADELRAAMKGLGTDEDTLIEI
                       RAAMKGLGTDE
```

```
                       RAAMKGLGTDEDTLIEI
                       RAAMKGLGTDEDTLIEIL
                       RAAMKGLGTDEDTLIEILA
                       RAAMKGLGTDEDTLIEILASRT
                         AMKGLGTDEDTL
                         AMKGLGTDEDTLIE
                         AMKGLGTDEDTLIEI
                         AMKGLGTDEDTLIEIL
                         AMKGLGTDEDTLIEILA
                         AMKGLGTDEDTLIEILASRT
                          MKGLGTDEDTLIEI
                          MKGLGTDEDTLIEIL
                           KGLGTDEDTLIEILA
                                        LASRTNKEIR
                                        LASRTNKEIR
                                         ASRTNKEIR
                                         ASRTNKEIR
                                         ASRTNKEIR
                                             NKEIR
                                             NKEIR

151      160       170       180       190       200
DINRVYREELKRDLAKDITSDTSGDFRNALLSLAKGDRSEDFGVNEDLAD
DI
DINRVYREEL
DI
DIN
DINRVYREEL
DINRV
DINRVYREEL
          KRDLAKDITSDTSGDFRNALL
               KDITSDTSGDFRNALL
                  TSDTSGDFRNA
                  TSDTSGDFRNALL
                  TSDTSGDFRNALLSLA
                   SDTSGDFRNA
                   SDTSGDFRNALL
                   SDTSGDFRNALLSL
                   SDTSGDFRNALLSLA
                      SGDFRNALLSLA
                               SLAKGDRSEDFGV
                               SLAKGDRSEDFGVNEDLA
                               SLAKGDRSEDFGVNEDLAD
                               SLAKGDRSEDFGVNEDLAD
                                  KGDRSEDFGVNEDLAD
                                  KGDRSEDFGVNEDLAD

201      210       220       230       240       250
SDARALYEAGERRKGTDVNVFNTILTTRSYPQLRRVFQKYTKYSKHDMNK
SDARA
SDARAL
SDARA
SDARALYEA
     LYEAGERRKGTDVNVFNT
     LYEAGERRKGTDVNVFNTILT
         GERRKGTDVNVFNTILT
                  NVFNTILT
                  NVFNTILTTRSYPQLRRV
                   VFNTILTTRSYPQLRRV
                       ILTTRSYPQL
                       ILTTRSYPQLRRV
                          TRSYPQLRRV
                          TRSYPQLRRVFQ
```

```
                                    FQKYTKYSKHDM
                                    FQKYTKYSKHDMNK
                                    FQKYTKYSKHDMNK
                                    FQKYTKYSKHDMNK
                                    FQKYTKYSKHDMNK
                                         KYSKHDMNK

251      260       270       280       290       300
VLDLELKGDIEKCLTAIVKCATSKPAFFAEKLHQAMKGVGTRHKALIRIM
V
VL
VLDL
VLDLEL
VLDL
    ELKGDIEKC
                    ATSKPAFFAEKL
                    ATSKPAFFAEKLH
                    ATSKPAFFAEKLHQ
                    ATSKPAFFAEKLHQA
                    ATSKPAFFAEKLHQAM
                    ATSKPAFFAEKLHQAMKGV
                    ATSKPAFFAEKLHQAMKGVGT
                    ATSKPAFFAEKLHQAMKGVGTRHKA
                     TSKPAFFAEKL
                     TSKPAFFAEKLHQAMKGV
                          FFAEKLHQ
                          FFAEKLHQA
                          FFAEKLHQAM
                          FFAEKLHQAMKGV
                          FFAEKLHQAMKGVGTRHKALIRIM
                                HQAMKGVGTRHKAL
                                HQAMKGVGTRHKALI
                                HQAMKGVGTRHKALIRI
                                HQAMKGVGTRHKALIRIM
                                HQAMKGVGTRHKALIRIM
                                  AMKGVGTRHKALIRIM
                                   MKGVGTRHKALIRIM
                                     GVGTRHKALIRIM
                                       GTRHKALIRIM
                                              IRIM
                                                 M
                                                 M
                                                 M

301      310       320       330       340   346
VSRSEIDMNDIKAFYQKMYGISLCQAILDETKGDYEKILVALCGGN
V
VSRSEIDMNDIKA
VSRSEIDMNDIKA
VSRSEIDMNDIKAFYQ
VSRSEIDMNDIKAFYQKM
VSRSEIDMNDI
VSRSEIDMNDIKA
VSRSEIDMNDIKAFYQ
VSRSEIDMNDIKAFYQKM
VSRSEIDMNDIKAFYQKMYGI
VSRSEIDMNDIKAFYQKMYGISLC
 SRSEIDMNDIKA
 SRSEIDMNDIKAFYQ
  RSEIDMNDIKA
  RSEIDMNDIKAFYQ
    EIDMNDIKA
             FYQKMYGI
```

```
FYQKMYGISLC
FYQKMYGISLCQA
           QAILDETKGDYEKI
           QAILDETKGDYEKILV
           QAILDETKGDYEKILVA
           QAILDETKGDYEKILVALC
           QAILDETKGDYEKILVALCGGN
             ILDETKGDYEKIL
             ILDETKGDYEKILV
             ILDETKGDYEKILVA
             ILDETKGDYEKILVAL
             ILDETKGDYEKILVALC
             ILDETKGDYEKILVALCGGN
              LDETKGDYEKILV
              LDETKGDYEKILVALCGGN
               DETKGDYEKILV
                  KGDYEKILVALC
                  KGDYEKILVALCGGN
```

**Folded ANXA1 1200 sec (9)**

```
1        10        20        30        40        50
MAMVSEFLKQAWFIENEEQEYVQTVKSSKGGPGSAVSPYPTFNPSSDVAA
MAMVSEFLKQA
 AMVSEFLKQ
 AMVSEFLKQA
  MVSEFLKQ
  MVSEFLKQA
   VSEFLKQA
   VSEFLKQAWFIENEEQEYVQTV
    SEFLKQAWFIENEEQEYVQTV
     EFLKQAWFIENEEQEYVQT
     EFLKQAWFIENEEQEYVQTV
     EFLKQAWFIENEEQEYVQTVKS
        KQAWFIENEEQEYVQTV
          AWFIENEEQ
          AWFIENEEQEYVQT
          AWFIENEEQEYVQTV
          AWFIENEEQEYVQTVKS
           WFIENEEQ
           WFIENEEQEYV
           WFIENEEQEYVQ
           WFIENEEQEYVQT
           WFIENEEQEYVQTV
           WFIENEEQEYVQTVKS
           WFIENEEQEYVQTVKSS
           WFIENEEQEYVQTVKSSKGGPGSA
              ENEEQEYVQTV
                   EYVQTVKSSKGGPGSAVSPYPT
                      QTVKSSKGGPGSAVSPYPT
                       TVKSSKGGPGSAVSPYPT
                        VKSSKGGPGSAVSPYPT
                        VKSSKGGPGSAVSPYPTFNPSSD
                        VKSSKGGPGSAVSPYPTFNPSSDVA
                         KSSKGGPGSAVSPYPT
                         KSSKGGPGSAVSPYPTFNPSS
                         KSSKGGPGSAVSPYPTFNPSSD
                         KSSKGGPGSAVSPYPTFNPSSDV
                         KSSKGGPGSAVSPYPTFNPSSDVA
                         KSSKGGPGSAVSPYPTFNPSSDVAA
                          SSKGGPGSAVSPYPT
                           SKGGPGSAVSPYPT
                            KGGPGSAVSPYPT
                                       PTFNPSSDVAA
                                        TFNPSSDVAA
                                         FNPSSDVAA
                                         FNPSSDVAA
                                         FNPSSDVAA
                                         FNPSSDVAA
                                         FNPSSDVAA
                                         FNPSSDVAA
                                         FNPSSDVAA
                                         FNPSSDVAA
                                         FNPSSDVAA
                                          NPSSDVAA
                                           PSSDVAA
                                            SSDVAA
                                                 A

51       60        70        80        90        100
LHKAIMVKGVDEATIIDILTKRNNAQRQQIKAAYLQETGKPLDETLKKAL
```

```
LHKA
LHKA
L
LH
LHKA
LHKAIM
LHKAIMV
LHKAIMVK
LHKAIMVKGVDEA
LHKAIMVKGVDEAT
LHKA
LHKA
LHKA
LHKAIMVKGVDEATIIDIL
    IMVKGVDEA
    IMVKGVDEATII
    IMVKGVDEATIIDI
    IMVKGVDEATIIDIL
    IMVKGVDEATIIDILT
    IMVKGVDEATIIDILTK
    IMVKGVDEATIIDILTKR
    IMVKGVDEATIIDILTKRN
    IMVKGVDEATIIDILTKRNNA
    IMVKGVDEATIIDILTKRNNAQ
     MVKGVDEATIIDIL
      VKGVDEATIIDI
      VKGVDEATIIDIL
       KGVDEATIIDIL
       KGVDEATIIDILTKRN
       KGVDEATIIDILTKRNNAQRQQIKA
                   TKRNNAQRQQIKAAYL
                   TKRNNAQRQQIKAAYLQET
                   TKRNNAQRQQIKAAYLQETGKPLDE
                      NNAQRQQIKAAYL
                      NNAQRQQIKAAYLQET
                       NAQRQQIKAAYL
                       NAQRQQIKAAYLQET
                              KAAYLQET
                                AYLQETGKP
                                AYLQETGKPL
                                AYLQETGKPLD
                                AYLQETGKPLDE
                                AYLQETGKPLDET
                                AYLQETGKPLDETL
                                AYLQETGKPLDETLK
                                AYLQETGKPLDETLKKA
                                AYLQETGKPLDETLKKAL
                                AYLQETGKPLDETLKKAL
                                AYLQETGKPLDETLKKAL
                                AYLQETGKPLDETLKKAL
                                AYLQETGKPLDETLKKAL
                                AYLQETGKPLDETLKKAL
                                AYLQETGKPLDETLKKAL
                                AYLQETGKPLDETLKKAL
                                 YLQETGKPLD
                                 YLQETGKPLDE
                                 YLQETGKPLDET
                                 YLQETGKPLDETL
                                 YLQETGKPLDETLKKA
                                 YLQETGKPLDETLKKAL
                                 YLQETGKPLDETLKKAL
                                 YLQETGKPLDETLKKAL
                                   QETGKPLDET
```

```
                                   QETGKPLDETLKKAL
                                   QETGKPLDETLKKAL
                                    ETGKPLDETLKKAL
                                      GKPLDETLKKAL
                                        PLDETLKKAL
                                             LKKAL
                                              KKAL
                                                AL
                                                 L
                                                 L

101      110       120       130       140       150
TGHLEEVVLALLKTPAQFDADELRAAMKGLGTDEDTLIEILASRTNKEIR
T
TG
TGH
TGHL
TGHLE
TGHLEE
TGHLEEV
T
TG
TGHLEEV
T
TGHLEEV
TGHLEEV
TGHLEEV
TGHLEEV
TGHLEEV
TGHLEEV
TGHLEEV
TGHLEEV
TGHLEEVVLA
       VLALLKTP
       VLALLKTPA
       VLALLKTPAQ
       VLALLKTPAQF
       VLALLKTPAQFD
       VLALLKTPAQFDA
       VLALLKTPAQFDAD
       VLALLKTPAQFDADE
       VLALLKTPAQFDADEL
       VLALLKTPAQFDADELR
       VLALLKTPAQFDADELRA
       VLALLKTPAQFDADELRAA
       VLALLKTPAQFDADELRAAM
       VLALLKTPAQFDADELRAAMK
       VLALLKTPAQFDADELRAAMKG
       VLALLKTPAQFDADELRAAMKGLGT
        LALLKTPAQFDADEL
        LALLKTPAQFDADELRA
        LALLKTPAQFDADELRAA
         ALLKTPAQFDADEL
         ALLKTPAQFDADELRA
         ALLKTPAQFDADELRAA
         ALLKTPAQFDADELRAAM
          LLKTPAQFDADEL
          LLKTPAQFDADELRA
          LLKTPAQFDADELRAA
          LLKTPAQFDADELRAAM
           LKTPAQFDADELRA
           LKTPAQFDADELRAA
```

```
            KTPAQFDADEL
            KTPAQFDADELRA
            KTPAQFDADELRAA
            KTPAQFDADELRAAM
                QFDADELRA
                QFDADELRAA
                QFDADELRAAMKGLGTDEDTLIEI
                QFDADELRAAMKGLGTDEDTLIEIL
                       RAAMKGLGTDE
                       RAAMKGLGTDEDTLIEI
                       RAAMKGLGTDEDTLIEIL
                       RAAMKGLGTDEDTLIEILA
                       RAAMKGLGTDEDTLIEILASRT
                         AMKGLGTDEDTL
                         AMKGLGTDEDTLIE
                         AMKGLGTDEDTLIEI
                         AMKGLGTDEDTLIEIL
                         AMKGLGTDEDTLIEILA
                         AMKGLGTDEDTLIEILASRT
                          MKGLGTDEDTLIEI
                          MKGLGTDEDTLIEIL
                          MKGLGTDEDTLIEILASRT
                           KGLGTDEDTLIEILA
                                        LASRTNKEIR
                                        LASRTNKEIR
                                         ASRTNKEIR
                                         ASRTNKEIR
                                         ASRTNKEIR
                                             NKEIR
                                             NKEIR
                                             NKEIR

151      160       170       180       190       200
DINRVYREELKRDLAKDITSDTSGDFRNALLSLAKGDRSEDFGVNEDLAD
DI
DINRVYREEL
DI
DIN
DINRV
DINRVYREEL
DINRV
DINRVYREEL
DINRVYREELKRDLAKDIT
     YREELKRDLAKDITSDT
          KRDLAKDITSDT
          KRDLAKDITSDTSGDFRNALL
          KRDLAKDITSDTSGDFRNALLSLA
               KDITSDTSGDFRNALL
                  TSDTSGDFRNA
                  TSDTSGDFRNALL
                  TSDTSGDFRNALLSLA
                   SDTSGDFRNA
                   SDTSGDFRNALL
                   SDTSGDFRNALLSL
                   SDTSGDFRNALLSLA
                      SGDFRNALLSL
                      SGDFRNALLSLA
                               SLAKGDRSEDFGV
                               SLAKGDRSEDFGVNEDLA
                               SLAKGDRSEDFGVNEDLAD
                               SLAKGDRSEDFGVNEDLAD
                                  KGDRSEDFGVNEDLAD
                                  KGDRSEDFGVNEDLAD
```

```
                                                AD

201      210       220       230       240       250
SDARALYEAGERRKGTDVNVFNTILTTRSYPQLRRVFQKYTKYSKHDMNK
SDA
SDARA
SDARAL
SDARA
SDARALYEA
SDARALYEA
     LYEAGERRKGTDVNVFNT
     LYEAGERRKGTDVNVFNTILT
         GERRKGTDVNVFNTILT
                  NVFNTILT
                  NVFNTILTTRSYPQLRRV
                   VFNTILTTRSYPQLRRV
                       ILTTRSYPQL
                       ILTTRSYPQLRRV
                          TRSYPQLRRV
                          TRSYPQLRRVFQ
                                    FQKYTKYSKHDM
                                    FQKYTKYSKHDMNK
                                    FQKYTKYSKHDMNK
                                    FQKYTKYSKHDMNK
                                    FQKYTKYSKHDMNK
                                      KYTKYSKHDMNK
                                         KYSKHDMNK
                                         KYSKHDMNK
                                         KYSKHDMNK
                                         KYSKHDMNK
                                             KHDMNK

251      260       270       280       290       300
VLDLELKGDIEKCLTAIVKCATSKPAFFAEKLHQAMKGVGTRHKALIRIM
V
VL
VLDL
VLDLEL
VLDLELKGDIEKC
VL
VLDL
VLDLELKGDIEKC
VLDLELKGDIEKCLT
VLDLELKGDIEKC
    ELKGDIEKC
    ELKGDIEKCLTAIVKC
      KGDIEKCLTAIVKC
             LTAIVKCATSKPA
               AIVKCATSKPA
               AIVKCATSKPAFFAEKL
                 VKCATSKPAFFAEKL
                  KCATSKPAFFAEKL
                    ATSKPAFFAEKL
                    ATSKPAFFAEKLH
                    ATSKPAFFAEKLHQ
                    ATSKPAFFAEKLHQA
                    ATSKPAFFAEKLHQAM
                    ATSKPAFFAEKLHQAMKGV
                    ATSKPAFFAEKLHQAMKGVGT
                    ATSKPAFFAEKLHQAMKGVGTRHKA
                     TSKPAFFAEKL
```

```
                     TSKPAFFAEKLHQ
                     TSKPAFFAEKLHQAMKGV
                      SKPAFFAEKL
                          FFAEKLHQ
                          FFAEKLHQA
                          FFAEKLHQAM
                          FFAEKLHQAMKGV
                          FFAEKLHQAMKGVGTRHKALIRI
                          FFAEKLHQAMKGVGTRHKALIRIM
                          FFAEKLHQAMKGVGTRHKALIRIM
                                HQAMKGVGTRHKAL
                                HQAMKGVGTRHKALI
                                HQAMKGVGTRHKALIRI
                                HQAMKGVGTRHKALIRIM
                                HQAMKGVGTRHKALIRIM
                                HQAMKGVGTRHKALIRIM
                                  AMKGVGTRHKALIRIM
                                   MKGVGTRHKALIRIM
                                   MKGVGTRHKALIRIM
                                     GVGTRHKALIRIM
                                       GTRHKALIRIM
                                              IRIM
                                               RIM
                                                 M
                                                 M
                                                 M
                                                 M
                                                 M

301      310       320       330       340   346
VSRSEIDMNDIKAFYQKMYGISLCQAILDETKGDYEKILVALCGGN
V
V
VS
VS
VSRSEIDMNDIKA
VSRSEIDMNDIKA
VSRSEIDMNDI
VSRSEIDMNDIKA
VSRSEIDMNDIKAFYQ
VSRSEIDMNDIKAFYQKM
VSRSEIDMNDIKAFYQKMYGISLC
VSRSEIDMNDI
VSRSEIDMNDIKA
VSRSEIDMNDIKAFY
VSRSEIDMNDIKAFYQ
VSRSEIDMNDIKAFYQKM
VSRSEIDMNDIKAFYQKMYGI
VSRSEIDMNDIKAFYQKMYGISLC
 SRSEIDMNDIKA
 SRSEIDMNDIKAFYQ
 SRSEIDMNDIKAFYQKM
 SRSEIDMNDIKAFYQKMYGISLC
  RSEIDMNDIKA
  RSEIDMNDIKAFYQ
    EIDMNDIKA
             FYQKMYGI
             FYQKMYGISLC
             FYQKMYGISLCQA
                KMYGISLCQA
                        QAILDETKGDYEKI
                        QAILDETKGDYEKIL
```

QAILDETKGDYEKILV
QAILDETKGDYEKILVA
QAILDETKGDYEKILVALC
QAILDETKGDYEKILVALCGGN
ILDETKGDYEKIL
ILDETKGDYEKILV
ILDETKGDYEKILVA
ILDETKGDYEKILVAL
ILDETKGDYEKILVALC
ILDETKGDYEKILVALCGGN
LDETKGDYEKILV
LDETKGDYEKILVALC
LDETKGDYEKILVALCGGN
DETKGDYEKILV
KGDYEKILVALC
KGDYEKILVALCGGN

**Folded ANXA1 2400 sec**

```
1        10        20        30        40        50
MAMVSEFLKQAWFIENEEQEYVQTVKSSKGGPGSAVSPYPTFNPSSDVAA
 AMVSEFLKQ
 AMVSEFLKQA
  MVSEFLKQ
  MVSEFLKQA
   VSEFLKQA
   VSEFLKQAWFIENEEQEYVQTV
    SEFLKQAWFIENEEQEYVQTV
     EFLKQAWFIENEEQEYVQT
     EFLKQAWFIENEEQEYVQTV
     EFLKQAWFIENEEQEYVQTVKS
        KQAWFIENEEQEYVQTV
          AWFIENEEQ
          AWFIENEEQEYVQT
          AWFIENEEQEYVQTV
          AWFIENEEQEYVQTVKS
           WFIENEEQ
           WFIENEEQEYV
           WFIENEEQEYVQ
           WFIENEEQEYVQT
           WFIENEEQEYVQTV
           WFIENEEQEYVQTVKS
           WFIENEEQEYVQTVKSS
           WFIENEEQEYVQTVKSSKGGPGSA
              ENEEQEYVQTV
                   EYVQTVKSSKGGPGSAVSPYPT
                      QTVKSSKGGPGSAVSPYPT
                       TVKSSKGGPGSAVSPYPT
                        VKSSKGGPGSAVSPYPT
                        VKSSKGGPGSAVSPYPTFNPSSD
                        VKSSKGGPGSAVSPYPTFNPSSDVA
                         KSSKGGPGSAVSPYPT
                         KSSKGGPGSAVSPYPTFNPSS
                         KSSKGGPGSAVSPYPTFNPSSD
                         KSSKGGPGSAVSPYPTFNPSSDV
                         KSSKGGPGSAVSPYPTFNPSSDVA
                         KSSKGGPGSAVSPYPTFNPSSDVAA
                          SSKGGPGSAVSPYPT
                           SKGGPGSAVSPYPT
                            KGGPGSAVSPYPT
                                       PTFNPSSDVAA
                                        TFNPSSDVAA
                                         FNPSSDVAA
                                         FNPSSDVAA
                                         FNPSSDVAA
                                         FNPSSDVAA
                                         FNPSSDVAA
                                         FNPSSDVAA
                                         FNPSSDVAA
                                         FNPSSDVAA
                                         FNPSSDVAA
                                          NPSSDVAA
                                           PSSDVAA
                                            SSDVAA
                                                 A

51       60        70        80        90        100
LHKAIMVKGVDEATIIDILTKRNNAQRQQIKAAYLQETGKPLDETLKKAL
LHKA
```

```
LHKA
L
LH
LHKA
LHKAIM
LHKAIMV
LHKAIMVK
LHKAIMVKGVDEA
LHKAIMVKGVDEAT
LHKA
LHKA
LHKA
LHKAIMVKGVDEATIIDIL
    IMVKGVDEA
    IMVKGVDEATI
    IMVKGVDEATII
    IMVKGVDEATIIDI
    IMVKGVDEATIIDIL
    IMVKGVDEATIIDILT
    IMVKGVDEATIIDILTK
    IMVKGVDEATIIDILTKR
    IMVKGVDEATIIDILTKRN
    IMVKGVDEATIIDILTKRNNA
    IMVKGVDEATIIDILTKRNNAQ
     MVKGVDEATIIDIL
      VKGVDEATIIDI
      VKGVDEATIIDIL
      VKGVDEATIIDILT
       KGVDEATIIDIL
       KGVDEATIIDILTKRN
       KGVDEATIIDILTKRNNAQRQQIKA
                   TKRNNAQRQQIKAAYL
                   TKRNNAQRQQIKAAYLQET
                   TKRNNAQRQQIKAAYLQETGKPLDE
                    KRNNAQRQQIKAAYL
                      NNAQRQQIKAAYL
                      NNAQRQQIKAAYLQET
                       NAQRQQIKAAYL
                       NAQRQQIKAAYLQET
                              KAAYLQET
                                AYLQETGKP
                                AYLQETGKPL
                                AYLQETGKPLD
                                AYLQETGKPLDE
                                AYLQETGKPLDET
                                AYLQETGKPLDETL
                                AYLQETGKPLDETLK
                                AYLQETGKPLDETLKKA
                                AYLQETGKPLDETLKKAL
                                AYLQETGKPLDETLKKAL
                                AYLQETGKPLDETLKKAL
                                AYLQETGKPLDETLKKAL
                                AYLQETGKPLDETLKKAL
                                AYLQETGKPLDETLKKAL
                                AYLQETGKPLDETLKKAL
                                AYLQETGKPLDETLKKAL
                                 YLQETGKPLD
                                 YLQETGKPLDE
                                 YLQETGKPLDET
                                 YLQETGKPLDETL
                                 YLQETGKPLDETLKKA
                                 YLQETGKPLDETLKKAL
                                 YLQETGKPLDETLKKAL
```

```
                                   QETGKPLDET
                                   QETGKPLDETLKKAL
                                   QETGKPLDETLKKAL
                                    ETGKPLDETLKKAL
                                      GKPLDETLKKA
                                      GKPLDETLKKAL
                                        PLDETLKKAL
                                             LKKAL
                                              KKAL
                                                AL
                                                 L
                                                 L

101      110       120       130       140       150
TGHLEEVVLALLKTPAQFDADELRAAMKGLGTDEDTLIEILASRTNKEIR
T
TG
TGH
TGHL
TGHLE
TGHLEE
TGHLEEV
T
TGHLEEV
T
TGHLEEV
TGHLEEV
TGHLEEV
TGHLEEV
TGHLEEV
TGHLEEV
TGHLEEV
TGHLEEV
TGHLEEVVLA
       VLALLKTP
       VLALLKTPA
       VLALLKTPAQ
       VLALLKTPAQF
       VLALLKTPAQFD
       VLALLKTPAQFDA
       VLALLKTPAQFDAD
       VLALLKTPAQFDADE
       VLALLKTPAQFDADEL
       VLALLKTPAQFDADELR
       VLALLKTPAQFDADELRA
       VLALLKTPAQFDADELRAA
       VLALLKTPAQFDADELRAAM
       VLALLKTPAQFDADELRAAMK
       VLALLKTPAQFDADELRAAMKG
       VLALLKTPAQFDADELRAAMKGLGT
        LALLKTPAQFDADEL
        LALLKTPAQFDADELRA
        LALLKTPAQFDADELRAA
         ALLKTPAQFDADEL
         ALLKTPAQFDADELRA
         ALLKTPAQFDADELRAA
         ALLKTPAQFDADELRAAM
          LLKTPAQFDADEL
          LLKTPAQFDADELRA
          LLKTPAQFDADELRAA
          LLKTPAQFDADELRAAM
           LKTPAQFDADELRA
```

```
           LKTPAQFDADELRAA
            KTPAQFDADEL
            KTPAQFDADELRA
            KTPAQFDADELRAA
            KTPAQFDADELRAAM
                QFDADELRA
                QFDADELRAA
                QFDADELRAAMKGLGTDEDTLIEI
                QFDADELRAAMKGLGTDEDTLIEIL
                       RAAMKGLGTDE
                       RAAMKGLGTDEDTLIEI
                       RAAMKGLGTDEDTLIEIL
                       RAAMKGLGTDEDTLIEILA
                       RAAMKGLGTDEDTLIEILASRT
                         AMKGLGTDED
                         AMKGLGTDEDTL
                         AMKGLGTDEDTLIE
                         AMKGLGTDEDTLIEI
                         AMKGLGTDEDTLIEIL
                         AMKGLGTDEDTLIEILA
                         AMKGLGTDEDTLIEILASRT
                          MKGLGTDEDTLIEI
                          MKGLGTDEDTLIEIL
                          MKGLGTDEDTLIEILASRT
                           KGLGTDEDTLIEI
                           KGLGTDEDTLIEIL
                           KGLGTDEDTLIEILA
                                        LASRTNKEIR
                                        LASRTNKEIR
                                         ASRTNKEIR
                                         ASRTNKEIR
                                         ASRTNKEIR
                                         ASRTNKEIR
                                             NKEIR
                                             NKEIR
                                             NKEIR

151      160       170       180       190       200
DINRVYREELKRDLAKDITSDTSGDFRNALLSLAKGDRSEDFGVNEDLAD
DI
DINRVYREEL
DI
DIN
DINRV
DINRVYREEL
DINRV
DINRVYREEL
DINRVYREELKRDLAKDIT
     YREELKRDLAKDITSDT
          KRDLAKDITSDT
          KRDLAKDITSDTSGDFRNALL
          KRDLAKDITSDTSGDFRNALLSLA
               KDITSDTSGDFRNALL
                  TSDTSGDFRNA
                  TSDTSGDFRNALL
                  TSDTSGDFRNALLSLA
                   SDTSGDFRNA
                   SDTSGDFRNALL
                   SDTSGDFRNALLSL
                   SDTSGDFRNALLSLA
                      SGDFRNALLSL
                      SGDFRNALLSLA
```

```
                               SLAKGDRSEDFGV
                               SLAKGDRSEDFGVNEDLA
                               SLAKGDRSEDFGVNEDLAD
                               SLAKGDRSEDFGVNEDLAD
                               SLAKGDRSEDFGVNEDLAD
                                LAKGDRSEDFGVNEDLAD
                                  KGDRSEDFGVNEDLAD
                                  KGDRSEDFGVNEDLAD
                                                AD

201      210       220       230       240       250
SDARALYEAGERRKGTDVNVFNTILTTRSYPQLRRVFQKYTKYSKHDMNK
SDA
SDARA
SDARAL
SDARA
SDARA
SDARALYEA
SDARALYEA
     LYEAGERRKGTDVNVFNT
     LYEAGERRKGTDVNVFNTILT
         GERRKGTDVNVFNTILT
                  NVFNTILT
                  NVFNTILTTRSYPQLRRV
                   VFNTILTTRSYPQLRRV
                       ILTTRSYPQL
                       ILTTRSYPQLRRV
                          TRSYPQLRRV
                          TRSYPQLRRVFQ
                                    FQKYTKYSKHDM
                                    FQKYTKYSKHDMNK
                                    FQKYTKYSKHDMNK
                                    FQKYTKYSKHDMNK
                                    FQKYTKYSKHDMNK
                                    FQKYTKYSKHDMNK
                                      KYTKYSKHDMNK
                                         KYSKHDMNK
                                         KYSKHDMNK
                                         KYSKHDMNK
                                         KYSKHDMNK
                                             KHDMNK

251      260       270       280       290       300
VLDLELKGDIEKCLTAIVKCATSKPAFFAEKLHQAMKGVGTRHKALIRIM
V
VL
VLDL
VLDLEL
VL
VLDL
VLDLELKGDIEKC
VLDLELKGDIEKCLT
VLDLELKGDIEKC
    ELKGDIEKC
    ELKGDIEKCLTAIVKC
             LTAIVKCA
             LTAIVKCATSKPA
             LTAIVKCATSKPAFFAEKL
               AIVKCATSKPA
               AIVKCATSKPAFFAEKL
                 VKCATSKPAFFAEKL
```

```
                    ATSKPAFFAEKL
                    ATSKPAFFAEKLH
                    ATSKPAFFAEKLHQ
                    ATSKPAFFAEKLHQA
                    ATSKPAFFAEKLHQAM
                    ATSKPAFFAEKLHQAMKGV
                    ATSKPAFFAEKLHQAMKGVGT
                    ATSKPAFFAEKLHQAMKGVGTRHKA
                     TSKPAFFAEKL
                     TSKPAFFAEKLHQ
                     TSKPAFFAEKLHQAMKGV
                      SKPAFFAEKL
                          FFAEKLHQ
                          FFAEKLHQA
                          FFAEKLHQAM
                          FFAEKLHQAMKGV
                          FFAEKLHQAMKGVGTRHKALIRI
                          FFAEKLHQAMKGVGTRHKALIRIM
                          FFAEKLHQAMKGVGTRHKALIRIM
                                HQAMKGVGTRHKAL
                                HQAMKGVGTRHKALI
                                HQAMKGVGTRHKALIRI
                                HQAMKGVGTRHKALIRIM
                                HQAMKGVGTRHKALIRIM
                                HQAMKGVGTRHKALIRIM
                                  AMKGVGTRHKALIRIM
                                   MKGVGTRHKALIRIM
                                     GVGTRHKALIRIM
                                       GTRHKALIRIM
                                              IRIM
                                               RIM
                                                 M
                                                 M
                                                 M
                                                 M
                                                 M

301      310       320       330       340   346
VSRSEIDMNDIKAFYQKMYGISLCQAILDETKGDYEKILVALCGGN
V
V
VS
V
VS
VSRSEIDMNDIKA
VSRSEIDMNDIKA
VSRSEIDMNDI
VSRSEIDMNDIKA
VSRSEIDMNDIKAFYQ
VSRSEIDMNDIKAFYQKM
VSRSEIDMNDIKAFYQKMYGISLC
VSRSEIDMNDI
VSRSEIDMNDIKA
VSRSEIDMNDIKAFY
VSRSEIDMNDIKAFYQ
VSRSEIDMNDIKAFYQKM
VSRSEIDMNDIKAFYQKMYGI
VSRSEIDMNDIKAFYQKMYGISLC
 SRSEIDMNDIKA
 SRSEIDMNDIKAFYQ
 SRSEIDMNDIKAFYQKM
 SRSEIDMNDIKAFYQKMYGISLC
  RSEIDMNDIKA
```

```
RSEIDMNDIKAFYQ
  EIDMNDIKA
           FYQKMYGI
           FYQKMYGISLC
           FYQKMYGISLCQA
              KMYGISLCQA
                      QAILDETKGDYEKI
                      QAILDETKGDYEKIL
                      QAILDETKGDYEKILV
                      QAILDETKGDYEKILVA
                      QAILDETKGDYEKILVALC
                      QAILDETKGDYEKILVALCGGN
                        ILDETKGDYEKIL
                        ILDETKGDYEKILV
                        ILDETKGDYEKILVA
                        ILDETKGDYEKILVAL
                        ILDETKGDYEKILVALC
                        ILDETKGDYEKILVALCGGN
                         LDETKGDYEKILV
                         LDETKGDYEKILVALC
                         LDETKGDYEKILVALCGGN
                          DETKGDYEKILV
                             KGDYEKILVALC
                             KGDYEKILVALCGGN
```

**Folded ANXA1 3600 sec**

```
1        10        20        30        40        50
MAMVSEFLKQAWFIENEEQEYVQTVKSSKGGPGSAVSPYPTFNPSSDVAA
MAMVSEFLKQA
 AMVSEFLKQ
 AMVSEFLKQA
  MVSEFLKQ
  MVSEFLKQA
   VSEFLKQA
    SEFLKQAWFIENEEQEYVQTV
     EFLKQAWFIENEEQEYVQT
     EFLKQAWFIENEEQEYVQTV
     EFLKQAWFIENEEQEYVQTVKS
        KQAWFIENEEQEYVQTV
          AWFIENEEQ
          AWFIENEEQEYVQT
          AWFIENEEQEYVQTV
          AWFIENEEQEYVQTVKS
           WFIENEEQ
           WFIENEEQEYV
           WFIENEEQEYVQ
           WFIENEEQEYVQT
           WFIENEEQEYVQTV
           WFIENEEQEYVQTVKS
           WFIENEEQEYVQTVKSS
           WFIENEEQEYVQTVKSSKGGPGSA
              ENEEQEYVQTV
                   EYVQTVKSSKGGPGSAVSPYPT
                      QTVKSSKGGPGSAVSPYPT
                       TVKSSKGGPGSAVSPYPT
                        VKSSKGGPGSAVSPYPT
                        VKSSKGGPGSAVSPYPTFNPSSD
                        VKSSKGGPGSAVSPYPTFNPSSDVA
                         KSSKGGPGSAVSPYPT
                         KSSKGGPGSAVSPYPTFNPSS
                         KSSKGGPGSAVSPYPTFNPSSD
                         KSSKGGPGSAVSPYPTFNPSSDV
                         KSSKGGPGSAVSPYPTFNPSSDVA
                         KSSKGGPGSAVSPYPTFNPSSDVAA
                          SSKGGPGSAVSPYPT
                           SKGGPGSAVSPYPT
                            KGGPGSAVSPYPT
                                       PTFNPSSDVAA
                                        TFNPSSDVAA
                                         FNPSSDVAA
                                         FNPSSDVAA
                                         FNPSSDVAA
                                         FNPSSDVAA
                                         FNPSSDVAA
                                         FNPSSDVAA
                                         FNPSSDVAA
                                         FNPSSDVAA
                                          NPSSDVAA
                                           PSSDVAA
                                            SSDVAA
                                                 A

51       60        70        80        90        100
LHKAIMVKGVDEATIIDILTKRNNAQRQQIKAAYLQETGKPLDETLKKAL
LHKA
```

```
LHKA
L
LHKA
LHKAIM
LHKAIMV
LHKAIMVK
LHKAIMVKGVDEA
LHKA
LHKA
LHKA
LHKAIMVKGVDEATIIDIL
    IMVKGVDEA
    IMVKGVDEATI
    IMVKGVDEATII
    IMVKGVDEATIIDI
    IMVKGVDEATIIDIL
    IMVKGVDEATIIDILT
    IMVKGVDEATIIDILTK
    IMVKGVDEATIIDILTKR
    IMVKGVDEATIIDILTKRN
    IMVKGVDEATIIDILTKRNNA
    IMVKGVDEATIIDILTKRNNAQ
     MVKGVDEATIIDIL
      VKGVDEATIIDI
      VKGVDEATIIDIL
      VKGVDEATIIDILT
      VKGVDEATIIDILTKRN
       KGVDEATIIDIL
       KGVDEATIIDILT
       KGVDEATIIDILTKRN
       KGVDEATIIDILTKRNNAQRQQIKA
                   TKRNNAQRQQIKAAYL
                   TKRNNAQRQQIKAAYLQET
                   TKRNNAQRQQIKAAYLQETGKPLDE
                    KRNNAQRQQIKAAYL
                      NNAQRQQIKAAYL
                      NNAQRQQIKAAYLQET
                       NAQRQQIKAAYL
                       NAQRQQIKAAYLQET
                              KAAYLQET
                                AYLQETGKP
                                AYLQETGKPL
                                AYLQETGKPLD
                                AYLQETGKPLDE
                                AYLQETGKPLDET
                                AYLQETGKPLDETL
                                AYLQETGKPLDETLK
                                AYLQETGKPLDETLKKA
                                AYLQETGKPLDETLKKAL
                                AYLQETGKPLDETLKKAL
                                AYLQETGKPLDETLKKAL
                                AYLQETGKPLDETLKKAL
                                AYLQETGKPLDETLKKAL
                                AYLQETGKPLDETLKKAL
                                AYLQETGKPLDETLKKAL
                                AYLQETGKPLDETLKKAL
                                 YLQETGKPLDE
                                 YLQETGKPLDET
                                 YLQETGKPLDETLKKA
                                 YLQETGKPLDETLKKAL
                                 YLQETGKPLDETLKKAL
                                   QETGKPLDET
                                   QETGKPLDETLKKAL
```

```
                                   QETGKPLDETLKKAL
                                    ETGKPLDETLKKAL
                                      GKPLDETLKKA
                                      GKPLDETLKKAL
                                        PLDETLKKAL
                                             LKKAL
                                              KKAL
                                                AL
                                                 L
                                                 L

101      110       120       130       140       150
TGHLEEVVLALLKTPAQFDADELRAAMKGLGTDEDTLIEILASRTNKEIR
T
TG
TGH
TGHL
TGHLE
TGHLEE
TGHLEEV
T
TGHLEEV
T
TGHLEEV
TGHLEEV
TGHLEEV
TGHLEEV
TGHLEEV
TGHLEEV
TGHLEEV
TGHLEEV
TGHLEEVVLA
       VLALLKTP
       VLALLKTPA
       VLALLKTPAQ
       VLALLKTPAQF
       VLALLKTPAQFD
       VLALLKTPAQFDA
       VLALLKTPAQFDAD
       VLALLKTPAQFDADE
       VLALLKTPAQFDADEL
       VLALLKTPAQFDADELR
       VLALLKTPAQFDADELRA
       VLALLKTPAQFDADELRAA
       VLALLKTPAQFDADELRAAM
       VLALLKTPAQFDADELRAAMK
       VLALLKTPAQFDADELRAAMKG
       VLALLKTPAQFDADELRAAMKGLGT
        LALLKTPAQFDADEL
        LALLKTPAQFDADELRA
        LALLKTPAQFDADELRAA
         ALLKTPAQFDADEL
         ALLKTPAQFDADELRA
         ALLKTPAQFDADELRAA
         ALLKTPAQFDADELRAAM
          LLKTPAQFDADEL
          LLKTPAQFDADELRA
          LLKTPAQFDADELRAA
          LLKTPAQFDADELRAAM
           LKTPAQFDADEL
           LKTPAQFDADELRA
           LKTPAQFDADELRAA
```

```
            KTPAQFDADEL
            KTPAQFDADELRA
            KTPAQFDADELRAA
                QFDADELRA
                QFDADELRAA
                QFDADELRAAMKGLGTDEDTLIEIL
                       RAAMKGLGTDE
                       RAAMKGLGTDEDTLIEI
                       RAAMKGLGTDEDTLIEIL
                       RAAMKGLGTDEDTLIEILA
                       RAAMKGLGTDEDTLIEILASRT
                         AMKGLGTDED
                         AMKGLGTDEDTL
                         AMKGLGTDEDTLIE
                         AMKGLGTDEDTLIEI
                         AMKGLGTDEDTLIEIL
                         AMKGLGTDEDTLIEILA
                         AMKGLGTDEDTLIEILASRT
                          MKGLGTDEDTLIEI
                          MKGLGTDEDTLIEIL
                          MKGLGTDEDTLIEILASRT
                           KGLGTDEDTLIEI
                           KGLGTDEDTLIEILA
                                        LASRTNKEIR
                                        LASRTNKEIR
                                        LASRTNKEIR
                                         ASRTNKEIR
                                         ASRTNKEIR
                                         ASRTNKEIR
                                         ASRTNKEIR
                                             NKEIR
                                             NKEIR
                                             NKEIR

151      160       170       180       190       200
DINRVYREELKRDLAKDITSDTSGDFRNALLSLAKGDRSEDFGVNEDLAD
DI
DINRV
DINRVYREEL
DI
DIN
DINRV
DINRVYREEL
DINRV
DINRVYREEL
DINRVYREELKRDLAKDIT
     YREELKRDLAKDITSDT
          KRDLAKDITSDT
          KRDLAKDITSDTSGDFRNALL
          KRDLAKDITSDTSGDFRNALLSLA
              AKDITSDTSGDFRNALL
               KDITSDTSGDFRNALL
                  TSDTSGDFRNA
                  TSDTSGDFRNALL
                   SDTSGDFRNA
                   SDTSGDFRNALL
                   SDTSGDFRNALLSL
                   SDTSGDFRNALLSLA
                      SGDFRNALLSL
                      SGDFRNALLSLA
                               SLAKGDRSEDFGV
                               SLAKGDRSEDFGVNEDLA
```

```
                               SLAKGDRSEDFGVNEDLAD
                               SLAKGDRSEDFGVNEDLAD
                               SLAKGDRSEDFGVNEDLAD
                                LAKGDRSEDFGVNEDLAD
                                  KGDRSEDFGVNEDLAD
                                  KGDRSEDFGVNEDLAD
                                                AD

201      210       220       230       240       250
SDARALYEAGERRKGTDVNVFNTILTTRSYPQLRRVFQKYTKYSKHDMNK
SDA
SDARA
SDARAL
SDARA
SDARA
SDARALYEA
SDARALYEA
     LYEAGERRKGTDVNVFNT
     LYEAGERRKGTDVNVFNTILT
         GERRKGTDVNVFNT
         GERRKGTDVNVFNTILT
                  NVFNTILT
                  NVFNTILTTRSYPQLRRV
                   VFNTILTTRSYPQLRRV
                       ILTTRSYPQL
                       ILTTRSYPQLRRV
                          TRSYPQLRRV
                          TRSYPQLRRVFQ
                                    FQKYTKYSKHDM
                                    FQKYTKYSKHDMNK
                                    FQKYTKYSKHDMNK
                                    FQKYTKYSKHDMNK
                                    FQKYTKYSKHDMNK
                                    FQKYTKYSKHDMNK
                                      KYTKYSKHDMNK
                                         KYSKHDMNK
                                         KYSKHDMNK
                                         KYSKHDMNK
                                         KYSKHDMNK
                                             KHDMNK

251      260       270       280       290       300
VLDLELKGDIEKCLTAIVKCATSKPAFFAEKLHQAMKGVGTRHKALIRIM
V
VL
VLDL
VLDLEL
VLDLELKGDIEKC
VL
VLDL
VLDLELKGDIEKC
VLDLELKGDIEKCLT
VLDLELKGDIEKC
    ELKGDIEKC
    ELKGDIEKCLTAIVKC
      KGDIEKCLTAIVKC
             LTAIVKCATSKPA
               AIVKCATSKPA
               AIVKCATSKPAFFAEKL
                 VKCATSKPAFFAEKL
                    ATSKPAFFAEKL
```

```
                    ATSKPAFFAEKLH
                    ATSKPAFFAEKLHQ
                    ATSKPAFFAEKLHQA
                    ATSKPAFFAEKLHQAM
                    ATSKPAFFAEKLHQAMKGV
                    ATSKPAFFAEKLHQAMKGVGT
                    ATSKPAFFAEKLHQAMKGVGTRHKA
                     TSKPAFFAEKL
                     TSKPAFFAEKLHQ
                     TSKPAFFAEKLHQAMKGV
                      SKPAFFAEKL
                          FFAEKLHQ
                          FFAEKLHQA
                          FFAEKLHQAM
                          FFAEKLHQAMKGV
                          FFAEKLHQAMKGVGTRHKALIRI
                          FFAEKLHQAMKGVGTRHKALIRIM
                          FFAEKLHQAMKGVGTRHKALIRIM
                                HQAMKGVGTRHKAL
                                HQAMKGVGTRHKALI
                                HQAMKGVGTRHKALIRI
                                HQAMKGVGTRHKALIRIM
                                HQAMKGVGTRHKALIRIM
                                HQAMKGVGTRHKALIRIM
                                  AMKGVGTRHKALIRIM
                                   MKGVGTRHKALIRIM
                                   MKGVGTRHKALIRIM
                                   MKGVGTRHKALIRIM
                                   MKGVGTRHKALIRIM
                                     GVGTRHKALIRIM
                                       GTRHKALIRIM
                                              IRIM
                                               RIM
                                                 M
                                                 M
                                                 M
                                                 M
                                                 M

301      310       320       330       340   346
VSRSEIDMNDIKAFYQKMYGISLCQAILDETKGDYEKILVALCGGN
V
V
VS
V
VS
VSRSEIDMNDIKA
VSRSEIDMNDIKA
VSRSEIDMNDI
VSRSEIDMNDIKA
VSRSEIDMNDIKAFYQ
VSRSEIDMNDIKAFYQKM
VSRSEIDMNDIKAFYQKMYGISLC
VSRSEIDMNDI
VSRSEIDMNDIKA
VSRSEIDMNDIKAFY
VSRSEIDMNDIKAFYQ
VSRSEIDMNDIKAFYQKM
VSRSEIDMNDIKAFYQKMYGI
VSRSEIDMNDIKAFYQKMYGISLC
 SRSEIDMNDIKA
 SRSEIDMNDIKAFYQ
```

```
SRSEIDMNDIKAFYQKM
SRSEIDMNDIKAFYQKMYGISLC
 RSEIDMNDIKA
 RSEIDMNDIKAFYQ
 RSEIDMNDIKAFYQKM
   EIDMNDIKA
            FYQKMYGI
            FYQKMYGISLC
            FYQKMYGISLCQA
               KMYGISLC
               KMYGISLCQA
                       QAILDETKGDYEKI
                       QAILDETKGDYEKIL
                       QAILDETKGDYEKILV
                       QAILDETKGDYEKILVA
                       QAILDETKGDYEKILVALC
                       QAILDETKGDYEKILVALCGGN
                         ILDETKGDYEKIL
                         ILDETKGDYEKILV
                         ILDETKGDYEKILVA
                         ILDETKGDYEKILVAL
                         ILDETKGDYEKILVALC
                         ILDETKGDYEKILVALCGGN
                          LDETKGDYEKILV
                          LDETKGDYEKILVALC
                          LDETKGDYEKILVALCGGN
                           DETKGDYEKILV
                              KGDYEKILVALC
                              KGDYEKILVALCGGN
```

### Supplementary data 4

#### Folded ANXA1 15 sec 1 $\mu$M HTRA1

```
1        10        20        30        40        50
MAMVSEFLKQAWFIENEEQEYVQTVKSSKGGPGSAVSPYPTFNPSSDVAA
                         KSSKGGPGSAVSPYPTFNPSSDVA

51       60        70        80        90        100
LHKAIMVKGVDEATIIDILTKRNNAQRQQIKAAYLQETGKPLDETLKKAL

101      110       120       130       140       150
TGHLEEVVLALLKTPAQFDADELRAAMKGLGTDEDTLIEILASRTNKEIR

151      160       170       180       190       200
DINRVYREELKRDLAKDITSDTSGDFRNALLSLAKGDRSEDFGVNEDLAD

201      210       220       230       240       250
SDARALYEAGERRKGTDVNVFNTILTTRSYPQLRRVFQKYTKYSKHDMNK

251      260       270       280       290       300
VLDLELKGDIEKCLTAIVKCATSKPAFFAEKLHQAMKGVGTRHKALIRIM

301      310       320       330       340   346
VSRSEIDMNDIKAFYQKMYGISLCQAILDETKGDYEKILVALCGGN
```

#### Folded ANXA1 30 sec 1 $\mu$M HTRA1

```
1        10        20        30        40        50
MAMVSEFLKQAWFIENEEQEYVQTVKSSKGGPGSAVSPYPTFNPSSDVAA
           WFIENEEQEYVQTV
                         KSSKGGPGSAVSPYPT

51       60        70        80        90        100
LHKAIMVKGVDEATIIDILTKRNNAQRQQIKAAYLQETGKPLDETLKKAL
                                AYLQETGKPLDETL
                                AYLQETGKPLDETLKKAL

101      110       120       130       140       150
TGHLEEVVLALLKTPAQFDADELRAAMKGLGTDEDTLIEILASRTNKEIR
TGHLEEV
       VLALLKTPAQFDADEL
       VLALLKTPAQFDADELRA

151      160       170       180       190       200
DINRVYREELKRDLAKDITSDTSGDFRNALLSLAKGDRSEDFGVNEDLAD

201      210       220       230       240       250
SDARALYEAGERRKGTDVNVFNTILTTRSYPQLRRVFQKYTKYSKHDMNK

251      260       270       280       290       300
VLDLELKGDIEKCLTAIVKCATSKPAFFAEKLHQAMKGVGTRHKALIRIM

301      310       320       330       340   346
VSRSEIDMNDIKAFYQKMYGISLCQAILDETKGDYEKILVALCGGN
```

**Folded ANXA1 15 sec 2 $\mu$M HTRA1**

```
1         10         20         30         40         50
MAMVSEFLKQAWFIENEEQEYVQTVKSSKGGPGSAVSPYPTFNPSSDVAA

51        60         70         80         90         100
LHKAIMVKGVDEATIIDILTKRNNAQRQQIKAAYLQETGKPLDETLKKAL
                                AYLQETGKPLDETL

101       110        120        130        140        150
TGHLEEVVLALLKTPAQFDADELRAAMKGLGTDEDTLIEILASRTNKEIR

151       160        170        180        190        200
DINRVYREELKRDLAKDITSDTSGDFRNALLSLAKGDRSEDFGVNEDLAD

201       210        220        230        240        250
SDARALYEAGERRKGTDVNVFNTILTTRSYPQLRRVFQKYTKYSKHDMNK

251       260        270        280        290        300
VLDLELKGDIEKCLTAIVKCATSKPAFFAEKLHQAMKGVGTRHKALIRIM

301       310        320        330        340   346
VSRSEIDMNDIKAFYQKMYGISLCQAILDETKGDYEKILVALCGGN
```

**Folded ANXA1 30 sec 2 μM HTRA1**

```
1        10        20        30        40        50
MAMVSEFLKQAWFIENEEQEYVQTVKSSKGGPGSAVSPYPTFNPSSDVAA
           WFIENEEQEYVQTV
                                         FNPSSDVAA

51       60        70        80        90        100
LHKAIMVKGVDEATIIDILTKRNNAQRQQIKAAYLQETGKPLDETLKKAL
LHKAIMVKGVDEAT
    IMVKGVDEATIIDI
                                AYLQETGKPLDE
                                AYLQETGKPLDETL
                                AYLQETGKPLDETLK
                                AYLQETGKPLDETLKKA
                                AYLQETGKPLDETLKKAL
                                AYLQETGKPLDETLKKAL
                                   QETGKPLDETLKKAL

101      110       120       130       140       150
TGHLEEVVLALLKTPAQFDADELRAAMKGLGTDEDTLIEILASRTNKEIR
TG
TGHLEEV
TGHLEEV
       VLALLKTPAQFDADEL
       VLALLKTPAQFDADELRA
       VLALLKTPAQFDADELRAA

151      160       170       180       190       200
DINRVYREELKRDLAKDITSDTSGDFRNALLSLAKGDRSEDFGVNEDLAD

201      210       220       230       240       250
SDARALYEAGERRKGTDVNVFNTILTTRSYPQLRRVFQKYTKYSKHDMNK

251      260       270       280       290       300
VLDLELKGDIEKCLTAIVKCATSKPAFFAEKLHQAMKGVGTRHKALIRIM

301      310       320       330       340   346
VSRSEIDMNDIKAFYQKMYGISLCQAILDETKGDYEKILVALCGGN
```

**Folded ANXA1 15 sec 4 $\mu$M HTRA1**

```
1        10        20        30        40        50
MAMVSEFLKQAWFIENEEQEYVQTVKSSKGGPGSAVSPYPTFNPSSDVAA
           WFIENEEQEYVQTV
                         KSSKGGPGSAVSPYPTFNPSSDVA

51       60        70        80        90        100
LHKAIMVKGVDEATIIDILTKRNNAQRQQIKAAYLQETGKPLDETLKKAL
    IMVKGVDEATIIDI
                                AYLQETGKPLDETL
                                AYLQETGKPLDETLKKAL
                                AYLQETGKPLDETLKKAL

101      110       120       130       140       150
TGHLEEVVLALLKTPAQFDADELRAAMKGLGTDEDTLIEILASRTNKEIR
TG
TGHLEEV
       VLALLKTPAQFDADELRA

151      160       170       180       190       200
DINRVYREELKRDLAKDITSDTSGDFRNALLSLAKGDRSEDFGVNEDLAD

201      210       220       230       240       250
SDARALYEAGERRKGTDVNVFNTILTTRSYPQLRRVFQKYTKYSKHDMNK

251      260       270       280       290       300
VLDLELKGDIEKCLTAIVKCATSKPAFFAEKLHQAMKGVGTRHKALIRIM

301      310       320       330       340   346
VSRSEIDMNDIKAFYQKMYGISLCQAILDETKGDYEKILVALCGGN
                            DETKGDYEKILV
```

**Folded ANXA1 30 sec 4 µM HTRA1**

```
1        10        20        30        40        50
MAMVSEFLKQAWFIENEEQEYVQTVKSSKGGPGSAVSPYPTFNPSSDVAA
          AWFIENEEQEYVQTV
           WFIENEEQEYVQTV
                        VKSSKGGPGSAVSPYPT
                         KSSKGGPGSAVSPYPT
                         KSSKGGPGSAVSPYPTFNPSSDVA
                            KGGPGSAVSPYPT
                                         FNPSSDVAA
                                         FNPSSDVAA
                                         FNPSSDVAA

51       60        70        80        90        100
LHKAIMVKGVDEATIIDILTKRNNAQRQQIKAAYLQETGKPLDETLKKAL
LHKA
LHKAIMVK
LHKAIMVKGVDEAT
    IMVKGVDEATIIDI
                                AYLQETGKPLD
                                AYLQETGKPLDE
                                AYLQETGKPLDET
                                AYLQETGKPLDETL
                                AYLQETGKPLDETLK
                                AYLQETGKPLDETLKKA
                                AYLQETGKPLDETLKKAL
                                AYLQETGKPLDETLKKAL
                                AYLQETGKPLDETLKKAL
                                   QETGKPLDETLKKAL

101      110       120       130       140       150
TGHLEEVVLALLKTPAQFDADELRAAMKGLGTDEDTLIEILASRTNKEIR
TG
TGHLEEV
TGHLEEV
       VLALLKTPAQFDADEL
       VLALLKTPAQFDADELRA
       VLALLKTPAQFDADELRAA
                         AMKGLGTDEDTLIEIL
                         AMKGLGTDEDTLIEILA

151      160       170       180       190       200
DINRVYREELKRDLAKDITSDTSGDFRNALLSLAKGDRSEDFGVNEDLAD

201      210       220       230       240       250
SDARALYEAGERRKGTDVNVFNTILTTRSYPQLRRVFQKYTKYSKHDMNK

251      260       270       280       290       300
VLDLELKGDIEKCLTAIVKCATSKPAFFAEKLHQAMKGVGTRHKALIRIM

301      310       320       330       340   346
VSRSEIDMNDIKAFYQKMYGISLCQAILDETKGDYEKILVALCGGN
```

### Supplementary data 5

#### Overview

| ACE project | Title |
|---|---|
| ACE_0653-01 | Time-resolved analysis of proteolytic products of ANXA1 (concentration constant 0 or 20 $\mu$M, time points 0, 15, 30 sec) by recombinant HTRA1 (concentrations: 0, 1, 2 or 4 $\mu$M) |
| ACE_0653-02 | Time-resolved analysis of proteolytic products of ANXA1 (concentration constant 0 or 20 $\mu$M, time points 0, 15, 30, 60, 90, 120, 300, 600, 1200, 2400 or 3600 sec) by recombinant HTRA1 (concentrations: 0 or 20 $\mu$M) |
| ACE_0653-03 | Time-resolved analysis of proteolytic products of ANXA1-denat (concentration constant 0 or 20 $\mu$M, time points 0, 15, 30, 60, 90, 120, 300 sec) by recombinant HTRA1 (concentrations: 0 or 20 $\mu$M) |

ANXA1 = folded ANXA1; ANXA1-denat = chemically denatured ANXA1

All 160 samples are part of one experiment (ACE_0653) and were generated at the same time. In order to reduce LC-MS variability the samples were thematically split in three part projects (ACE_0653-01, ACE_0653-02 and ACE_0653-03). In between the blocks, the LC and MS were calibrated. All samples were then analysed in one MaxQuant search (details *see* **Section "MaxQuant Search"**).

### File legend

| ACE ID | Organism | Organ/ cell line | Treatment/ experimental setup |
|---|---|---|---|
| ACE_0653-01_MK01 | Human (overexpressed in *E. coli*) | Recomb. protein | HTRA1 0 $\mu$M + ANXA1 20 $\mu$M, 0 sec, supernatant after acetone precipitation |
| ACE_0653-01_MK02 | Human (overexpressed in *E. coli*) | Recomb. protein | HTRA1 0 $\mu$M + ANXA1 20 $\mu$M, 0 sec, supernatant after acetone precipitation |
| ACE_0653-01_MK03 | Human (overexpressed in *E. coli*) | Recomb. protein | HTRA1 0 $\mu$M + ANXA1 20 $\mu$M, 0 sec, supernatant after acetone precipitation |
| ACE_0653-01_MK04 | Human (overexpressed in *E. coli*) | Recomb. protein | HTRA1 0 $\mu$M + ANXA1 20 $\mu$M, 0 sec, supernatant after acetone precipitation |
| ACE_0653-01_MK05 | Human (overexpressed in *E. coli*) | Recomb. protein | HTRA1 0 $\mu$M + ANXA1 20 $\mu$M, 30 sec, supernatant after acetone precipitation |
| ACE_0653-01_MK06 | Human (overexpressed in *E. coli*) | Recomb. protein | HTRA1 0 $\mu$M + ANXA1 20 $\mu$M, 30 sec, supernatant after acetone precipitation |
| ACE_0653-01_MK07 | Human (overexpressed in *E. coli*) | Recomb. protein | HTRA1 0 $\mu$M + ANXA1 20 $\mu$M, 30 sec, supernatant after acetone precipitation |
| ACE_0653-01_MK08 | Human (overexpressed in *E. coli*) | Recomb. protein | HTRA1 0 $\mu$M + ANXA1 20 $\mu$M, 30 sec, supernatant after acetone precipitation |
| ACE_0653-01_MK09 | Human (overexpressed in *E. coli*) | Recomb. protein | HTRA1 4 $\mu$M + ANXA1 0 $\mu$M, 0 sec, supernatant after acetone precipitation |
| ACE_0653-01_MK10 | Human (overexpressed in *E. coli*) | Recomb. protein | HTRA1 4 $\mu$M + ANXA1 0 $\mu$M, 0 sec, supernatant after acetone precipitation |
| ACE_0653-01_MK11 | Human (overexpressed in *E. coli*) | Recomb. protein | HTRA1 4 $\mu$M + ANXA1 0 $\mu$M, 0 sec, supernatant after acetone precipitation |
| ACE_0653-01_MK12 | Human (overexpressed in *E. coli*) | Recomb. protein | HTRA1 4 $\mu$M + ANXA1 0 $\mu$M, 0 sec, supernatant after acetone precipitation |
| ACE_0653-01_MK13 | Human (overexpressed in *E. coli*) | Recomb. protein | HTRA1 4 $\mu$M + ANXA1 0 $\mu$M, 30 sec, supernatant after acetone precipitation |
| ACE_0653-01_MK14 | Human (overexpressed in *E. coli*) | Recomb. protein | HTRA1 4 $\mu$M + ANXA1 0 $\mu$M, 30 sec, supernatant after acetone precipitation |
| ACE_0653-01_MK15 | Human (overexpressed in *E. coli*) | Recomb. protein | HTRA1 4 $\mu$M + ANXA1 0 $\mu$M, 30 sec, supernatant after acetone precipitation |
| ACE_0653-01_MK16 | Human (overexpressed in *E. coli*) | Recomb. protein | HTRA1 4 $\mu$M + ANXA1 0 $\mu$M, 30 sec, supernatant after acetone precipitation |
| ACE_0653-01_MK17 | Human (overexpressed in *E. coli*) | Recomb. protein | HTRA1 1 $\mu$M + ANXA1 20 $\mu$M, 0 sec, supernatant after acetone precipitation |
| ACE_0653-01_MK18 | Human (overexpressed in *E. coli*) | Recomb. protein | HTRA1 1 $\mu$M + ANXA1 20 $\mu$M, 0 sec, supernatant after acetone precipitation |
| ACE_0653-01_MK19 | Human (overexpressed in *E. coli*) | Recomb. protein | HTRA1 1 $\mu$M + ANXA1 20 $\mu$M, 0 sec, supernatant after acetone precipitation |
| ACE_0653-01_MK20 | Human (overexpressed in *E. coli*) | Recomb. protein | HTRA1 1 $\mu$M + ANXA1 20 $\mu$M, 0 sec, supernatant after acetone precipitation |
| ACE_0653-01_MK21 | Human (overexpressed in *E. coli*) | Recomb. protein | HTRA1 1 $\mu$M + ANXA1 20 $\mu$M, 15 sec, supernatant after acetone precipitation |
| ACE_0653-01_MK22 | Human (overexpressed in *E. coli*) | Recomb. protein | HTRA1 1 $\mu$M + ANXA1 20 $\mu$M, 15 sec, supernatant after acetone precipitation |
| ACE_0653-01_MK23 | Human (overexpressed in *E. coli*) | Recomb. protein | HTRA1 1 $\mu$M + ANXA1 20 $\mu$M, 15 sec, supernatant after acetone precipitation |
| ACE_0653-01_MK24 | Human (overexpressed in *E. coli*) | Recomb. protein | HTRA1 1 $\mu$M + ANXA1 20 $\mu$M, 15 sec, supernatant after acetone precipitation |
| ACE_0653-01_MK25 | Human (overexpressed in *E. coli*) | Recomb. protein | HTRA1 1 $\mu$M + ANXA1 20 $\mu$M, 30 sec, supernatant after acetone precipitation |
| ACE_0653-01_MK26 | Human (overexpressed in *E. coli*) | Recomb. protein | HTRA1 1 $\mu$M + ANXA1 20 $\mu$M, 30 sec, supernatant after acetone precipitation |
| ACE_0653-01_MK27 | Human (overexpressed in *E. coli*) | Recomb. protein | HTRA1 1 $\mu$M + ANXA1 20 $\mu$M, 30 sec, supernatant after acetone precipitation |
| ACE_0653-01_MK28 | Human (overexpressed in *E. coli*) | Recomb. protein | HTRA1 1 $\mu$M + ANXA1 20 $\mu$M, 30 sec, supernatant after acetone precipitation |
| ACE_0653-01_MK29 | Human (overexpressed in *E. coli*) | Recomb. protein | HTRA1 2 $\mu$M + ANXA1 20 $\mu$M, 0 sec, supernatant after acetone precipitation |
| ACE_0653-01_MK30 | Human (overexpressed in *E. coli*) | Recomb. protein | HTRA1 2 $\mu$M + ANXA1 20 $\mu$M, 0 sec, supernatant after acetone precipitation |
| ACE_0653-01_MK31 | Human (overexpressed in *E. coli*) | Recomb. protein | HTRA1 2 $\mu$M + ANXA1 20 $\mu$M, 0 sec, supernatant after acetone precipitation |
| ACE_0653-01_MK32 | Human (overexpressed in *E. coli*) | Recomb. protein | HTRA1 2 $\mu$M + ANXA1 20 $\mu$M, 0 sec, supernatant after acetone precipitation |
| ACE_0653-01_MK33 | Human (overexpressed in *E. coli*) | Recomb. protein | HTRA1 2 $\mu$M + ANXA1 20 $\mu$M, 15 sec, supernatant after acetone precipitation |
| ACE_0653-01_MK34 | Human (overexpressed in *E. coli*) | Recomb. protein | HTRA1 2 $\mu$M + ANXA1 20 $\mu$M, 15 sec, supernatant after acetone precipitation |
| ACE_0653-01_MK35 | Human (overexpressed in *E. coli*) | Recomb. protein | HTRA1 2 $\mu$M + ANXA1 20 $\mu$M, 15 sec, supernatant after acetone precipitation |
| ACE_0653-01_MK36 | Human (overexpressed in *E. coli*) | Recomb. protein | HTRA1 2 $\mu$M + ANXA1 20 $\mu$M, 15 sec, supernatant after acetone precipitation |
| ACE_0653-01_MK37 | Human (overexpressed in *E. coli*) | Recomb. protein | HTRA1 2 $\mu$M + ANXA1 20 $\mu$M, 30 sec, supernatant after acetone precipitation |
| ACE_0653-01_MK38 | Human (overexpressed in *E. coli*) | Recomb. protein | HTRA1 2 $\mu$M + ANXA1 20 $\mu$M, 30 sec, supernatant after acetone precipitation |
| ACE_0653-01_MK39 | Human (overexpressed in *E. coli*) | Recomb. protein | HTRA1 2 $\mu$M + ANXA1 20 $\mu$M, 30 sec, supernatant after acetone precipitation |
| ACE_0653-01_MK40 | Human (overexpressed in *E. coli*) | Recomb. protein | HTRA1 2 $\mu$M + ANXA1 20 $\mu$M, 30 sec, supernatant after acetone precipitation |
| ACE_0653-01_MK41 | Human (overexpressed in *E. coli*) | Recomb. protein | HTRA1 4 $\mu$M + ANXA1 20 $\mu$M, 0 sec, supernatant after acetone precipitation |

| | | | |
|---|---|---|---|
| ACE_0653-01_MK42 | Human (overexpressed in *E. coli*) | Recomb. protein | HTRA1 4 $\mu$M + ANXA1 20 $\mu$M, 0 sec, supernatant after acetone precipitation |
| ACE_0653-01_MK43 | Human (overexpressed in *E. coli*) | Recomb. protein | HTRA1 4 $\mu$M + ANXA1 20 $\mu$M, 0 sec, supernatant after acetone precipitation |
| ACE_0653-01_MK44 | Human (overexpressed in *E. coli*) | Recomb. protein | HTRA1 4 $\mu$M + ANXA1 20 $\mu$M, 0 sec, supernatant after acetone precipitation |
| ACE_0653-01_MK45 | Human (overexpressed in *E. coli*) | Recomb. protein | HTRA1 4 $\mu$M + ANXA1 20 $\mu$M, 15 sec, supernatant after acetone precipitation |
| ACE_0653-01_MK46 | Human (overexpressed in *E. coli*) | Recomb. protein | HTRA1 4 $\mu$M + ANXA1 20 $\mu$M, 15 sec, supernatant after acetone precipitation |
| ACE_0653-01_MK47 | Human (overexpressed in *E. coli*) | Recomb. protein | HTRA1 4 $\mu$M + ANXA1 20 $\mu$M, 15 sec, supernatant after acetone precipitation |
| ACE_0653-01_MK48 | Human (overexpressed in *E. coli*) | Recomb. protein | HTRA1 4 $\mu$M + ANXA1 20 $\mu$M, 15 sec, supernatant after acetone precipitation |
| ACE_0653-01_MK49 | Human (overexpressed in *E. coli*) | Recomb. protein | HTRA1 4 $\mu$M + ANXA1 20 $\mu$M, 30 sec, supernatant after acetone precipitation |
| ACE_0653-01_MK50 | Human (overexpressed in *E. coli*) | Recomb. protein | HTRA1 4 $\mu$M + ANXA1 20 $\mu$M, 30 sec, supernatant after acetone precipitation |
| ACE_0653-01_MK51 | Human (overexpressed in *E. coli*) | Recomb. protein | HTRA1 4 $\mu$M + ANXA1 20 $\mu$M, 30 sec, supernatant after acetone precipitation |
| ACE_0653-01_MK52 | Human (overexpressed in *E. coli*) | Recomb. protein | HTRA1 4 $\mu$M + ANXA1 20 $\mu$M, 30 sec, supernatant after acetone precipitation |
| ACE_0653-02_MK053 | Human (overexpressed in *E. coli*) | Recomb. protein | HTRA1 0 $\mu$M + ANXA1 20 $\mu$M, 0 sec, supernatant after acetone precipitation |
| ACE_0653-02_MK054 | Human (overexpressed in *E. coli*) | Recomb. protein | HTRA1 0 $\mu$M + ANXA1 20 $\mu$M, 0 sec, supernatant after acetone precipitation |
| ACE_0653-02_MK055 | Human (overexpressed in *E. coli*) | Recomb. protein | HTRA1 0 $\mu$M + ANXA1 20 $\mu$M, 0 sec, supernatant after acetone precipitation |
| ACE_0653-02_MK056 | Human (overexpressed in *E. coli*) | Recomb. protein | HTRA1 0 $\mu$M + ANXA1 20 $\mu$M, 0 sec, supernatant after acetone precipitation |
| ACE_0653-02_MK057 | Human (overexpressed in *E. coli*) | Recomb. protein | HTRA1 0 $\mu$M + ANXA1 20 $\mu$M, 3600 sec, supernatant after acetone precipitation |
| ACE_0653-02_MK058 | Human (overexpressed in *E. coli*) | Recomb. protein | HTRA1 0 $\mu$M + ANXA1 20 $\mu$M, 3600 sec, supernatant after acetone precipitation |
| ACE_0653-02_MK059 | Human (overexpressed in *E. coli*) | Recomb. protein | HTRA1 0 $\mu$M + ANXA1 20 $\mu$M, 3600 sec, supernatant after acetone precipitation |
| ACE_0653-02_MK060 | Human (overexpressed in *E. coli*) | Recomb. protein | HTRA1 0 $\mu$M + ANXA1 20 $\mu$M, 3600 sec, supernatant after acetone precipitation |
| ACE_0653-02_MK061 | Human (overexpressed in *E. coli*) | Recomb. protein | HTRA1 20 $\mu$M + ANXA1 0 $\mu$M, 0 sec, supernatant after acetone precipitation |
| ACE_0653-02_MK062 | Human (overexpressed in *E. coli*) | Recomb. protein | HTRA1 20 $\mu$M + ANXA1 0 $\mu$M, 0 sec, supernatant after acetone precipitation |
| ACE_0653-02_MK063 | Human (overexpressed in *E. coli*) | Recomb. protein | HTRA1 20 $\mu$M + ANXA1 0 $\mu$M, 0 sec, supernatant after acetone precipitation |
| ACE_0653-02_MK064 | Human (overexpressed in *E. coli*) | Recomb. protein | HTRA1 20 $\mu$M + ANXA1 0 $\mu$M, 0 sec, supernatant after acetone precipitation |
| ACE_0653-02_MK065 | Human (overexpressed in *E. coli*) | Recomb. protein | HTRA1 20 $\mu$M + ANXA1 0 $\mu$M, 3600 sec, supernatant after acetone precipitation |
| ACE_0653-02_MK066 | Human (overexpressed in *E. coli*) | Recomb. protein | HTRA1 20 $\mu$M + ANXA1 0 $\mu$M, 3600 sec, supernatant after acetone precipitation |
| ACE_0653-02_MK067 | Human (overexpressed in *E. coli*) | Recomb. protein | HTRA1 20 $\mu$M + ANXA1 0 $\mu$M, 3600 sec, supernatant after acetone precipitation |
| ACE_0653-02_MK068 | Human (overexpressed in *E. coli*) | Recomb. protein | HTRA1 20 $\mu$M + ANXA1 0 $\mu$M, 3600 sec, supernatant after acetone precipitation |
| ACE_0653-02_MK069 | Human (overexpressed in *E. coli*) | Recomb. protein | HTRA1 20 $\mu$M + ANXA1 20 $\mu$M, 0 sec, supernatant after acetone precipitation |
| ACE_0653-02_MK070 | Human (overexpressed in *E. coli*) | Recomb. protein | HTRA1 20 $\mu$M + ANXA1 20 $\mu$M, 0 sec, supernatant after acetone precipitation |
| ACE_0653-02_MK071 | Human (overexpressed in *E. coli*) | Recomb. protein | HTRA1 20 $\mu$M + ANXA1 20 $\mu$M, 0 sec, supernatant after acetone precipitation |
| ACE_0653-02_MK072 | Human (overexpressed in *E. coli*) | Recomb. protein | HTRA1 20 $\mu$M + ANXA1 20 $\mu$M, 0 sec, supernatant after acetone precipitation |
| ACE_0653-02_MK073 | Human (overexpressed in *E. coli*) | Recomb. protein | HTRA1 20 $\mu$M + ANXA1 20 $\mu$M, 15 sec, supernatant after acetone precipitation |
| ACE_0653-02_MK074 | Human (overexpressed in *E. coli*) | Recomb. protein | HTRA1 20 $\mu$M + ANXA1 20 $\mu$M, 15 sec, supernatant after acetone precipitation |
| ACE_0653-02_MK075 | Human (overexpressed in *E. coli*) | Recomb. protein | HTRA1 20 $\mu$M + ANXA1 20 $\mu$M, 15 sec, supernatant after acetone precipitation |
| ACE_0653-02_MK076 | Human (overexpressed in *E. coli*) | Recomb. protein | HTRA1 20 $\mu$M + ANXA1 20 $\mu$M, 15 sec, supernatant after acetone precipitation |
| ACE_0653-02_MK077 | Human (overexpressed in *E. coli*) | Recomb. protein | HTRA1 20 $\mu$M + ANXA1 20 $\mu$M, 30 sec, supernatant after acetone precipitation |
| ACE_0653-02_MK078 | Human (overexpressed in *E. coli*) | Recomb. protein | HTRA1 20 $\mu$M + ANXA1 20 $\mu$M, 30 sec, supernatant after acetone precipitation |
| ACE_0653-02_MK079 | Human (overexpressed in *E. coli*) | Recomb. protein | HTRA1 20 $\mu$M + ANXA1 20 $\mu$M, 30 sec, supernatant after acetone precipitation |
| ACE_0653-02_MK080 | Human (overexpressed in *E. coli*) | Recomb. protein | HTRA1 20 $\mu$M + ANXA1 20 $\mu$M, 30 sec, supernatant after acetone precipitation |
| ACE_0653-02_MK081 | Human (overexpressed in *E. coli*) | Recomb. protein | HTRA1 20 $\mu$M + ANXA1 20 $\mu$M, 60 sec, supernatant after acetone precipitation |
| ACE_0653-02_MK082 | Human (overexpressed in *E. coli*) | Recomb. protein | HTRA1 20 $\mu$M + ANXA1 20 $\mu$M, 60 sec, supernatant after acetone precipitation |
| ACE_0653-02_MK083 | Human (overexpressed in *E. coli*) | Recomb. protein | HTRA1 20 $\mu$M + ANXA1 20 $\mu$M, 60 sec, supernatant after acetone precipitation |
| ACE_0653-02_MK084 | Human (overexpressed in *E. coli*) | Recomb. protein | HTRA1 20 $\mu$M + ANXA1 20 $\mu$M, 60 sec, supernatant after acetone precipitation |

| | | | |
|---|---|---|---|
| ACE_0653-02_MK085 | Human (overexpressed in *E. coli*) | Recomb. protein | HTRA1 20 $\mu$M + ANXA1 20 $\mu$M, 90 sec, supernatant after acetone precipitation |
| ACE_0653-02_MK086 | Human (overexpressed in *E. coli*) | Recomb. protein | HTRA1 20 $\mu$M + ANXA1 20 $\mu$M, 90 sec, supernatant after acetone precipitation |
| ACE_0653-02_MK087 | Human (overexpressed in *E. coli*) | Recomb. protein | HTRA1 20 $\mu$M + ANXA1 20 $\mu$M, 90 sec, supernatant after acetone precipitation |
| ACE_0653-02_MK088 | Human (overexpressed in *E. coli*) | Recomb. protein | HTRA1 20 $\mu$M + ANXA1 20 $\mu$M, 90 sec, supernatant after acetone precipitation |
| ACE_0653-02_MK089 | Human (overexpressed in *E. coli*) | Recomb. protein | HTRA1 20 $\mu$M + ANXA1 20 $\mu$M, 120 sec, supernatant after acetone precipitation |
| ACE_0653-02_MK090 | Human (overexpressed in *E. coli*) | Recomb. protein | HTRA1 20 $\mu$M + ANXA1 20 $\mu$M, 120 sec, supernatant after acetone precipitation |
| ACE_0653-02_MK091 | Human (overexpressed in *E. coli*) | Recomb. protein | HTRA1 20 $\mu$M + ANXA1 20 $\mu$M, 120 sec, supernatant after acetone precipitation |
| ACE_0653-02_MK092 | Human (overexpressed in *E. coli*) | Recomb. protein | HTRA1 20 $\mu$M + ANXA1 20 $\mu$M, 120 sec, supernatant after acetone precipitation |
| ACE_0653-02_MK093 | Human (overexpressed in *E. coli*) | Recomb. protein | HTRA1 20 $\mu$M + ANXA1 20 $\mu$M, 300 sec, supernatant after acetone precipitation |
| ACE_0653-02_MK094 | Human (overexpressed in *E. coli*) | Recomb. protein | HTRA1 20 $\mu$M + ANXA1 20 $\mu$M, 300 sec, supernatant after acetone precipitation |
| ACE_0653-02_MK095 | Human (overexpressed in *E. coli*) | Recomb. protein | HTRA1 20 $\mu$M + ANXA1 20 $\mu$M, 300 sec, supernatant after acetone precipitation |
| ACE_0653-02_MK096 | Human (overexpressed in *E. coli*) | Recomb. protein | HTRA1 20 $\mu$M + ANXA1 20 $\mu$M, 300 sec, supernatant after acetone precipitation |
| ACE_0653-02_MK097 | Human (overexpressed in *E. coli*) | Recomb. protein | HTRA1 20 $\mu$M + ANXA1 20 $\mu$M, 600 sec, supernatant after acetone precipitation |
| ACE_0653-02_MK098 | Human (overexpressed in *E. coli*) | Recomb. protein | HTRA1 20 $\mu$M + ANXA1 20 $\mu$M, 600 sec, supernatant after acetone precipitation |
| ACE_0653-02_MK099 | Human (overexpressed in *E. coli*) | Recomb. protein | HTRA1 20 $\mu$M + ANXA1 20 $\mu$M, 600 sec, supernatant after acetone precipitation |
| ACE_0653-02_MK100 | Human (overexpressed in *E. coli*) | Recomb. protein | HTRA1 20 $\mu$M + ANXA1 20 $\mu$M, 600 sec, supernatant after acetone precipitation |
| ACE_0653-02_MK101 | Human (overexpressed in *E. coli*) | Recomb. protein | HTRA1 20 $\mu$M + ANXA1 20 $\mu$M, 1200 sec, supernatant after acetone precipitation |
| ACE_0653-02_MK102 | Human (overexpressed in *E. coli*) | Recomb. protein | HTRA1 20 $\mu$M + ANXA1 20 $\mu$M, 1200 sec, supernatant after acetone precipitation |
| ACE_0653-02_MK103 | Human (overexpressed in *E. coli*) | Recomb. protein | HTRA1 20 $\mu$M + ANXA1 20 $\mu$M, 1200 sec, supernatant after acetone precipitation |
| ACE_0653-02_MK104 | Human (overexpressed in *E. coli*) | Recomb. protein | HTRA1 20 $\mu$M + ANXA1 20 $\mu$M, 1200 sec, supernatant after acetone precipitation |
| ACE_0653-02_MK105 | Human (overexpressed in *E. coli*) | Recomb. protein | HTRA1 20 $\mu$M + ANXA1 20 $\mu$M, 2400 sec, supernatant after acetone precipitation |
| ACE_0653-02_MK106 | Human (overexpressed in *E. coli*) | Recomb. protein | HTRA1 20 $\mu$M + ANXA1 20 $\mu$M, 2400 sec, supernatant after acetone precipitation |
| ACE_0653-02_MK107 | Human (overexpressed in *E. coli*) | Recomb. protein | HTRA1 20 $\mu$M + ANXA1 20 $\mu$M, 2400 sec, supernatant after acetone precipitation |
| ACE_0653-02_MK108 | Human (overexpressed in *E. coli*) | Recomb. protein | HTRA1 20 $\mu$M + ANXA1 20 $\mu$M, 2400 sec, supernatant after acetone precipitation |
| ACE_0653-02_MK109 | Human (overexpressed in *E. coli*) | Recomb. protein | HTRA1 20 $\mu$M + ANXA1 20 $\mu$M, 3600 sec, supernatant after acetone precipitation |
| ACE_0653-02_MK110 | Human (overexpressed in *E. coli*) | Recomb. protein | HTRA1 20 $\mu$M + ANXA1 20 $\mu$M, 3600 sec, supernatant after acetone precipitation |
| ACE_0653-02_MK111 | Human (overexpressed in *E. coli*) | Recomb. protein | HTRA1 20 $\mu$M + ANXA1 20 $\mu$M, 3600 sec, supernatant after acetone precipitation |
| ACE_0653-02_MK112 | Human (overexpressed in *E. coli*) | Recomb. protein | HTRA1 20 $\mu$M + ANXA1 20 $\mu$M, 3600 sec, supernatant after acetone precipitation |
| ACE_0653-03_MK113 | Human (overexpressed in *E. coli*) | Recomb. protein | HTRA1 0 $\mu$M + “denat.” ANXA1 20 $\mu$M, 0 sec, supernatant after acetone precipitation |
| ACE_0653-03_MK114 | Human (overexpressed in *E. coli*) | Recomb. protein | HTRA1 0 $\mu$M + “denat.” ANXA1 20 $\mu$M, 0 sec, supernatant after acetone precipitation |
| ACE_0653-03_MK115 | Human (overexpressed in *E. coli*) | Recomb. protein | HTRA1 0 $\mu$M + “denat.” ANXA1 20 $\mu$M, 0 sec, supernatant after acetone precipitation |
| ACE_0653-03_MK116 | Human (overexpressed in *E. coli*) | Recomb. protein | HTRA1 0 $\mu$M + “denat.” ANXA1 20 $\mu$M, 0 sec, supernatant after acetone precipitation |
| ACE_0653-03_MK117 | Human (overexpressed in *E. coli*) | Recomb. protein | HTRA1 0 $\mu$M + “denat.” ANXA1 20 $\mu$M, 600 sec, supernatant after acetone precipitation |
| ACE_0653-03_MK118 | Human (overexpressed in *E. coli*) | Recomb. protein | HTRA1 0 $\mu$M + “denat.” ANXA1 20 $\mu$M, 600 sec, supernatant after acetone precipitation |
| ACE_0653-03_MK119 | Human (overexpressed in *E. coli*) | Recomb. protein | HTRA1 0 $\mu$M + “denat.” ANXA1 20 $\mu$M, 600 sec, supernatant after acetone precipitation |
| ACE_0653-03_MK120 | Human (overexpressed in *E. coli*) | Recomb. protein | HTRA1 0 $\mu$M + “denat.” ANXA1 20 $\mu$M, 600 sec, supernatant after acetone precipitation |
| ACE_0653-03_MK121 | Human (overexpressed in *E. coli*) | Recomb. protein | HTRA1 20 $\mu$M + “denat.” ANXA1 0 $\mu$M, 0 sec, supernatant after acetone precipitation |
| ACE_0653-03_MK122 | Human (overexpressed in *E. coli*) | Recomb. protein | HTRA1 20 $\mu$M + “denat.” ANXA1 0 $\mu$M, 0 sec, supernatant after acetone precipitation |
| ACE_0653-03_MK123 | Human (overexpressed in *E. coli*) | Recomb. protein | HTRA1 20 $\mu$M + “denat.” ANXA1 0 $\mu$M, 0 sec, supernatant after acetone precipitation |
| ACE_0653-03_MK124 | Human (overexpressed in *E. coli*) | Recomb. protein | HTRA1 20 $\mu$M + “denat.” ANXA1 0 $\mu$M, 0 sec, supernatant after acetone precipitation |
| ACE_0653-03_MK125 | Human (overexpressed in *E. coli*) | Recomb. protein | HTRA1 20 $\mu$M + “denat.” ANXA1 0 $\mu$M, 3600 sec, supernatant after acetone precipitation |
| ACE_0653-03_MK126 | Human (overexpressed in *E. coli*) | Recomb. protein | HTRA1 20 $\mu$M + “denat.” ANXA1 0 $\mu$M, 3600 sec, supernatant after acetone precipitation |
| ACE_0653-03_MK127 | Human (overexpressed in *E. coli*) | Recomb. protein | HTRA1 20 $\mu$M + “denat.” ANXA1 0 $\mu$M, 3600 sec, supernatant after acetone precipitation |

| | | | |
|---|---|---|---|
| ACE_0653-03_MK128 | Human (overexpressed in *E. coli*) | Recomb. protein | HTRA1 20 $\mu$M + “denat.” ANXA1 0 $\mu$M, 3600 sec, supernatant after acetone precipitation |
| ACE_0653-03_MK129 | Human (overexpressed in *E. coli*) | Recomb. protein | HTRA1 20 $\mu$M + “denat.” ANXA1 20 $\mu$M, 0 sec, supernatant after acetone precipitation |
| ACE_0653-03_MK130 | Human (overexpressed in *E. coli*) | Recomb. protein | HTRA1 20 $\mu$M + “denat.” ANXA1 20 $\mu$M, 0 sec, supernatant after acetone precipitation |
| ACE_0653-03_MK131 | Human (overexpressed in *E. coli*) | Recomb. protein | HTRA1 20 $\mu$M + “denat.” ANXA1 20 $\mu$M, 0 sec, supernatant after acetone precipitation |
| ACE_0653-03_MK132 | Human (overexpressed in *E. coli*) | Recomb. protein | HTRA1 20 $\mu$M + “denat.” ANXA1 20 $\mu$M, 0 sec, supernatant after acetone precipitation |
| ACE_0653-03_MK133 | Human (overexpressed in *E. coli*) | Recomb. protein | HTRA1 20 $\mu$M + “denat.” ANXA1 20 $\mu$M, 15 sec, supernatant after acetone precipitation |
| ACE_0653-03_MK134 | Human (overexpressed in *E. coli*) | Recomb. protein | HTRA1 20 $\mu$M + “denat.” ANXA1 20 $\mu$M, 15 sec, supernatant after acetone precipitation |
| ACE_0653-03_MK135 | Human (overexpressed in *E. coli*) | Recomb. protein | HTRA1 20 $\mu$M + “denat.” ANXA1 20 $\mu$M, 15 sec, supernatant after acetone precipitation |
| ACE_0653-03_MK136 | Human (overexpressed in *E. coli*) | Recomb. protein | HTRA1 20 $\mu$M + “denat.” ANXA1 20 $\mu$M, 15 sec, supernatant after acetone precipitation |
| ACE_0653-03_MK137 | Human (overexpressed in *E. coli*) | Recomb. protein | HTRA1 20 $\mu$M + “denat.” ANXA1 20 $\mu$M, 30 sec, supernatant after acetone precipitation |
| ACE_0653-03_MK138 | Human (overexpressed in *E. coli*) | Recomb. protein | HTRA1 20 $\mu$M + “denat.” ANXA1 20 $\mu$M, 30 sec, supernatant after acetone precipitation |
| ACE_0653-03_MK139 | Human (overexpressed in *E. coli*) | Recomb. protein | HTRA1 20 $\mu$M + “denat.” ANXA1 20 $\mu$M, 30 sec, supernatant after acetone precipitation |
| ACE_0653-03_MK140 | Human (overexpressed in *E. coli*) | Recomb. protein | HTRA1 20 $\mu$M + “denat.” ANXA1 20 $\mu$M, 30 sec, supernatant after acetone precipitation |
| ACE_0653-03_MK141 | Human (overexpressed in *E. coli*) | Recomb. protein | HTRA1 20 $\mu$M + “denat.” ANXA1 20 $\mu$M, 60 sec, supernatant after acetone precipitation |
| ACE_0653-03_MK142 | Human (overexpressed in *E. coli*) | Recomb. protein | HTRA1 20 $\mu$M + “denat.” ANXA1 20 $\mu$M, 60 sec, supernatant after acetone precipitation |
| ACE_0653-03_MK143 | Human (overexpressed in *E. coli*) | Recomb. protein | HTRA1 20 $\mu$M + “denat.” ANXA1 20 $\mu$M, 60 sec, supernatant after acetone precipitation |
| ACE_0653-03_MK144 | Human (overexpressed in *E. coli*) | Recomb. protein | HTRA1 20 $\mu$M + “denat.” ANXA1 20 $\mu$M, 60 sec, supernatant after acetone precipitation |
| ACE_0653-03_MK145 | Human (overexpressed in *E. coli*) | Recomb. protein | HTRA1 20 $\mu$M + “denat.” ANXA1 20 $\mu$M, 90 sec, supernatant after acetone precipitation |
| ACE_0653-03_MK146 | Human (overexpressed in *E. coli*) | Recomb. protein | HTRA1 20 $\mu$M + “denat.” ANXA1 20 $\mu$M, 90 sec, supernatant after acetone precipitation |
| ACE_0653-03_MK147 | Human (overexpressed in *E. coli*) | Recomb. protein | HTRA1 20 $\mu$M + “denat.” ANXA1 20 $\mu$M, 90 sec, supernatant after acetone precipitation |
| ACE_0653-03_MK148 | Human (overexpressed in *E. coli*) | Recomb. protein | HTRA1 20 $\mu$M + “denat.” ANXA1 20 $\mu$M, 90 sec, supernatant after acetone precipitation |
| ACE_0653-03_MK149 | Human (overexpressed in *E. coli*) | Recomb. protein | HTRA1 20 $\mu$M + “denat.” ANXA1 20 $\mu$M, 120 sec, supernatant after acetone precipitation |
| ACE_0653-03_MK150 | Human (overexpressed in *E. coli*) | Recomb. protein | HTRA1 20 $\mu$M + “denat.” ANXA1 20 $\mu$M, 120 sec, supernatant after acetone precipitation |
| ACE_0653-03_MK151 | Human (overexpressed in *E. coli*) | Recomb. protein | HTRA1 20 $\mu$M + “denat.” ANXA1 20 $\mu$M, 120 sec, supernatant after acetone precipitation |
| ACE_0653-03_MK152 | Human (overexpressed in *E. coli*) | Recomb. protein | HTRA1 20 $\mu$M + “denat.” ANXA1 20 $\mu$M, 120 sec, supernatant after acetone precipitation |
| ACE_0653-03_MK153 | Human (overexpressed in *E. coli*) | Recomb. protein | HTRA1 20 $\mu$M + “denat.” ANXA1 20 $\mu$M, 300 sec, supernatant after acetone precipitation |
| ACE_0653-03_MK154 | Human (overexpressed in *E. coli*) | Recomb. protein | HTRA1 20 $\mu$M + “denat.” ANXA1 20 $\mu$M, 300 sec, supernatant after acetone precipitation |
| ACE_0653-03_MK155 | Human (overexpressed in *E. coli*) | Recomb. protein | HTRA1 20 $\mu$M + “denat.” ANXA1 20 $\mu$M, 300 sec, supernatant after acetone precipitation |
| ACE_0653-03_MK156 | Human (overexpressed in *E. coli*) | Recomb. protein | HTRA1 20 $\mu$M + “denat.” ANXA1 20 $\mu$M, 300 sec, supernatant after acetone precipitation |
| ACE_0653-03_MK157 | Human (overexpressed in *E. coli*) | Recomb. protein | HTRA1 20 $\mu$M + “denat.” ANXA1 20 $\mu$M, 600 sec, supernatant after acetone precipitation |
| ACE_0653-03_MK158 | Human (overexpressed in *E. coli*) | Recomb. protein | HTRA1 20 $\mu$M + “denat.” ANXA1 20 $\mu$M, 600 sec, supernatant after acetone precipitation |
| ACE_0653-03_MK159 | Human (overexpressed in *E. coli*) | Recomb. protein | HTRA1 20 $\mu$M + “denat.” ANXA1 20 $\mu$M, 600 sec, supernatant after acetone precipitation |
| ACE_0653-03_MK160 | Human (overexpressed in *E. coli*) | Recomb. protein | HTRA1 20 $\mu$M + “denat.” ANXA1 20 $\mu$M, 600 sec, supernatant after acetone precipitation |

### LC Settings

| | |
|---|---|
| MS device | Orbitrap Elite |
| LC device | Evosep One |
| Ion source | Thermo Nanospray Flex |
| **Analytical column** | EV1064 Analytical Column – 60 & 100 samples/day |
| Column diameter | Length ($L_C$) = 8 cm; ID = 100; OD = PEEK; emitter EV-1086 Stainless steel emitter |
| Stationary phase | Dr Maisch C18 AQ, 3 $\mu$m beads |
| Particle diameter ($d_p$) | 3 $\mu$m |
| Pore size | 120 Å |
| Column ID | AC_EVO-003 |
| **Solvents** | A: 0.1% FA in UPLC water<br>B: 0.1% FA in UPLC ACN |
| Gradient | 21 min gradient |

### MS Settings

| Project | MS | general | MS1 | MS2 | MS2 | MS3 | Comments; special settings |
|---|---|---|---|---|---|---|---|
| ACE_0653 | Elite | Tune v2.7.0.1112 SP2<br>Gradient: 21 min | Analyzer: FT<br>Res.: 60000<br>SR: 300 - 1500<br>AGC: 3× $10^6$<br>AcT: 50<br>RF: 30<br>SF: --<br>DDM: NS/15 | Analyzer: IT<br>Res./ScR: -/rapid<br>SR: Auto<br>AGC: 1× $10^4$<br>AcT: 60<br>CS: +2 and higher<br>IsM: IT<br>IsW: 2.0<br>Frag.: CID<br>NCE: 35 | | | Classic orbitrap experiment: MS1 in Orbitrap at high resolution and data dependent MS2 in Iontrap at rapid scan rate. Dynamic exclusion enabled (exclude after n times=1; Exclusion duration (s)= 30; mass tolerance= ± 100 ppm), exclusion list size: 500 |

**FT**= Fourier Transform (Orbitrap); **IT**= Iontrap; **Q**= Quadrupol; **Res.**= max. Resolution at 200 m/z (Lumos) or 400 m/z (Elite) [FWHM (full width at half maximum)]; **ScR**= scan rate for measurements in the IT; **SR**= scan range [m/z]; **AGC**= automatic gain control, max number of acquired ions per measurement; **AcT**= max. Ion acquisition time [ms]; **CS**= charge states used for fragmentation; **IsM**= Isolation mode (Q or IT), MS2 isolation and further is only done in IT; **IsW**= Isolation window [m/z], value followed by scan mode the isolation is based on (MS1, MS2 …) **Frag.**= Fragmentation method; **HCD**= Higher-energy collisional dissociation; **CID**= Collision-induced dissociation; **ETD**= Electron-transfer dissociation**; EThcD=** Electron-Transfer/Higher-Energy Collision Dissociation; **sHCD**= stepped HCD**; NCE**= normalized collision energy; **cycles**: number of MSn recorded or max cycle time; **RF**= RF Lens [%]; **SF**= Source Fragmentation [V]; **DDM**: Data dependent Mode (cycle time in seconds, CT/[s] or number of scans, NS); **NS**= Number of data dependent scans

### MaxQuant Search

| | |
|---|---|
| Program & version | MaxQuant v2.0.2.0 |
| Search engine | Andromeda |
| Settings | Basically default; LFQ and MBR were turned on. Normalization was turned off |
| Static modification | none |
| Digestion mode | unspecific |
| Dynamic modification | Acetyl (N-term); Oxidation (M) |
| Modification included in quantification | Oxidation (M) |
| Databases | 1. Contaminants<br>2. ACE_0653_UP000000625_83333.fasta |

The unfiltered results of the MaxQuant Search can be found here:

| | | | |
|---|---|---|---|
| ACE_0653_all_in_modificationSpecificPep... | 1/13/2022 10:03 AM | Text Document | 6,024 KB |
| ACE_0653_all_in_parameters.txt | 1/13/2022 10:03 AM | Text Document | 4 KB |
| ACE_0653_all_in_peptides.txt | 1/13/2022 10:03 AM | Text Document | 7,183 KB |
| ACE_0653_all_in_proteinGroups.txt | 1/13/2022 10:03 AM | Text Document | 2,253 KB |
| ACE_0653_all_in_summary.txt | 1/13/2022 10:04 AM | Text Document | 42 KB |
| ACE_0653_all_in_tables.pdf | 1/13/2022 10:03 AM | Adobe Acrobat D... | 179 KB |

### Supplementary data 6

### Example for how UMSAP calculates the relative frequency of cleavages

In a MS experiment the following peptides were identified. They all share the same P1 site at V231

A - RGHYV
B - GHYV
C - DFRTGH
D - DFRTGHKL
E - DFRTGHKLM
F - DFRTGHKLMT

Average intensities. 0 values mean that the peptide was not detected in the given time point or the intensity values are not significantly different to the control experiments.

| Time Point | A | B | C | D | E | F |
|---|---|---|---|---|---|---|
| 5 min | 2 | 0 | 0 | 3 | 3 | 10 |
| 15 min | 4 | 1 | 0 | 0 | 6 | 5 |
| 30 min | 8 | 3 | 5 | 0 | 0 | 1 |

The first non-zero average intensity along the time points is taken as reference for the calculation of the relative intensity.

| | Relative Intensities | | | | | | Relative cleavage frequency |
|---|---|---|---|---|---|---|---|
| Time-point | A | B | C | D | E | F | P1 (V231) |
| 5 min | 2/2 = 1 | 0/1 = 0 | 0/5 = 0 | 3/3 = 1 | 3/3 = 1 | 10/10 = 1 | 1+0+0+1+1+1 = 4 |
| 15 min | 4/2 = 2 | 1/1 = 1 | 0/5 = 0 | 0/3 = 0 | 6/3 = 2 | 5/10 = 0.5 | 2+1+0+0+2+0.5 = 5.5 |
| 30 min | 8/2 = 4 | 3/1 = 1 | 5/5 = 1 | 0/3 = 0 | 0/3 = 0 | 1/10 = 0.1 | 4+1+1+0+0+0.1 = 6.1 |

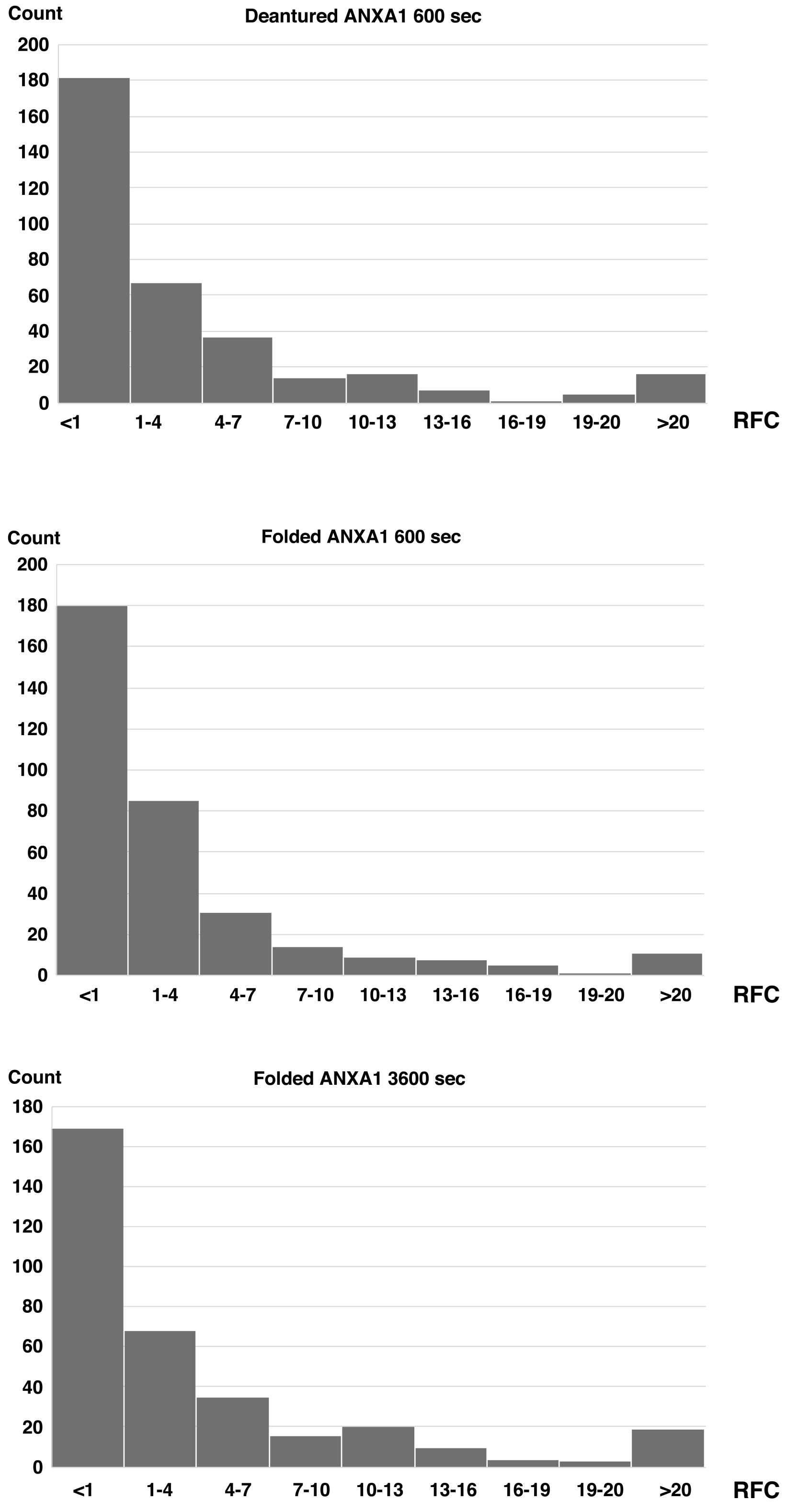

**SI Fig. 1. Histogram of relative cleavage efficiencies.** To classify cleavage sites, the relative cleavage efficiencies after each residue was calculated by UMSAP. The histogram was generated by Xcel using width of interval: 3, underflow bin: 0.99; overflow bin: 20 as parameters. RFC, relative frequency of cuts.

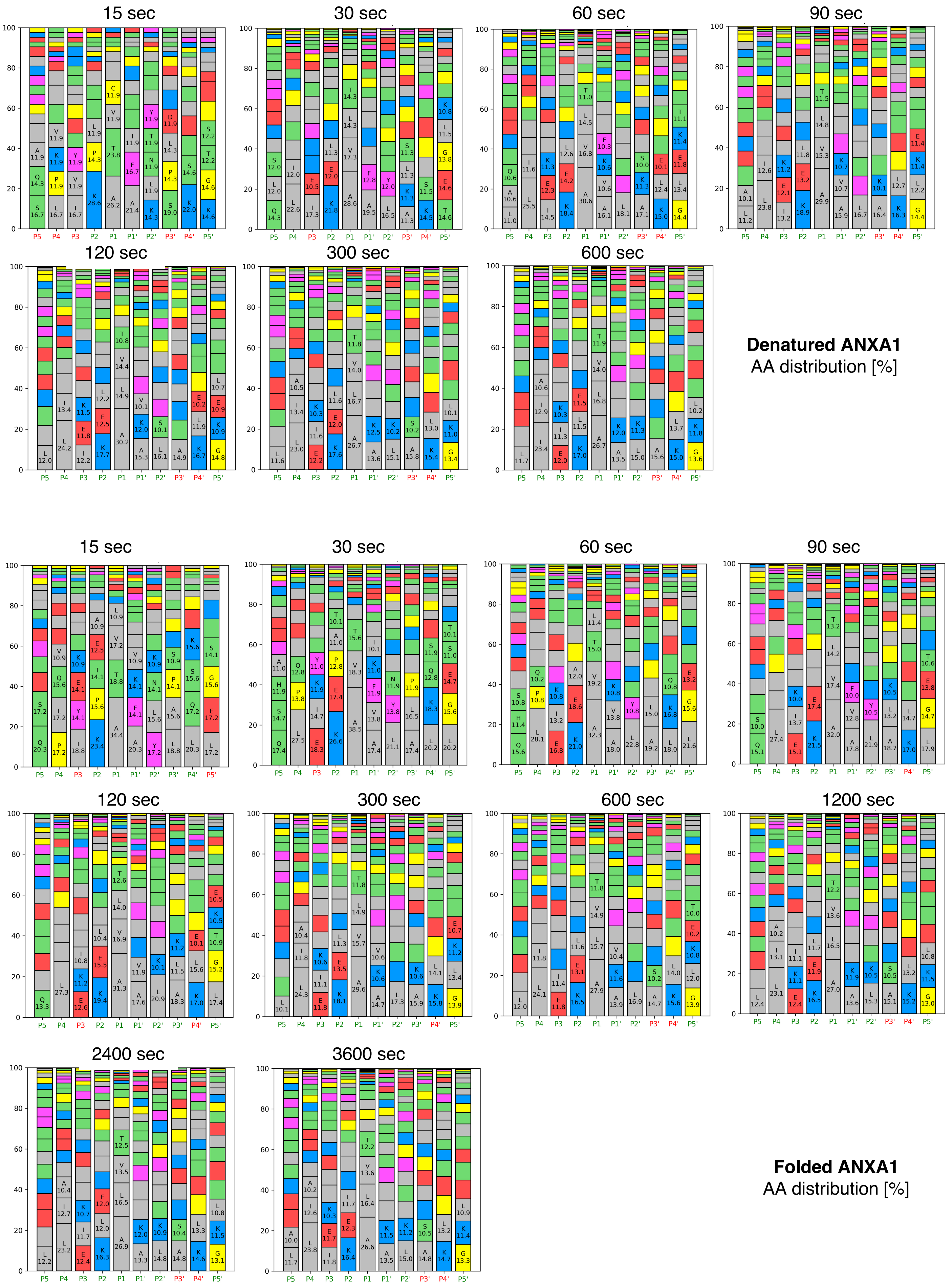

**SI Fig. 2. Relative amino acid distribution at sites P1-P5 and P1'-P5' after various times of incubation.** Red letter, no protease selectivity toward specific amino acids at this position; green letter, protease selectivity toward certain amino acids at this position.

| Denatured ANXA1 | | | | | | | | | | | Folded ANXA1 | | | | | | | | | |
|---|---|---|---|---|---|---|---|---|---|---|---|---|---|---|---|---|---|---|---|---|
| P5 | P4 | P3 | P2 | P1 | P1' | P2' | P3' | P4' | P5' | Time sec | P5 | P4 | P3 | P2 | P1 | P1' | P2' | P3' | P4' | P5' |
| | | | K 29<br>P 14<br>L 12 | A 26<br>T 24<br>V 12<br>C 12 | A 21<br>F 17<br>I 12 | K 14<br>L 12<br>N 12<br>T 12<br>Y 12 | | | K 15<br>G 15<br>T 12<br>S 12 | 15 | Q 20<br>S 17 | P 17<br>L 17<br>Q 16<br>V 11 | | K 24<br>P 16<br>T 14<br>E 13<br>A 11 | A 34<br>T 19<br>V 17<br>L 11 | A 20<br>F 14<br>K 14<br>V 11 | Y 17<br>L 16<br>N 14<br>K 11 | L 19<br>A 16<br>P 11<br>S 11 | L 20<br>Q 17<br>S 16<br>K 16 | L 17<br>E 17<br>G 16<br>S 14 |
| Q 14<br>L 12<br>S 12 | L 23<br>I 12 | | K 22<br>E 12<br>L 11 | A 29<br>V 17<br>L 14<br>T 14 | A 19<br>F 13 | L 17<br>Y 12 | | | T 15<br>E 15<br>G 14<br>L 12<br>K 11 | 30 | Q 17<br>S 15<br>H 12<br>A 11 | L 28<br>P 14<br>Q 13 | | K 27<br>E 17<br>P 13<br>A 11<br>T 10 | A 39<br>V 18<br>T 16 | A 17<br>V 14<br>F 12<br>K 11<br>I 10 | L 21<br>Y 14<br>N 12 | A 17<br>L 17<br>P 12 | L 20<br>K 18<br>Q 13<br>S 12 | L 20<br>G 16<br>E 15<br>S 11<br>T 10 |
| L 11<br>A 11<br>Q 11 | L 26<br>I 12 | I 15<br>E 12<br>K 11 | K 18<br>E 14<br>L 13 | A 31<br>V 17<br>L 15<br>T 11 | A 16<br>V 11<br>K 11<br>F 10 | L 18 | | | G 14<br>L 13<br>E 12<br>K 11<br>T 11 | 60 | Q 16<br>H 11<br>S 11 | L 28<br>P 11<br>Q 10 | E 17<br>I 13<br>K 11 | K 21<br>E 19<br>A 12 | A 32<br>V 19<br>T 15<br>L 11 | A 18<br>V 14<br>K 11 | L 23<br>Y 11 | A 19<br>L 15 | L 18<br>K 17<br>Q 11 | L 22<br>G 16<br>E 13 |
| L 11<br>A 10 | L 24<br>I 13 | I 13<br>E 12<br>K 11 | K 19<br>E 13<br>L 12 | A 30<br>L 15<br>V 15<br>T 12 | A 16<br>V 11<br>K 11 | L 17 | | | G 14<br>L 12<br>K 11<br>E 11 | 90 | Q 15<br>S 10 | L 27 | E 15<br>I 14<br>K 10 | K 22<br>E 17 | A 32<br>V 17<br>L 14<br>T 13 | A 18<br>V 13<br>F 10 | L 22<br>Y 11 | A 19<br>L 13<br>K 11 | | L 18<br>G 15<br>E 14<br>T 11 |
| L 12 | L 24<br>I 13 | I 12<br>E 12<br>K 12 | K 18<br>E 13<br>L 12 | A 30<br>L 15<br>V 15<br>T 11 | A 15<br>K 12<br>V 10 | L 16<br>S 10 | | | G 15<br>K 11<br>E 11<br>L 11 | 120 | Q 13 | L 27 | | K 19<br>E 16<br>L 10 | A 31<br>V 17<br>L 14<br>T 13 | A 18<br>V 12 | L 21<br>K 10 | A 18<br>L 12<br>K 11 | | L 17<br>G 15<br>T 11<br>K 11<br>E 11 |
| L 12 | L 23<br>I 13<br>A 11 | E 12<br>I 12<br>K 10 | K 18<br>E 12<br>L 12 | A 27<br>L 17<br>V 14<br>T 12 | A 14<br>K 13 | L 15<br>K 10 | | | G 13<br>K 11<br>L 10 | 300 | L 10 | L 24<br>I 12<br>A 10 | E 12<br>I 11<br>K 11 | K 18<br>E 14<br>L 11 | A 30<br>V 16<br>L 15<br>T 12 | A 15<br>K 11<br>V 11 | L 17 | A 16<br>K 11 | | G 14<br>L 13<br>K 11<br>E 11 |
| L 12 | L 23<br>I 13<br>A 11 | E 12<br>I 11<br>K 10 | K 18<br>E 12<br>L 12 | A 27<br>L 17<br>V 14<br>T 12 | A 14<br>K 12 | L 15<br>K 11 | | | G 14<br>K 12<br>L 10 | 600 | L 12 | L 24<br>I 12 | E 12<br>I 11 | K 17<br>E 13<br>L 12 | A 28<br>L 16<br>V 15<br>T 12 | A 14<br>K 12<br>V 10 | L 17 | | | G 14<br>L 12<br>K 11<br>E 11<br>T 10 |
| L | L,I,A | E,I,K | K,E,L | A,L,V,T | A,K | L,K | | | G,K,L | | | | | | | | | | | |
| | | | | | | | | | | 1200 | L 12 | L 23<br>I 13<br>A 10 | E 12<br>K 11<br>I 11 | K 17<br>E 12<br>L 12 | A 27<br>L 17<br>V 14<br>T 12 | A 14<br>K 12 | L 15<br>K 11 | | | G 13<br>K 12<br>L 11 |
| | | | | | | | | | | 2400 | L 12 | L 23<br>I 13<br>A 10 | E 12<br>I 12<br>K 11 | K 16<br>E 12<br>L 12 | A 27<br>L 17<br>V 14<br>T 13 | A 13<br>K 12 | L 15<br>K 11 | | | G 13<br>K 12<br>L 11 |
| | | | | | | | | | | 3600 | L 12<br>A 10 | L 24<br>I 17<br>A 10 | I 12<br>E 12<br>K 10 | K 16<br>E 12<br>L 12 | A 27<br>L 16<br>V 14<br>T 12 | A 14<br>K 12 | L 15<br>K 11 | | | G 13<br>K 11<br>L 11 |
| | | | | | | | | | | | L | L,I,A | E,I,K | K,E,L | A,L,V,T | A,K | L,K | | | G,K,L |

**SI Fig. 3. Consensus sequences derived from relative amino acid distribution at sites P1-P5 and P1'-P5' after various times of incubation.** Left, denatured ANXA1; right, folded ANXA1 were used as substrates. Numbers were taken from SI Fig. 2 and represent % of occurrence of the residue indicated at the given position. Only residues found at >10% are shown. Red letters indicate that consensus sequences given at the bottom of a cell are reached. Empty fields, no protease selectivity toward specific amino acids at this position.

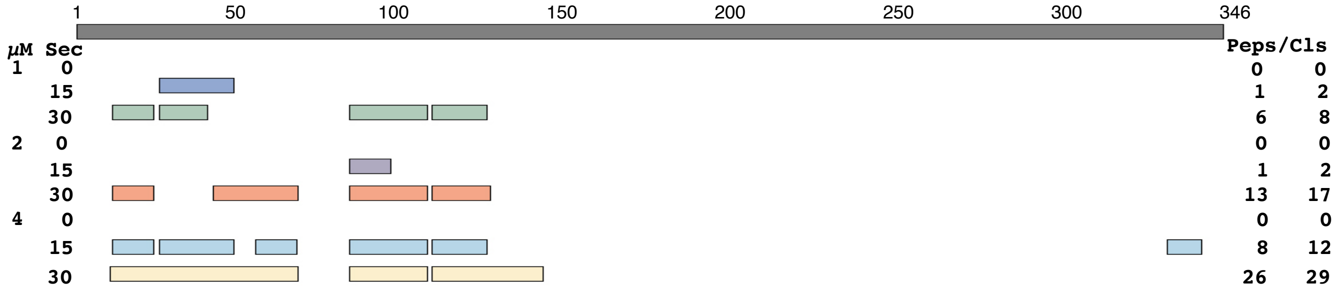

| | | | | | | | | | | | | | | | | | | | | | | | | | | | | | | | | | | | | | | | | | | | | | | | | | | | |
|---|---|---|---|---|---|---|---|---|---|---|---|---|---|---|---|---|---|---|---|---|---|---|---|---|---|---|---|---|---|---|---|---|---|---|---|---|---|---|---|---|---|---|---|---|---|---|---|---|---|---|---|
| | 1 | 2 | 3 | 4 | 5 | 6 | 7 | 8 | 9 | 10 | 11 | 12 | 13 | 14 | 15 | 16 | 17 | 18 | 19 | 20 | 21 | 22 | 23 | 24 | 25 | 26 | 27 | 28 | 29 | 30 | 31 | 32 | 33 | 34 | 35 | 36 | 37 | 38 | 39 | 40 | 41 | 42 | 43 | 44 | 45 | 46 | 47 | 48 | 49 | 50 | Residue no. |
| | A | M | V | S | E | F | L | K | Q | A | W | F | I | E | N | E | E | Q | E | Y | V | Q | T | V | K | S | S | K | G | G | P | G | S | A | V | S | P | Y | P | T | F | N | P | S | S | D | V | A | A | | Sequence |
| | | Helix 1 | | | | | | | | | | | | | | | | Helix 2 | | | | | | | | | | | LA | | | | | | | | | | | | | | Helix 3 | | | | | | | | 2ndary struct. |
| | | | | | | | | | | | | | | | | | | | | | | | | | | | | | | | | | | | | | | | | | | | | | | | | | | | Surf./buried |
| | | | | | | | | | | | | | | | | | | | | | | | | | | | | | | | | | | | | | | | | | | | | | | | | | | | B-factors |
| | : | : | * | * | | * | : | : | * | * | | : | : | : | : | | * | * | : | | : | : | | . | : | | | | | | * | | | | | | | | | | * | : | | * | : | * | * | | * | | Conservation |
| 15 sec, 1 µM HTRA1 | | | | | | | | | | 0 | 0 | | | | | | | | | | | | | 0 | 1 | | | 0 | | | | | | | | | | | | | 0 | | | | | | | | 1 | | Rel. freq. cuts |
| 30 sec, 1 µM HTRA1 | | | | | | | | | | 0 | 1 | | | | | | | | | | | | | 0 | 2 | | | 0 | | | | | | | | | | | | | 1 | | | | | | | | 0 | | |
| 15 sec, 2 µM HTRA1 | | | | | | | | | | 0 | 0 | | | | | | | | | | | | | 0 | 0 | | | 0 | | | | | | | | | | | | | 0 | | | | | | | | 0 | | |
| 30 sec, 2 µM HTRA1 | | | | | | | | | | 0 | 2 | | | | | | | | | | | | | 0 | 2 | | | 0 | | | | | | | | | | | | | 1 | | | | | | | | 0 | | |
| 15 sec, 4 µM HTRA1 | | | | | | | | | | 0 | 1 | | | | | | | | | | | | | 0 | 1 | | | 0 | | | | | | | | | | | | | 0 | | | | | | | | 1 | | |
| 30 sec, 4 µM HTRA1 | | | | | | | | | | 1 | 2 | | | | | | | | | | | | | 1 | 6 | | | 1 | | | | | | | | | | | | | 8 | | | | | | | | 1 | | |

| | | | | | | | | | | | | | | | | | | | | | | | | | | | | | | | | | | | | | | | | | | | | | | | | | | | |
|---|---|---|---|---|---|---|---|---|---|---|---|---|---|---|---|---|---|---|---|---|---|---|---|---|---|---|---|---|---|---|---|---|---|---|---|---|---|---|---|---|---|---|---|---|---|---|---|---|---|---|---|
| | 51 | 52 | 53 | 54 | 55 | 56 | 57 | 58 | 59 | 60 | 61 | 62 | 63 | 64 | 65 | 66 | 67 | 68 | 69 | 70 | 71 | 72 | 73 | 74 | 75 | 76 | 77 | 78 | 79 | 80 | 81 | 82 | 83 | 84 | 85 | 86 | 87 | 88 | 89 | 90 | 91 | 92 | 93 | 94 | 95 | 96 | 97 | 98 | 99 | 100 | Residue no. |
| | H | K | A | I | M | V | K | G | V | D | E | A | T | I | I | D | I | L | T | K | R | N | N | A | Q | R | Q | Q | I | K | A | A | Y | L | Q | E | T | G | K | P | L | D | E | T | L | K | K | A | L | | Sequence |
| | Helix 3 | | | | | | LB | | | | Helix 4 | | | | | | | | | | | LC | Helix 5 | | | | | | | | | | | | | | | | | LD | Helix 6 | | | | | | | | | | 2ndary struct. |
| | | | | | | | | | | | | | | | | | | | | | | | | | | | | | | | | | | | | | | | | | | | | | | | | | | | Surf./buried |
| | | | | | | | | | | | | | | | | | | | | | | | | | | | | | | | | | | | | | | | | | | | | | | | | | | | B-factors |
| | . | : | . | : | | . | : | * | * | * | * | * | * | * | : | : | : | | * | | * | | * | * | * | * | | : | : | * | * | . | * | | : | | . | * | : | | * | : | * | . | : | : | : | . | * | | Conservation |
| 15 sec, 1 µM HTRA1 | | | | 0 | | | 0 | | | | | | | 0 | | | | 0 | | | | | | | | | | | | | | 0 | | | 0 | | | | | | | | 0 | 0 | 0 | 0 | 0 | | 0 | 0 | Rel. freq. cuts |
| 30 sec, 1 µM HTRA1 | | | | 0 | | | 0 | | | | | | | 0 | | | | 0 | | | | | | | | | | | | | | 2 | | | 0 | | | | | | | | 0 | 0 | 0 | 1 | 0 | | 0 | 0 | |
| 15 sec, 2 µM HTRA1 | | | | 0 | | | 0 | | | | | | | 0 | | | | 0 | | | | | | | | | | | | | | 1 | | | 0 | | | | | | | | 0 | 0 | 0 | 1 | 0 | | 0 | 0 | |
| 30 sec, 2 µM HTRA1 | | | | 1 | | | 0 | | | | | | | 1 | | | | 1 | | | | | | | | | | | | | | 9 | | | 1 | | | | | | | | 0 | 1 | 0 | 1 | 1 | | 1 | 0 | |
| 15 sec, 4 µM HTRA1 | | | | 1 | | | 0 | | | | | | | 0 | | | | 1 | | | | | | | | | | | | | | 6 | | | 0 | | | | | | | | 0 | 0 | 0 | 1 | 0 | | 0 | 0 | |
| 30 sec, 4 µM HTRA1 | | | | 2 | | | 1 | | | | | | | 2 | | | | 1 | | | | | | | | | | | | | | 16 | | | 1 | | | | | | | | 1 | 4 | 1 | 1 | 1 | | 1 | 1 | |

| | | | | | | | | | | | | | | | | | | | | | | | | | | | | | | | | | | | | | | | | | | | | | | | | | | | |
|---|---|---|---|---|---|---|---|---|---|---|---|---|---|---|---|---|---|---|---|---|---|---|---|---|---|---|---|---|---|---|---|---|---|---|---|---|---|---|---|---|---|---|---|---|---|---|---|---|---|---|---|
| | 101 | 102 | 103 | 104 | 105 | 106 | 107 | 108 | 109 | 110 | 111 | 112 | 113 | 114 | 115 | 116 | 117 | 118 | 119 | 120 | 121 | 122 | 123 | 124 | 125 | 126 | 127 | 128 | 129 | 130 | 131 | 132 | 133 | 134 | 135 | 136 | 137 | 138 | 139 | 140 | 141 | 142 | 143 | 144 | 145 | 146 | 147 | 148 | 149 | 150 | Residue no. |
| | G | H | L | E | E | V | V | L | A | L | L | K | T | P | A | Q | F | D | A | D | E | L | R | A | A | M | K | G | L | G | T | D | E | D | T | L | I | E | I | L | A | S | R | T | N | K | E | I | R | | Sequence |
| | Helix 7 | | | | | | | | | | | | | Helix 8 | | | | | | | | | | | | | | | LE | | | | Helix 9 | | | | | | | | | | | | Helix 10 | | | | | | 2ndary struct. |
| | | | | | | | | | | | | | | | | | | | | | | | | | | | | | | | | | | | | | | | | | | | | | | | | | | | Surf./buried |
| | | | | | | | | | | | | | | | | | | | | | | | | | | | | | | | | | | | | | | | | | | | | | | | | | | | B-factors |
| | . | * | : | : | : | : | | : | * | : | : | * | * | * | * | : | : | * | . | : | * | * | * | . | . | * | * | . | | * | * | * | * | : | * | * | | * | * | * | | : | * | . | * | . | : | * | : | | Conservation |
| 15 sec, 1 µM HTRA1 | | 0 | | | | | 0 | | | | | | | | | | | | | | | | 0 | | 0 | 0 | | | | | | | | | | | | | | | 0 | 0 | | | | | | | | | Rel. freq. cuts |
| 30 sec, 1 µM HTRA1 | | 0 | | | | | 3 | | | | | | | | | | | | | | | | 1 | | 1 | 0 | | | | | | | | | | | | | | | 0 | 0 | | | | | | | | | |
| 15 sec, 2 µM HTRA1 | | 0 | | | | | 0 | | | | | | | | | | | | | | | | 0 | | 0 | 0 | | | | | | | | | | | | | | | 0 | 0 | | | | | | | | | |
| 30 sec, 2 µM HTRA1 | | 1 | | | | | 15 | | | | | | | | | | | | | | | | 4 | | 4 | 1 | | | | | | | | | | | | | | | 0 | 0 | | | | | | | | | |
| 15 sec, 4 µM HTRA1 | | 1 | | | | | 5 | | | | | | | | | | | | | | | | 0 | | 1 | 0 | | | | | | | | | | | | | | | 0 | 0 | | | | | | | | | |
| 30 sec, 4 µM HTRA1 | | 2 | | | | | 17 | | | | | | | | | | | | | | | | 4 | | 6 | 3 | | | | | | | | | | | | | | | 1 | 1 | | | | | | | | | |

| | | | | | | | | | | | | | | | | | | | | | | | | | | | | | | | | | | | | | | | | | | | | | | | | | | | |
|---|---|---|---|---|---|---|---|---|---|---|---|---|---|---|---|---|---|---|---|---|---|---|---|---|---|---|---|---|---|---|---|---|---|---|---|---|---|---|---|---|---|---|---|---|---|---|---|---|---|---|---|
| | 151 | 152 | 153 | 154 | 155 | 156 | 157 | 158 | 159 | 160 | 161 | 162 | 163 | 164 | 165 | 166 | 167 | 168 | 169 | 170 | 171 | 172 | 173 | 174 | 175 | 176 | 177 | 178 | 179 | 180 | 181 | 182 | 183 | 184 | 185 | 186 | 187 | 188 | 189 | 190 | 191 | 192 | 193 | 194 | 195 | 196 | 197 | 198 | 199 | 200 | Residue no. |
| | I | N | R | V | Y | R | E | E | L | K | R | D | L | A | K | D | I | T | S | D | T | S | G | D | F | R | N | A | L | L | S | L | A | K | G | D | R | S | E | D | F | G | V | N | E | D | L | A | D | | Sequence |
| | Helix 10 | | | | | | | | | LF | | | Helix 11 | | | | | | | | | | Helix 12 | | | | | | | | | | | | | LG | | | | | | | | | Helix 13 | | | | | | 2ndary struct. |
| | | | | | | | | | | | | | | | | | | | | | | | | | | | | | | | | | | | | | | | | | | | | | | | | | | | Surf./buried |
| | | | | | | | | | | | | | | | | | | | | | | | | | | | | | | | | | | | | | | | | | | | | | | | | | | | B-factors |
| | | | | | * | : | : | | * | * | : | * | : | : | : | | | | * | * | * | * | * | . | : | . | . | * | * | : | | * | * | * | . | : | * | . | : | : | | | : | * | : | : | : | * | : | | Conservation |
| 15 sec, 1 µM HTRA1 | | | | | | | | | | | | | | | | | | | | | | | | | | | | | | | | | | | | | | | | | | | | | | | | | | | Rel. freq. cuts |
| 30 sec, 1 µM HTRA1 | | | | | | | | | | | | | | | | | | | | | | | | | | | | | | | | | | | | | | | | | | | | | | | | | | | |
| 15 sec, 2 µM HTRA1 | | | | | | | | | | | | | | | | | | | | | | | | | | | | | | | | | | | | | | | | | | | | | | | | | | | |
| 30 sec, 2 µM HTRA1 | | | | | | | | | | | | | | | | | | | | | | | | | | | | | | | | | | | | | | | | | | | | | | | | | | | |
| 15 sec, 4 µM HTRA1 | | | | | | | | | | | | | | | | | | | | | | | | | | | | | | | | | | | | | | | | | | | | | | | | | | | |
| 30 sec, 4 µM HTRA1 | | | | | | | | | | | | | | | | | | | | | | | | | | | | | | | | | | | | | | | | | | | | | | | | | | | |

| | | | | | | | | | | | | | | | | | | | | | | | | | | | | | | | | | | | | | | | | | | | | | | | | | | | |
|---|---|---|---|---|---|---|---|---|---|---|---|---|---|---|---|---|---|---|---|---|---|---|---|---|---|---|---|---|---|---|---|---|---|---|---|---|---|---|---|---|---|---|---|---|---|---|---|---|---|---|---|
| | 201 | 202 | 203 | 204 | 205 | 206 | 207 | 208 | 209 | 210 | 211 | 212 | 213 | 214 | 215 | 216 | 217 | 218 | 219 | 220 | 221 | 222 | 223 | 224 | 225 | 226 | 227 | 228 | 229 | 230 | 231 | 232 | 233 | 234 | 235 | 236 | 237 | 238 | 239 | 240 | 241 | 242 | 243 | 244 | 245 | 246 | 247 | 248 | 249 | 250 | Residue no. |
| | D | A | R | A | L | Y | E | A | G | E | R | R | K | G | T | D | V | N | V | F | N | T | I | L | T | T | R | S | Y | P | Q | L | R | R | V | F | Q | K | Y | T | K | Y | S | K | H | D | M | N | K | | Sequence |
| | Helix 13 | | | | | | | | | | LH | | | | | | Helix 14 | | | | | | | | | | | | Helix 15 | | | | | | | | | | | | | | LI | | | | | | | | 2ndary struct. |
| | | | | | | | | | | | | | | | | | | | | | | | | | | | | | | | | | | | | | | | | | | | | | | | | | | | Surf./buried |
| | | | | | | | | | | | | | | | | | | | | | | | | | | | | | | | | | | | | | | | | | | | | | | | | | | | B-factors |
| | * | * | * | * | * | * | : | * | . | * | : | : | : | * | : | * | : | * | * | * | | * | : | * | * | : | : | . | : | | : | | | . | | * | * | | * | | * | * | : | : | . | * | * | : | * | | Conservation |
| 15 sec, 1 µM HTRA1 | | | | | | | | | | | | | | | | | | | | | | | | | | | | | | | | | | | | | | | | | | | | | | | | | | | Rel. freq. cuts |
| 30 sec, 1 µM HTRA1 | | | | | | | | | | | | | | | | | | | | | | | | | | | | | | | | | | | | | | | | | | | | | | | | | | | |
| 15 sec, 2 µM HTRA1 | | | | | | | | | | | | | | | | | | | | | | | | | | | | | | | | | | | | | | | | | | | | | | | | | | | |
| 30 sec, 2 µM HTRA1 | | | | | | | | | | | | | | | | | | | | | | | | | | | | | | | | | | | | | | | | | | | | | | | | | | | |
| 15 sec, 4 µM HTRA1 | | | | | | | | | | | | | | | | | | | | | | | | | | | | | | | | | | | | | | | | | | | | | | | | | | | |
| 30 sec, 4 µM HTRA1 | | | | | | | | | | | | | | | | | | | | | | | | | | | | | | | | | | | | | | | | | | | | | | | | | | | |

| | | | | | | | | | | | | | | | | | | | | | | | | | | | | | | | | | | | | | | | | | | | | | | | | | | | |
|---|---|---|---|---|---|---|---|---|---|---|---|---|---|---|---|---|---|---|---|---|---|---|---|---|---|---|---|---|---|---|---|---|---|---|---|---|---|---|---|---|---|---|---|---|---|---|---|---|---|---|---|
| | 251 | 252 | 253 | 254 | 255 | 256 | 257 | 258 | 259 | 260 | 261 | 262 | 263 | 264 | 265 | 266 | 267 | 268 | 269 | 270 | 271 | 272 | 273 | 274 | 275 | 276 | 277 | 278 | 279 | 280 | 281 | 282 | 283 | 284 | 285 | 286 | 287 | 288 | 289 | 290 | 291 | 292 | 293 | 294 | 295 | 296 | 297 | 298 | 299 | 300 | Residue no. |
| | L | D | L | E | L | K | G | D | I | E | K | C | L | T | A | I | V | K | C | A | T | S | K | P | A | F | F | A | E | K | L | H | Q | A | M | K | G | V | G | T | R | H | K | A | L | I | R | I | M | | Sequence |
| | LI | | | | | | Helix 16 | | | | | | | | | | | | | | | | | Helix 17 | | | | | | | | | | | | | | LJ | | | | Helix 18 | | | | | | | | | 2ndary struct. |
| | | | | | | | | | | | | | | | | | | | | | | | | | | | | | | | | | | | | | | | | | | | | | | | | | | | Surf./buried |
| | | | | | | | | | | | | | | | | | | | | | | | | | | | | | | | | | | | | | | | | | | | | | | | | | | | B-factors |
| | : | * | : | * | : | * | * | : | : | * | : | * | : | * | . | : | . | : | * | . | * | . | | * | | : | * | * | * | * | * | : | | * | * | * | * | | * | * | : | . | : | | * | * | * | * | * | | Conservation |
| 15 sec, 1 µM HTRA1 | | | | | | | | | | | | | | | | | | | | | | | | | | | | | | | | | | | | | | | | | | | | | | | | | | | Rel. freq. cuts |
| 30 sec, 1 µM HTRA1 | | | | | | | | | | | | | | | | | | | | | | | | | | | | | | | | | | | | | | | | | | | | | | | | | | | |
| 15 sec, 2 µM HTRA1 | | | | | | | | | | | | | | | | | | | | | | | | | | | | | | | | | | | | | | | | | | | | | | | | | | | |
| 30 sec, 2 µM HTRA1 | | | | | | | | | | | | | | | | | | | | | | | | | | | | | | | | | | | | | | | | | | | | | | | | | | | |
| 15 sec, 4 µM HTRA1 | | | | | | | | | | | | | | | | | | | | | | | | | | | | | | | | | | | | | | | | | | | | | | | | | | | |
| 30 sec, 4 µM HTRA1 | | | | | | | | | | | | | | | | | | | | | | | | | | | | | | | | | | | | | | | | | | | | | | | | | | | |

| | | | | | | | | | | | | | | | | | | | | | | | | | | | | | | | | | | | | | | | | | | | | | | | | | | | |
|---|---|---|---|---|---|---|---|---|---|---|---|---|---|---|---|---|---|---|---|---|---|---|---|---|---|---|---|---|---|---|---|---|---|---|---|---|---|---|---|---|---|---|---|---|---|---|---|---|---|---|---|
| | 301 | 302 | 303 | 304 | 305 | 306 | 307 | 308 | 309 | 310 | 311 | 312 | 313 | 314 | 315 | 316 | 317 | 318 | 319 | 320 | 321 | 322 | 323 | 324 | 325 | 326 | 327 | 328 | 329 | 330 | 331 | 332 | 333 | 334 | 335 | 336 | 337 | 338 | 339 | 340 | 341 | 342 | 343 | 344 | 345 | 346 | | | | | Residue no. |
| | S | R | S | E | I | D | M | N | D | I | K | A | F | Y | Q | K | M | Y | G | I | S | L | C | Q | A | I | L | D | E | T | K | G | D | Y | E | K | I | L | V | A | L | C | G | G | N | | | | | | Sequence |
| | Helix 18 | | | LK | | | Helix 19 | | | | | | | | | | | | | | LL | Helix 20 | | | | | | | | | | Helix 21 | | | | | | | | | | | | | | | | | | | 2ndary struct. |
| | | | | | | | | | | | | | | | | | | | | | | | | | | | | | | | | | | | | | | | | | | | | | | | | | | | Surf./buried |
| | | | | | | | | | | | | | | | | | | | | | | | | | | | | | | | | | | | | | | | | | | | | | | | | | | | B-factors |
| | * | * | | * | : | * | | : | : | * | * | | | * | : | * | | * | * | : | | * | * | * | * | * | : | * | : | | * | * | | * | * | . | * | * | * | * | * | | * | | : | | | | | | Conservation |
| 15 sec, 1 µM HTRA1 | | | | | | | | | | | | | | | | | | | | | | | | | | | | 0 | | | | | | | | | | | | 0 | | | | | | | | | | | Rel. freq. cuts |
| 30 sec, 1 µM HTRA1 | | | | | | | | | | | | | | | | | | | | | | | | | | | | 0 | | | | | | | | | | | | 0 | | | | | | | | | | | |
| 15 sec, 2 µM HTRA1 | | | | | | | | | | | | | | | | | | | | | | | | | | | | 0 | | | | | | | | | | | | 0 | | | | | | | | | | | |
| 30 sec, 2 µM HTRA1 | | | | | | | | | | | | | | | | | | | | | | | | | | | | 0 | | | | | | | | | | | | 0 | | | | | | | | | | | |
| 15 sec, 4 µM HTRA1 | | | | | | | | | | | | | | | | | | | | | | | | | | | | 1 | | | | | | | | | | | | 1 | | | | | | | | | | | |
| 30 sec, 4 µM HTRA1 | | | | | | | | | | | | | | | | | | | | | | | | | | | | 0 | | | | | | | | | | | | 0 | | | | | | | | | | | |

**SI Fig. 4. Proteolysis of folded ANXA1 at low HTRA1 concentrations.** 20 µM ANXA1 was incubated with 1, 2 or 4 µM HTRA1 at 37°C. Samples were taken at the timepoint indicated for MS analysis. Upper part, top bar: linear representation of the entire substrate protein. peptide sequences that align without gaps are grouped into so-called fragments shown as bars. The number of peptides identified (Peps) and the total number of cleavage sites (Cls) are given at the right. The identified peptides aligned to the primary amino acid sequence of ANXA1 are provided in Supplementary data 4. Lower part, P1 residues identified in n=4 independent experiments significantly enriched compared to controls. Structural elements i.e. helices and loops (2ndary struct.), B-factors and surface accessibility (surf. (grey)/buried (black) as detected in the crystal structure of folded ANXA1 (pdb:1HM6) are indicated. The colour code for B-factors indicates gradually structural rigidity (blue) to flexibility (red). Amino acid sequence conservation (Conservation) is derived from a multiple sequence alignment (Supplementary data 2). * = identical, : = conserved residues. Numbers below individual P1 positions indicate the relative frequency of cuts (Rel. freq. cuts) at each time point. No number indicates that no cleavage was detected at any of the time points investigated.

A

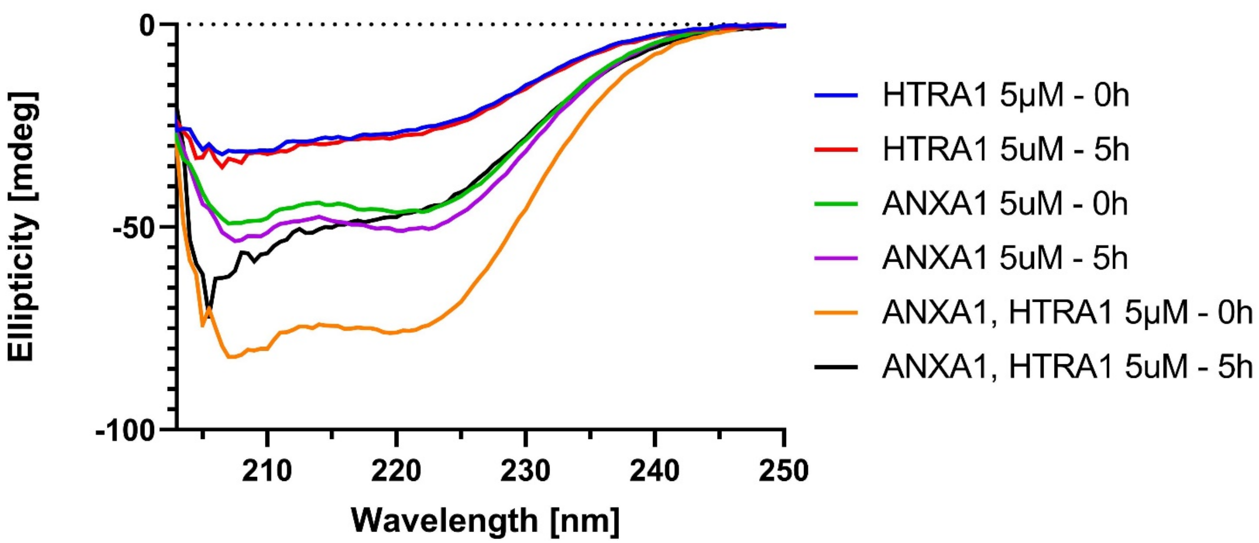

B

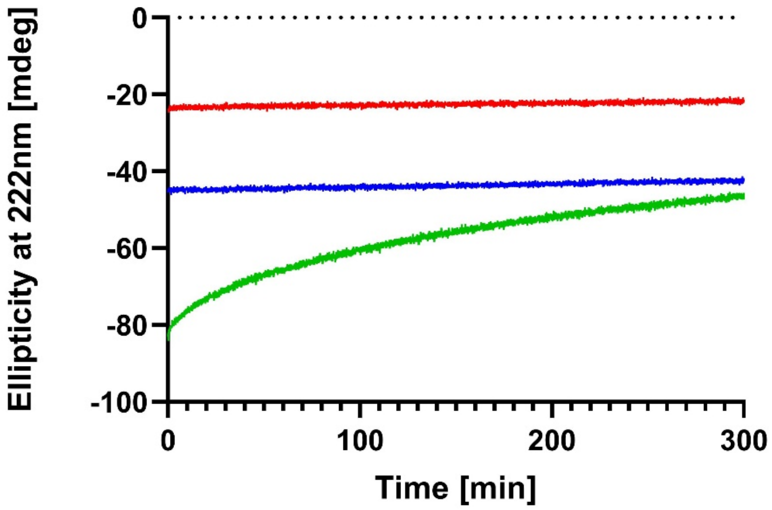

**SI Fig. 5. Circular dichroism spectroscopy.** A. CD spectra of HTRA1, ANXA1 or a mix of the two proteins at the times indicated. The large loss of the $\alpha$-helical minimum at 222 nm when ANXA1 has been proteolytically digested for 5 hours by HTRA1 can be observed. For each spectrum the photomultiplier HT voltage was less than 600 V throughout the scan. B. A time course showing the gain in ellipticity at 222 nm over 5 hours as 5 $\mu$M ANXA1 is proteolytically digested after mixing with 5 $\mu$M HTRA1 (*green*). 5 $\mu$M ANXA1 alone (*blue*) or 5 $\mu$M HTRA1 alone (*red*) control experiments are also shown and show no change.

**SI Table 1. List of P1 residues with the second highest relative numbers of cuts**

| Residue | Denatured 600 sec | Folded 600 sec | Folded 3600 sec |
|---|---|---|---|
| A 2 | | | x |
| S 5 | | | x |
| Q 10 | | x | |
| T 24 | x | x | x |
| S 27 | | x | x |
| M 56 | | | x |
| V 57 | x | | |
| L 69 | | | |
| T 70 | | | x |
| N 73 | | x | x |
| A 83 | x | x | |
| L 85 | x | x | x |
| T 88 | | x | x |
| T 95 | x | x | |
| A 99 | | | x |
| L109 | x | | x |
| L112 | | x | x |
| A116 | | | x |
| A126 | | x | |
| M127 | | | x |
| I140 | | | x |
| L141 | x | | x |
| T145 | x | | x |
| L160 | x | | x |
| T169 | x | | |
| L181 | | x | x |
| A184 | x | | |
| A205 | x | | |
| T223 | x | | |
| V236 | x | x | |
| T241 | | | x |
| L254 | | | x |
| C263 | x | | |
| T265 | | | x |
| C270 | x | | |
| A271 | x | | |
| A276 | | | |
| V289 | x | | x |
| I299 | x | | x |
| V301 | x | | x |
| S302 | x | | x |
| A313 | x | x | x |
| M318 | x | | |
| C324 | | x | x |
| A326 | x | | x |
| V340 | | x | x |
| C343 | | | x |

Class II P1 residues (11 - 20 cuts) detected in proteolysis experiments with denatured and folded ANXA1 at the timepoints indicated. x = detected

Table 1: P1 sites that are cleaved more efficiently in denatured vs folded ANXA1

| Res | 0 | 15 | 30 | 60 | 90 | 120 | 300 | 600 | 1200 | 2400 | 3600 |
|---|---|---|---|---|---|---|---|---|---|---|---|
| L69 | 0 | 0 | 0 | 2 | 4 | 8 | 16 | 21 | | | |
| L69 | 0 | 0 | 0 | 1 | 3 | 4 | 11 | 15 | 22 | 24 | 27 |
| T70 | 0 | 0 | 0 | 1 | 1 | 2 | 5 | 11 | | | |
| T70 | 0 | 0 | 0 | 1 | 3 | 4 | 5 | 5 | 6 | 9 | 12 |
| N73 | 0 | 0 | 0 | 2 | 6 | 7 | 10 | 5 | | | |
| N73 | 0 | 0 | 1 | 1 | 1 | 5 | 9 | 12 | 13 | 11 | 14 |
| T101 | 0 | 0 | 0 | 1 | 1 | 4 | 9 | 9 | | | |
| T101 | 0 | 0 | 0 | 1 | 1 | 2 | 4 | 4 | 4 | 4 | 4 |
| I140 | 1 | 1 | 2 | 7 | 6 | 8 | 9 | 10 | | | |
| I140 | 0 | 0 | 0 | 1 | 3 | 3 | 5 | 8 | 10 | 13 | 10 |
| L141 | 0 | 0 | 1 | 3 | 6 | 8 | 16 | 19 | | | |
| L141 | 0 | 0 | 0 | 0 | 1 | 2 | 5 | 9 | 13 | 15 | 15 |
| I152 | 0 | 0 | 0 | 0 | 1 | 3 | 6 | 4 | | | |
| I152 | 0 | 0 | 0 | 0 | 0 | 0 | 0 | 2 | 2 | 3 | 2 |
| V155 | 0 | 0 | 0 | 0 | 0 | 0 | 3 | 4 | | | |
| V155 | 0 | 0 | 0 | 0 | 0 | 0 | 0 | 1 | 3 | 3 | 5 |
| L160 | 0 | 0 | 1 | 4 | 3 | 7 | 12 | 14 | | | |
| L160 | 0 | 0 | 0 | 0 | 0 | 0 | 3 | 6 | 8 | 8 | 11 |
| L164 | 0 | 1 | 1 | 1 | 2 | 2 | 2 | 3 | | | |
| L164 | 0 | 0 | 0 | 0 | 0 | 0 | 0 | 0 | 0 | 0 | 1 |
| A165 | 0 | 0 | 0 | 1 | 2 | 1 | 1 | 1 | | | |
| A165 | 0 | 0 | 0 | 0 | 0 | 0 | 1 | 1 | 1 | 1 | 1 |
| I168 | 0 | 0 | 0 | 1 | 1 | 2 | 3 | 2 | | | |
| I168 | 0 | 0 | 0 | 0 | 0 | 0 | 2 | 4 | 4 | 4 | 3 |
| T169 | 0 | 0 | 0 | 2 | 2 | 7 | 9 | 12 | | | |
| T169 | 0 | 0 | 0 | 0 | 0 | 1 | 3 | 6 | 8 | 9 | 9 |
| T172 | 0 | 0 | 0 | 1 | 0 | 2 | 5 | 3 | | | |
| T172 | 0 | 0 | 0 | 0 | 0 | 0 | 1 | 2 | 5 | 5 | 7 |
| A179 | 0 | 0 | 0 | 0 | 0 | 0 | 2 | 4 | | | |
| A179 | 0 | 0 | 0 | 0 | 0 | 0 | 0 | 2 | 2 | 2 | 3 |
| L181 | 0 | 1 | 3 | 11 | 14 | 19 | 24 | 27 | | | |
| L181 | 0 | 0 | 0 | 2 | 5 | 7 | 12 | 15 | 17 | 18 | 18 |

| | | | | | | | | | | |
|---|---|---|---|---|---|---|---|---|---|---|
| S182 | 0 | 0 | 0 | 0 | 0 | 0 | 1 | 3 | | | |
| S182 | 0 | 0 | 0 | 0 | 0 | 0 | 0 | 0 | 0 | 1 | 2 |
| A184 | 0 | 0 | 0 | 2 | 1 | 7 | 10 | 7 | | | |
| A184 | 0 | 0 | 0 | 0 | 0 | 0 | 3 | 7 | 9 | 8 | 8 |
| A205 | 0 | 0 | 2 | 8 | 6 | 9 | 11 | 13 | | | |
| A205 | 0 | 0 | 0 | 0 | 1 | 2 | 4 | 6 | 7 | 8 | 10 |
| A209 | 0 | 0 | 1 | 2 | 4 | 6 | 5 | 7 | | | |
| A209 | 0 | 0 | 0 | 0 | 0 | 0 | 1 | 2 | 3 | 6 | 7 |
| V218 | 0 | 0 | 1 | 4 | 3 | 4 | 4 | 1 | | | |
| V218 | 0 | 0 | 0 | 0 | 0 | 0 | 1 | 2 | 3 | 4 | 4 |
| N219 | 0 | 0 | 1 | 1 | 1 | 1 | 1 | 1 | | | |
| N219 | 0 | 0 | 0 | 0 | 0 | 1 | 2 | 2 | 3 | 3 | 3 |
| T223 | 0 | 1 | 2 | 6 | 6 | 8 | 9 | 11 | | | |
| T223 | 0 | 0 | 0 | 1 | 2 | 2 | 4 | 5 | 7 | 9 | 10 |
| T226 | 0 | 0 | 2 | 3 | 4 | 7 | 6 | 2 | | | |
| T226 | 0 | 0 | 0 | 0 | 1 | 1 | 3 | 6 | 7 | 7 | 8 |
| V236 | 0 | 1 | 6 | 12 | 13 | 15 | 17 | 15 | | | |
| V236 | 0 | 0 | 0 | 1 | 5 | 6 | 11 | 15 | 18 | 18 | 21 |
| Q238 | 0 | 0 | 0 | 0 | 1 | 5 | 7 | 5 | | | |
| Q238 | 0 | 0 | 0 | 0 | 0 | 0 | 0 | 1 | 2 | 1 | 3 |
| T241 | 0 | 0 | 0 | 0 | 1 | 1 | 4 | 6 | | | |
| T241 | 0 | 0 | 0 | 0 | 0 | 0 | 0 | 1 | 4 | 10 | 13 |
| S244 | 0 | 0 | 0 | 0 | 0 | 0 | 1 | 2 | | | |
| S244 | 0 | 0 | 0 | 0 | 0 | 0 | 0 | 0 | 1 | 1 | 2 |
| V251 | 0 | 0 | 1 | 2 | 3 | 3 | 4 | 4 | | | |
| V251 | 0 | 0 | 0 | 0 | 0 | 1 | 2 | 2 | 2 | 2 | 2 |
| L252 | 0 | 0 | 0 | 1 | 1 | 1 | 2 | 3 | | | |
| L252 | 0 | 0 | 0 | 0 | 0 | 0 | 1 | 1 | 2 | 2 | 3 |
| L254 | 0 | 0 | 1 | 1 | 1 | 1 | 4 | 9 | | | |
| L254 | 0 | 0 | 0 | 0 | 1 | 1 | 1 | 3 | 5 | 8 | 11 |
| L256 | 0 | 0 | 0 | 1 | 1 | 1 | 3 | 5 | | | |
| L256 | 0 | 0 | 0 | 0 | 1 | 0 | 0 | 2 | 5 | 3 | 7 |
| C263 | 0 | 0 | 1 | 2 | 1 | 4 | 10 | 20 | | | |
| C263 | 0 | 0 | 0 | 0 | 0 | 0 | 1 | 1 | 6 | 6 | 9 |

| | | | | | | | | | | | |
|---|---|---|---|---|---|---|---|---|---|---|---|
| T265 | 0 | 0 | 0 | 0 | 1 | 4 | 6 | 8 | | | |
| T265 | 0 | 0 | 0 | 0 | 0 | 0 | 0 | 0 | 3 | 7 | 11 |
| I267 | 0 | 0 | 0 | 0 | 0 | 0 | 1 | 3 | | | |
| I267 | 0 | 0 | 0 | 0 | 0 | 0 | 0 | 0 | 1 | 1 | 5 |
| V268 | 0 | 0 | 0 | 0 | 0 | 0 | 1 | 1 | | | |
| V268 | 0 | 0 | 0 | 0 | 0 | 0 | 0 | 0 | 1 | 0 | 5 |
| C270 | 0 | 3 | 6 | 9 | 10 | 11 | 14 | 19 | | | |
| C270 | 0 | 0 | 1 | 4 | 5 | 8 | 15 | 22 | 26 | 25 | 33 |
| A271 | 0 | 1 | 2 | 3 | 3 | 7 | 9 | 12 | | | |
| A271 | 0 | 0 | 0 | 1 | 2 | 2 | 5 | 4 | 5 | 6 | 6 |
| A276 | 0 | 0 | 0 | 5 | 11 | 16 | 21 | 25 | | | |
| A276 | 0 | 0 | 0 | 0 | 1 | 5 | 11 | 14 | 20 | 23 | 28 |
| L282 | 0 | 2 | 3 | 7 | 10 | 13 | 20 | 25 | | | |
| L282 | 0 | 0 | 1 | 4 | 7 | 11 | 15 | 18 | 24 | 25 | 36 |
| Q284 | 0 | 1 | 1 | 3 | 4 | 5 | 8 | 8 | | | |
| Q284 | 0 | 0 | 0 | 0 | 1 | 1 | 3 | 4 | 5 | 5 | 5 |
| A285 | 0 | 1 | 1 | 3 | 4 | 6 | 8 | 10 | | | |
| A285 | 0 | 0 | 0 | 0 | 2 | 6 | 8 | 8 | 10 | 12 | 19 |
| K287 | 0 | 0 | 0 | 0 | 0 | 1 | 3 | 5 | | | |
| K287 | 0 | 0 | 0 | 0 | 0 | 0 | 0 | 1 | 1 | 2 | 2 |
| V289 | 0 | 1 | 2 | 6 | 10 | 11 | 11 | 11 | | | |
| V289 | 0 | 0 | 0 | 0 | 0 | 2 | 9 | 10 | 11 | 11 | 12 |
| T291 | 0 | 0 | 0 | 1 | 1 | 1 | 1 | 1 | | | |
| T291 | 0 | 0 | 0 | 0 | 0 | 0 | 1 | 4 | 5 | 5 | 6 |
| A295 | 0 | 0 | 0 | 0 | 1 | 1 | 1 | 2 | | | |
| A295 | 0 | 0 | 0 | 0 | 0 | 0 | 1 | 2 | 2 | 2 | 5 |
| L296 | 0 | 0 | 0 | 0 | 1 | 2 | 5 | 5 | | | |
| L296 | 0 | 0 | 0 | 0 | 0 | 0 | 1 | 2 | 4 | 4 | 3 |
| I299 | 0 | 0 | 0 | 4 | 7 | 9 | 13 | 15 | | | |
| I299 | 0 | 0 | 0 | 0 | 0 | 0 | 2 | 5 | 10 | 9 | 12 |
| M300 | 0 | 0 | 4 | 9 | 19 | 27 | 31 | 33 | | | |
| M300 | 0 | 0 | 0 | 0 | 3 | 5 | 20 | 26 | 31 | 31 | 35 |
| V301 | 1 | 1 | 1 | 3 | 7 | 8 | 12 | 19 | | | |
| V301 | 0 | 1 | 0 | 0 | 0 | 0 | 3 | 4 | 8 | 11 | 17 |

| | | | | | | | | | | | |
|---|---|---|---|---|---|---|---|---|---|---|---|
| S302 | 0 | 0 | 0 | 0 | 3 | 7 | 11 | 12 | | | |
| S302 | 0 | 0 | 0 | 0 | 0 | 0 | 1 | 3 | 5 | 7 | 11 |
| I311 | 0 | 0 | 0 | 0 | 1 | 2 | 3 | 3 | | | |
| I311 | 0 | 0 | 0 | 0 | 0 | 0 | 1 | 2 | 4 | 4 | 4 |
| A313 | 1 | 1 | 2 | 5 | 9 | 12 | 18 | 20 | | | |
| A313 | 0 | 0 | 1 | 0 | 2 | 2 | 12 | 16 | 20 | 22 | 24 |
| Q316 | 0 | 0 | 1 | 3 | 9 | 13 | 18 | 24 | | | |
| Q316 | 0 | 0 | 0 | 0 | 0 | 0 | 3 | 5 | 7 | 8 | 9 |
| M318 | 0 | 0 | 1 | 1 | 2 | 4 | 6 | 11 | | | |
| M318 | 0 | 0 | 0 | 0 | 1 | 0 | 4 | 6 | 9 | 9 | 11 |
| I321 | 0 | 0 | 0 | 0 | 2 | 5 | 6 | 6 | | | |
| I321 | 0 | 0 | 0 | 0 | 0 | 0 | 2 | 3 | 3 | 3 | 3 |
| C324 | 0 | 1 | 3 | 8 | 8 | 15 | 19 | 25 | | | |
| C324 | 0 | 1 | 0 | 1 | 2 | 4 | 11 | 14 | 20 | 23 | 24 |
| A326 | 0 | 2 | 6 | 11 | 14 | 16 | 16 | 22 | | | |
| A326 | 0 | 0 | 1 | 1 | 7 | 9 | 18 | 22 | 27 | 26 | 30 |
| I327 | 0 | 1 | 2 | 4 | 5 | 6 | 7 | 8 | | | |
| I327 | 0 | 0 | 0 | 0 | 0 | 1 | 6 | 5 | 7 | 8 | 8 |
| L328 | 0 | 0 | 1 | 2 | 4 | 3 | 4 | 4 | | | |
| L328 | 0 | 0 | 0 | 0 | 0 | 0 | 0 | 1 | 1 | 1 | 1 |
| T331 | 0 | 0 | 1 | 1 | 1 | 3 | 4 | 5 | | | |
| T331 | 0 | 0 | 0 | 0 | 0 | 0 | 2 | 3 | 5 | 5 | 6 |
| V340 | 0 | 0 | 3 | 7 | 10 | 9 | 11 | 11 | | | |
| V340 | 0 | 0 | 0 | 2 | 6 | 7 | 9 | 11 | 11 | 12 | 12 |
| A341 | 0 | 0 | 1 | 3 | 4 | 4 | 4 | 5 | | | |
| A341 | 0 | 0 | 0 | 0 | 0 | 1 | 2 | 4 | 5 | 5 | 5 |
| L342 | 0 | 0 | 1 | 3 | 4 | 4 | 3 | 4 | | | |
| L342 | 0 | 0 | 0 | 0 | 0 | 0 | 1 | 1 | 1 | 1 | 1 |
| C343 | 0 | 1 | 3 | 5 | 5 | 8 | 9 | 10 | | | |
| C343 | 0 | 0 | 0 | 0 | 1 | 3 | 6 | 6 | 9 | 10 | 11 |

Left column = P1 residues. Numbers in top row represent sec of incubation of ANXA1 with HTRA1; Numbers in all other lanes below represent the relative numbers of cuts at each P1 residue. For each P1 residue shown, the top and bottom lanes represent data obtained by digesting denatured or folded HTRA1, respectively. Res = amino acid residue and amino acid number in ANXA1.
